# Assessing the Reliability of LLM-Generated Phenotype-Genotype Associations Through External Validation

**DOI:** 10.64898/2026.08.13.744701

**Authors:** Caiwan Sun, Yi Xin, Sarah Zeng, Sudeep D. Sunthankar, Wu-Chen (Sander) Su, Jacob Lynn, Sergio Mundo, Mojgan Babanejad, Qiping Feng, Wei-Qi Wei

**Affiliations:** Department of Electrical and Computer Engineering, Vanderbilt University, Nashville, Tennessee, USA; Department of Computer Science, Vanderbilt University, Nashville, Tennessee, USA; Department of Medicine, Health, & Society, Vanderbilt University, Nashville, Tennessee, USA; Thomas P. Graham Jr. Division of Pediatric Cardiology and Center for Pediatric Precision Medicine, Department of Pediatrics, Monroe Carell Jr. Children’s Hospital and Vanderbilt University Medical Center, Nashville, Tennessee, USA; Division of Genetic Medicine and Clinical Pharmacology, Department of Medicine, Vanderbilt University Medical Center, Nashville, Tennessee, USA; Center for Precision Medicine, Department of Biomedical Informatics, Vanderbilt University Medical Center, Nashville, Tennessee, USA; Department of Biomedical Informatics, Vanderbilt University Medical Center, Nashville, Tennessee, USA

**Keywords:** large language model, clinical genomics, phenotype-genotype association, external validation, genomic knowledge base

## Abstract

**Background:** Phenotype-genotype associations underpin precision medicine by enabling disease prevention, early diagnosis, risk stratification, therapeutic target discovery, and personalized treatment. However, the rapid growth of scientific evidence has made manual curation of these associations increasingly labor-intensive, time-consuming, and incomplete. Large Language Models (LLMs) offer a potential path to scalable genomic generation and synthesis of this knowledge, but their ability to accurately identify phenotype-genotype associations and the extent to which these outputs are supported by established genomic knowledge bases remain unclear.

**Materials and Methods:** Four LLMs, Claude Sonnet 4.6, DeepSeek V4 Flash, Gemini 3 Flash Preview, and GPT-5.5, were benchmarked on six zero-shot task categories covering forward and reverse phenotype-gene and phenotype-SNP generation. A total of 4,196 associations were identified from curated inputs and evaluated through a multistage external verification pipeline comprising phenotype normalization, ontology mapping, genomic identifier validation against Ensembl, and evidence verification using both the GWAS Catalog and OMIM. Associations were assigned a fused evidence level of strong, moderate, weak, or none.

**Results:** Overall, 74.19% of generated associations were matched to at least one external genomic knowledge base; 9.15% received strong support and 54.46% moderate support. Phenotype-gene associations were more verifiable than phenotype-SNP associations (strong or moderate: 67.19% vs 54.06%). Among existing associations, Claude Sonnet 4.6 achieved the highest overall strong or moderate rate (69.2%), followed by GPT-5.5 (65.1%), DeepSeek V4 Flash (61.7%), and Gemini 3 Flash Preview (56.9%).

**Conclusion:** LLMs can support scalable generation of candidate phenotype-genotype associations. Performance varied substantially by relation type and was lower for SNP-level and rare disease associations, highlighting both the limitations of current genomic resources and the need for rigorous validation pipelines.

## INTRODUCTION

Phenotype-genotype associations link observable clinical features to their underlying genetic causes, and they contribute to the basis for genetic diagnosis, variant interpretation, and gene discovery [1]. In rare disease care, clinicians use these associations in both directions: starting from a patient’s phenotype to identify candidate genes, or starting from a variant identified by sequencing to assess whether its known genotype-phenotype associations align with the patient’s clinical presentation [2]. Genome sequencing now delivers molecular diagnoses across a broad spectrum of rare diseases and increasingly informs precision medicine [3]. To make this knowledge computable, phenotypes are recorded with standardized vocabularies such as the Human Phenotype Ontology (HPO) [4], and curated relationships between phenotypes, genes, and variants are maintained in resources such as OMIM, ClinVar, and ClinGen [2,5,6]. These associations are maintained mainly through manual expert curation, which is accurate but does not scale to the volume of genomic evidence now being produced [7]. Because curation requires reading and reconciling primary literature, newly published findings often take months or years to enter a clinically available database. Coverage is also uneven, and the same association can be interpreted differently across laboratories and resources [8]. Because genomic publications are growing faster than curation teams can process, the gap between published evidence and curated knowledge will continue to widen [7].

Large language models (LLMs) have recently been proposed to minimize this gap. They have advanced quickly across biomedical applications and show strong performance in clinical question answering, medical examination benchmarks, and biomedical natural language processing [9,10]. The performance is not uniform: LLMs remain task dependent and can fall behind locally trained models on certain clinical prediction tasks [11], and they struggle with zero-shot biomedical information extraction [12]. A more specific concern for genomics is hallucination, where a model produces a plausible but unsupported gene, variant, or association. Recent genomic benchmarks show that hallucination is still common in LLM-based genomic knowledge generation, across both gene-related and SNP-related tasks [13]. Because errors at this level can lead to pathogenicity misclassification and flawed clinical summaries, generated associations need traceable support from curated genomic resources rather than being trusted on their own [14,15]. Comparative evaluations of frontier models such as GPT, Gemini, Claude, and DeepSeek have become common in general medical settings [16-18], but systematic head-to-head comparisons in specialized genomics tasks remain scarce, especially for phenotype-gene and phenotype-SNP generation. Phenotype-driven gene generation has been studied with both free-text descriptions and HPO terms, and recent work suggests LLMs can be useful here [19,20]. However, their performance remains limited: in one benchmark, even the best-performing model, GPT-4, reached only 17% accuracy in top-50 gene prediction for rare disease diagnosis, behind dedicated bioinformatics tools [19,21]. Moving from phenotype-gene to phenotype-SNP generation introduces additional challenges. Genes are stable biological entities supported by well-curated symbol registries [22] whereas SNP identifiers are version sensitive, build dependent, and subject to deprecation across database releases. The evidence base for SNP-phenotype associations is also more fragmented. Common variant associations are indexed in the GWAS Catalog [23], while rare variant evidence is spread across resources such as ClinVar and ClinGen and across disease- or locus-specific reports [2,6].

Despite growing interest in LLMs for genomic applications, two gaps remain. First, most prior genomics studies evaluated only one or two models, and few have directly compared the latest frontier LLMs on identical phenotype-gene and phenotype-SNP generation tasks [13,19]. Second, existing evaluations often stop at the task or answer level rather than checking each generated relation against external genomic databases [21,24]. The external verifiability of LLM-generated phenotype-genotype relations, including identifier validity, phenotype normalization, and association-level evidence support, has not been evaluated in a systematic way, even as tool-augmented approaches are being developed to address this need [14,25,26]. This study addresses both gaps by benchmarking four frontier LLMs on six zero-shot forward and reverse phenotype-gene and phenotype-SNP generation tasks and verifying each generated association against curated external databases.

## MATERIALS AND METHODS

### Question Design

To systematically evaluate LLM performance across multiple dimensions of phenotype-genotype knowledge generation, we designed six zero-shot task categories covering different genomic entity types, disease prevalence, and query directions. Specifically, the benchmark assessed associations at both the gene and SNP levels, included common and rare disease phenotypes, and evaluated forward (phenotype-to-entity) and reverse (entity-to-phenotype) generation tasks. Curated phenotype, gene, and SNP inputs were selected to ensure representation across diverse disease domains and biological contexts. The complete input lists are provided in Table 1.

**Table 1.** Curated phenotype, gene, and SNP inputs for the six zero-shot query categories (the four columns are independent input sets, not row-wise associations).

| Common | Rare | Gene | SNP |
| --- | --- | --- | --- |
| Rheumatoid arthritis | Zellweger syndrome | APOE | rs3184504 |
| Hypertension | AGel amyloidosis | AKT1 | rs143334272 |
| Osteoporosis | Sanjad-Sakati syndrome | TP53 | rs72865845 |
| Coronary artery disease | Pituitary gigantism | VEGFA | rs6461667 |
| Type 2 diabetes | Opsismodysplasia | IL6 | rs78003935 |
| Lupus | Ectopia lentis syndrome | ERBB4 | rs35704505 |
| LDL cholesterol | Lamellar ichthyosis | SNED1 | rs2532859 |
| Alzheimer's disease | Camurati-Engelmann disease | GNA14 | rs6777684 |
| Lung cancer | Dysequilibrium syndrome | TAP2 | rs4101150 |
| Asthma | TEEBI hypertelorism syndrome | ESR1 | rs9349205 |

The inputs in Table 1 were curated to span the full range of evidence density in genomic knowledge bases. The common-disease column comprised ten frequent polygenic conditions, including type 2 diabetes and Alzheimer’s disease, all extensively represented in genome-wide association studies. The rare-disease column extended toward the sparse end, from well-described disorders such as Zellweger syndrome to ultra-rare Mendelian conditions such as TEEBI hypertelorism syndrome, which is documented in OMIM but essentially absent from the GWAS Catalog. The gene and SNP columns followed the same gradient, from extensively studied entities such as TP53 and APOE to less characterized ones such as SNED1 and GNA14, and from heavily catalogued variants such as rs3184504 to rarely catalogued ones such as rs9349205.

### Prompting Protocol and Response Handling

All model queries were driven by prompt text defined in code rather than interactive chat. For each task category, a fixed user message required the model to return a single JSON object. System prompts were task-conditioned: phenotype-related tasks for common and rare diseases used two distinct prompts, and gene-phenotype and SNP-phenotype tasks used a shared common-disease prompt across all models. Each input received one completion per task. JSON arrays were automatically expanded into tabular rows retaining model identifier, task category, and source input fields. Outputs were parsed automatically; decoding failures were logged and skipped, and free-text outputs were not manually edited before entry into the pipeline. Full prompt templates are provided in Supplementary File S1.

### Model Querying

Four LLMs were accessed via vendor APIs: GPT-5.5, Claude Sonnet 4.6, Gemini 3 Flash Preview, and DeepSeek V4 Flash. Web search was enabled for OpenAI, Anthropic, and Google; DeepSeek did not support web search in this configuration and was queried without it. Claude requests were configured with a maximum output of 32,000 tokens because no default token limit was available in the tested API configuration; all other sampling parameters were left at provider defaults. Reasoning depth was not explicitly controlled: GPT-5.5 used default reasoning.effort = medium, Claude Sonnet 4.6 used default effort = high, Gemini 3 Flash Preview used default dynamic thinking, and DeepSeek V4 Flash used the provider default (high) for chat completions. Because web-search capability, token budgets, and reasoning settings varied across vendors, this experiment should be interpreted as a comparison of different models under their closest available default API settings.

### Verification Framework

Figure 1 shows an overview of the verification pipeline. (1) Input & preprocessing: each generated association was verified through a multistage pipeline. (2) Phenotype normalization & ontology mapping: phenotype terms were normalized by lowercasing, removing modifier terms, and replacing common synonyms, mapped to EFO [27] as the primary ontology and MONDO [28] as fallback (Supplementary Data S3). (3) Entity validation: gene symbols and SNP identifiers were validated against Ensembl [29]; associations with unconfirmed entities were recorded as not found and excluded from further verification. (4) GWAS verification: for confirmed entities, gene outputs were matched against mapped and reported gene fields in the GWAS Catalog, and SNP outputs were matched against rsID fields. (5) OMIM verification: OMIM was queried using gene-based, SNP-bridged, or SNP-context lookup depending on task type. (6) Evidence Fusion & Outputs: GWAS and OMIM evidence were then combined into a fused evidence label (see Evaluation Metrics). Associations lacking external support were labeled unmatched, while those with unconfirmed entities were labeled not found. The final outputs comprised validated association-level tables, model-specific and task-specific summary statistics, and a detailed audit trail.

**Figure 1.**
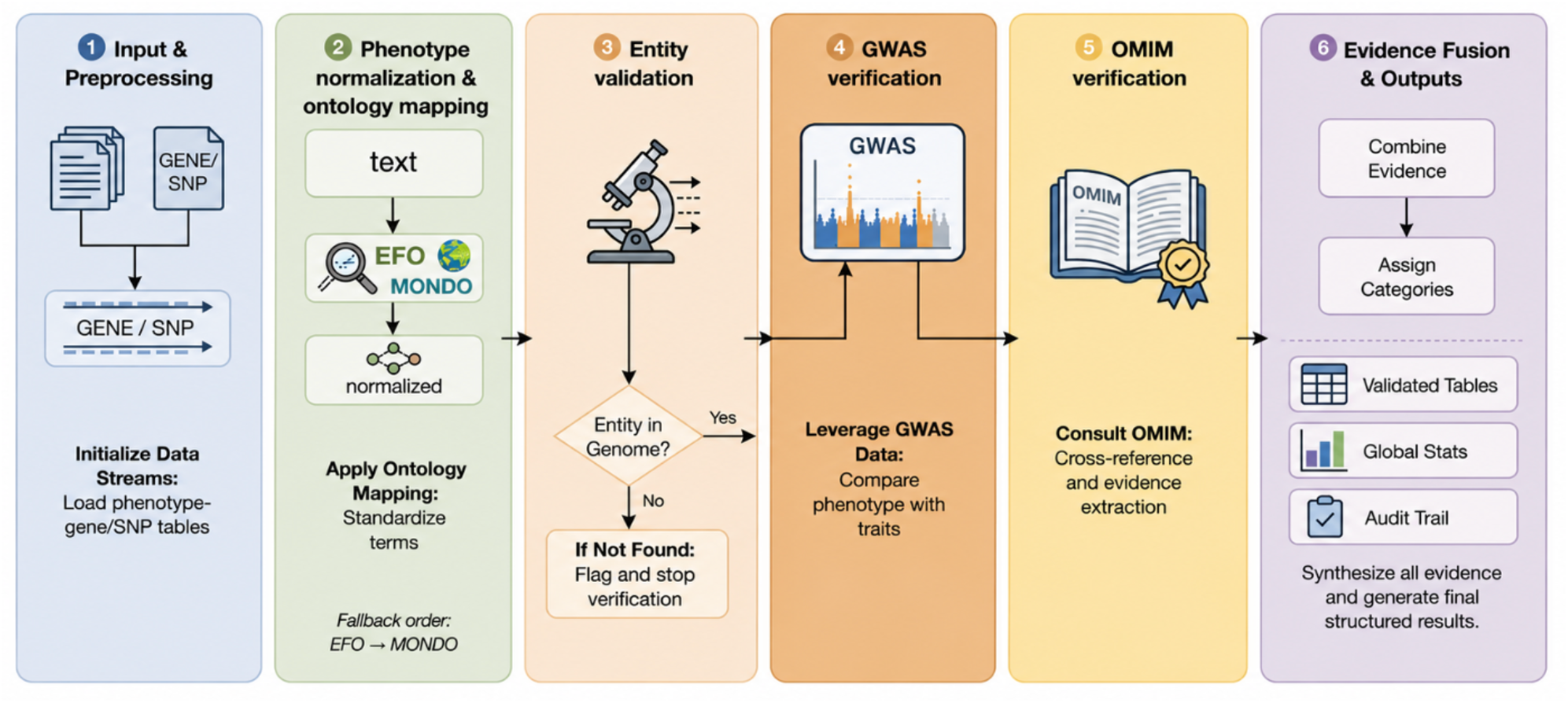
Overview of the phenotype-gene and phenotype-SNP verification pipeline.

### Evaluation Metrics

We evaluated generated phenotype-genotype associations according to their level of support in established and external genomic knowledge bases. Because no comprehensive reference standard exists for all phenotype-genotype relationships, particularly for rare diseases and SNP-level associations, we assessed whether each generated association could be supported by evidence from the GWAS Catalog and OMIM following phenotype normalization and genomic identifier validation.

We proposed fused evaluation metrics based on GWAS and OMIM. All generated associations were matched in the GWAS step at two levels. An exact GWAS phenotype match required identical strings, substring containment, or a phenotype token of ≥4 characters appearing within the GWAS trait string. A partial GWAS phenotype match required at least one shared token of ≥3 characters when the exact-match criterion was not met. GWAS support was labeled strong when the exact-match criterion was satisfied and p<5×10^-8 [30], weak when only a partial phenotype match was found, and none otherwise. OMIM support was labeled strong for direct phenotype matches, weak for partial or indirect matches, and none when no relevant evidence was found. The two sources were combined into a fused evidence level according to Table 2. Associations with strong or moderate fused evidence are termed documented throughout the manuscript, indicating external support from the GWAS Catalog or OMIM after Ensembl entity confirmation.

**Table 2.**
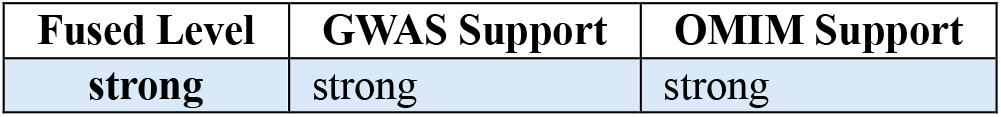
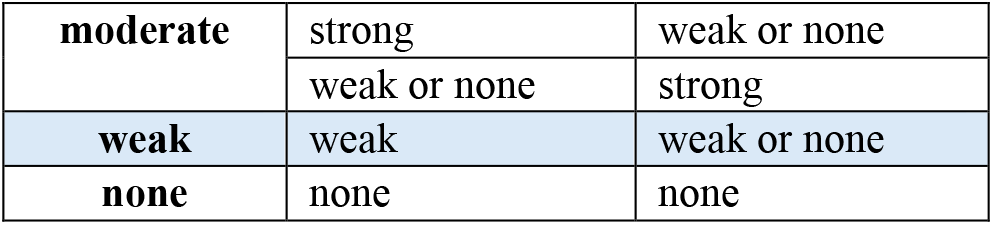
Fusion rules for assigning fused evidence levels from GWAS and OMIM support.

### Manual Validation of Unmatched Associations

We manually checked a random sample of 30 cases, 15 from the phenotype to gene outputs and 15 from the phenotype to SNP outputs, using a fixed random seed so the sample could be reproduced. We sorted each case into one of four groups. A case counted as retrieval failure if the association actually appeared in GWAS Catalog or OMIM but our pipeline simply failed to pull it out because of how the pipeline was built. A case counted as ontology mapping mismatch if the association was in GWAS Catalog or OMIM but listed under a different term or level of detail, for example a synonym, as long as the phenotype itself was the same. A case counted as incomplete database coverage if it was missing from GWAS Catalog and OMIM but could still be found with evidence in another established database. A case counted as model hallucination if no database had any supporting evidence at all, including cases where a real variant was simply linked to the wrong disease.

## RESULTS

### Model-generated relation corpus

Before external verification, we first examined the raw relation dataset generated by the four models. As shown in Figure 2, GPT-5.5 produced the largest number of associations overall, largely driven by the phenotype-gene-common task, whereas Claude Sonnet 4.6 and Gemini 3 Flash Preview generated intermediate-sized datasets, and DeepSeek V4 Flash produced the fewest associations overall. Across all models, the number of associations differed substantially by task. The phenotype-gene-common task produced the largest number of associations, while the phenotype-gene-rare, gene-phenotype, phenotype-snp-rare, and SNP-phenotype tasks produced far fewer. On the SNP side, the phenotype-snp-common and phenotype-snp-rare tasks produced a relatively comparable number of associations across models. In contrast, the SNP-phenotype task was more variable, with Gemini 3 Flash Preview and DeepSeek V4 Flash generating noticeably more associations than Claude Sonnet 4.6 and GPT-5.5. This higher variability may partly reflect the heterogeneous input composition of the SNP-phenotype task, which included both common and rare SNPs, whereas phenotype-snp-rare was restricted to rare phenotypes.

**Figure 2.**
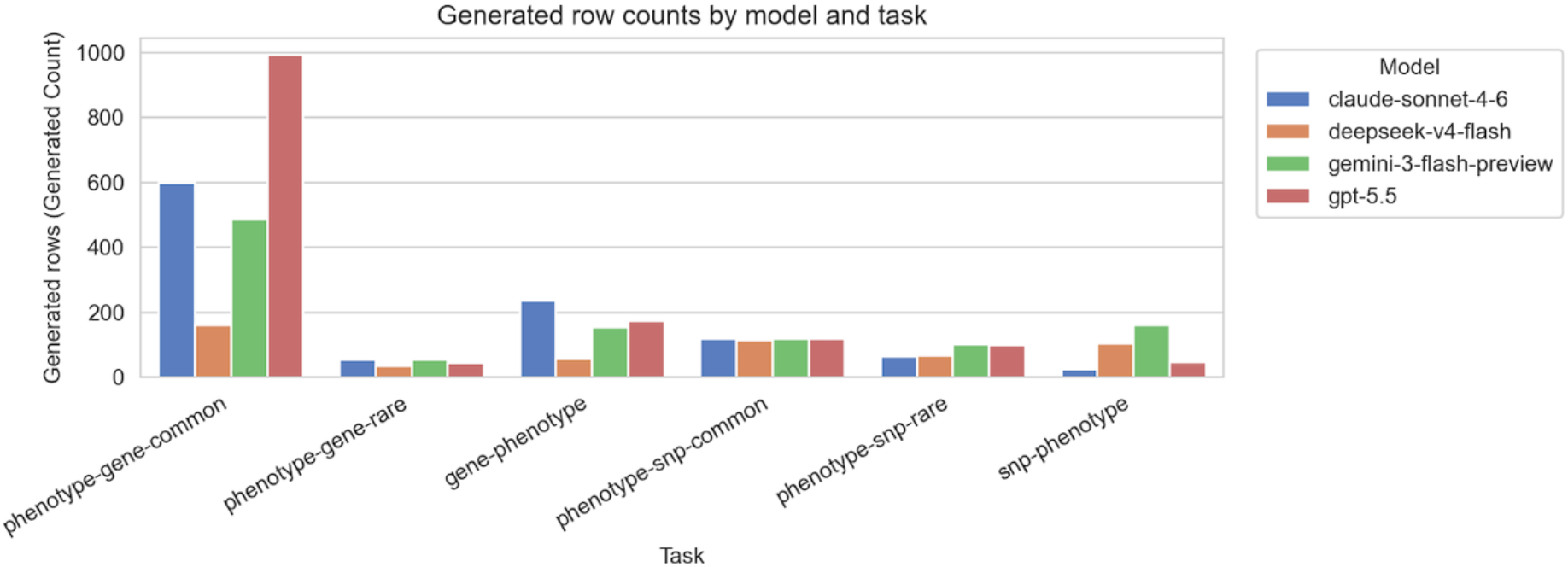
Distribution of generated relation counts by model across six task categories.

### Model-level performance

We next compared model-level performance across all generated relations. As shown in Figure 3A, GPT-5.5 produced the largest number of matched associations in absolute terms, followed by Claude Sonnet 4.6 and Gemini 3 Flash Preview, while DeepSeek V4 Flash generated the smallest dataset overall. Despite differences in output size, matched associations dominated across all models, with relatively smaller proportions of unconfirmed entities and unmatched entries. Notably, the absent fraction was small: overall, 97.26% of generated gene and SNP identifiers were confirmed as real entities in Ensembl, indicating that the generated genes and SNPs almost always corresponded to real genomic entities rather than fabricated identifiers. Whether each real entity was paired with a correct phenotype was examined through the manual validation procedure described in Methods. Of the 30 sampled unmatched relations, 9 (30.0%) reflected incomplete database coverage, 1 (3.3%) reflected ontology mapping mismatches, 2 (6.7%) reflected retrieval failures, and 18 (60.0%) were classified as model hallucination, including cases in which a model returned a real variant associated with a different disease. Although hallucinations were confined to the unmatched subset rather than the full set of generated associations, they carry clinical weight because an unsupported relation cannot be distinguished from a documented one at the point of use and could contribute to pathogenicity misclassification. Coupling generation with automated verification, requiring source attribution, and treating single-model outputs as hypotheses rather than conclusions can mitigate this risk. Although fabricated identifiers were rare, association-level errors were more frequent, indicating that unmatched associations arose from a mixture of database limitations (12 of 30) and true model errors (18 of 30). Full per-association annotations are provided in Supplementary Data S4. Within the matched subset, evidence levels were strongly skewed toward moderate support across all models (Figure 3B). Strong evidence accounted for only a small fraction of matched relations, while weak evidence remained limited. Most documented associations met at least one evidence criterion but not both, which accounts for the dominance of moderate over strong labels.

**Figure 3.**
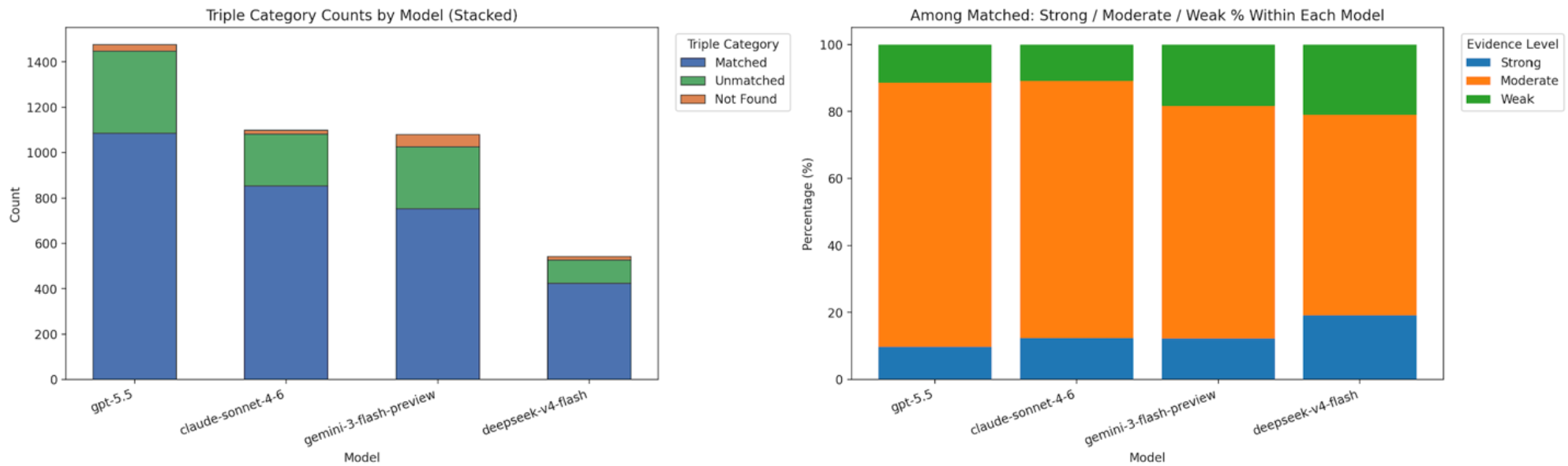
(A) Distribution of verification outcomes across models. (B) Distribution of evidence levels among matched relations across models.

To further evaluate the practical relevance of model outputs, we compared the proportion of documented associations (strong + moderate) across gene and SNP tasks (Figure 4). For gene-related associations, all models achieved consistently high documentation rates, ranging from approximately 64.5% to 76.5%. Among existing associations, SNP-relations showed substantially lower and more variable documentation rates, particularly for Gemini 3 Flash Preview (41.7%) and DeepSeek V4 Flash (48.6%), whereas Claude Sonnet 4.6 and GPT-5.5 retained higher and now nearly identical SNP documentation rates of 67.1% and 67.5%, respectively. DeepSeek V4 Flash showed the highest gene documentation rate but no corresponding advantage for SNP outputs, and across all models, gene-related associations were documented more consistently than SNP-related outputs. Chi-square tests with Bonferroni correction showed that GPT-5.5 did not differ significantly from either Claude Sonnet 4.6 or DeepSeek V4 Flash, and Gemini 3 Flash Preview did not differ significantly from DeepSeek V4 Flash; Claude Sonnet 4.6 differed significantly from both Gemini 3 Flash Preview and DeepSeek V4 Flash, and GPT-5.5 differed significantly from Gemini 3 Flash Preview (Table 3). Detailed validation summaries are provided in Supplementary Tables S1-S4, and the full relation-level outputs are provided in Supplementary Data S1 and S2.

**Table 3.** Chi-square tests and pairwise comparisons of documentation rates across four LLMs.

| Model | Overall |  | Gene tasks |  | SNP tasks |  |
| --- | --- | --- | --- | --- | --- | --- |
|  | doc/total | Rate (%) | doc/total | Rate (%) | doc/total | Rate (%) |
| Claude Sonnet 4.6 | 760/1099 | 69.2 | 619/889 | 69.6 | 141/210 | 67.1 |
| DeepSeek V4 Flash | 334/541 | 61.7 | 195/255 | 76.5 | 139/286 | 48.6 |
| Gemini 3 Flash Preview | 614/1079 | 56.9 | 454/695 | 65.3 | 160/384 | 41.7 |
| GPT-5.5 | 961/1477 | 65.1 | 782/1212 | 64.5 | 179/265 | 67.5 |
| Overall: $\chi^2(3) = 37.72$, $p = 3.24 \times 10^{-8}$ | | | | | | |
| Gene subgroup: $\chi^2(3) = 17.37$, $p = 5.92 \times 10^{-4}$ | | | | | | |
| SNP subgroup: $\chi^2(3) = 61.06$, $p = 3.48 \times 10^{-13}$ | | | | | | |

| Pairwise comparison | $\chi^2$ | df | Raw p | Bonferroni-corrected p |
| --- | --- | --- | --- | --- |
| Claude Sonnet 4.6 vs DeepSeek V4 Flash | 7.50 | 1 | 0.0062 | 0.0370 |
| Claude Sonnet 4.6 vs Gemini 3 Flash Preview | 24.65 | 1 | $6.87 \times 10^{-7}$ | $4.12 \times 10^{-6}$ |
| Claude Sonnet 4.6 vs GPT-5.5 | 4.24 | 1 | 0.0396 | 0.2376 |
| DeepSeek V4 Flash vs Gemini 3 Flash Preview | 1.63 | 1 | 0.2024 | 1.0000 |
| DeepSeek V4 Flash vs GPT-5.5 | 1.40 | 1 | 0.2367 | 1.0000 |
| Gemini 3 Flash Preview vs GPT-5.5 | 10.57 | 1 | 0.0012 | 0.0069 |

Documentation rates and chi-square tests were computed across all generated associations. doc = documented (strong + moderate); df = degrees of freedom.

**Figure 4.**
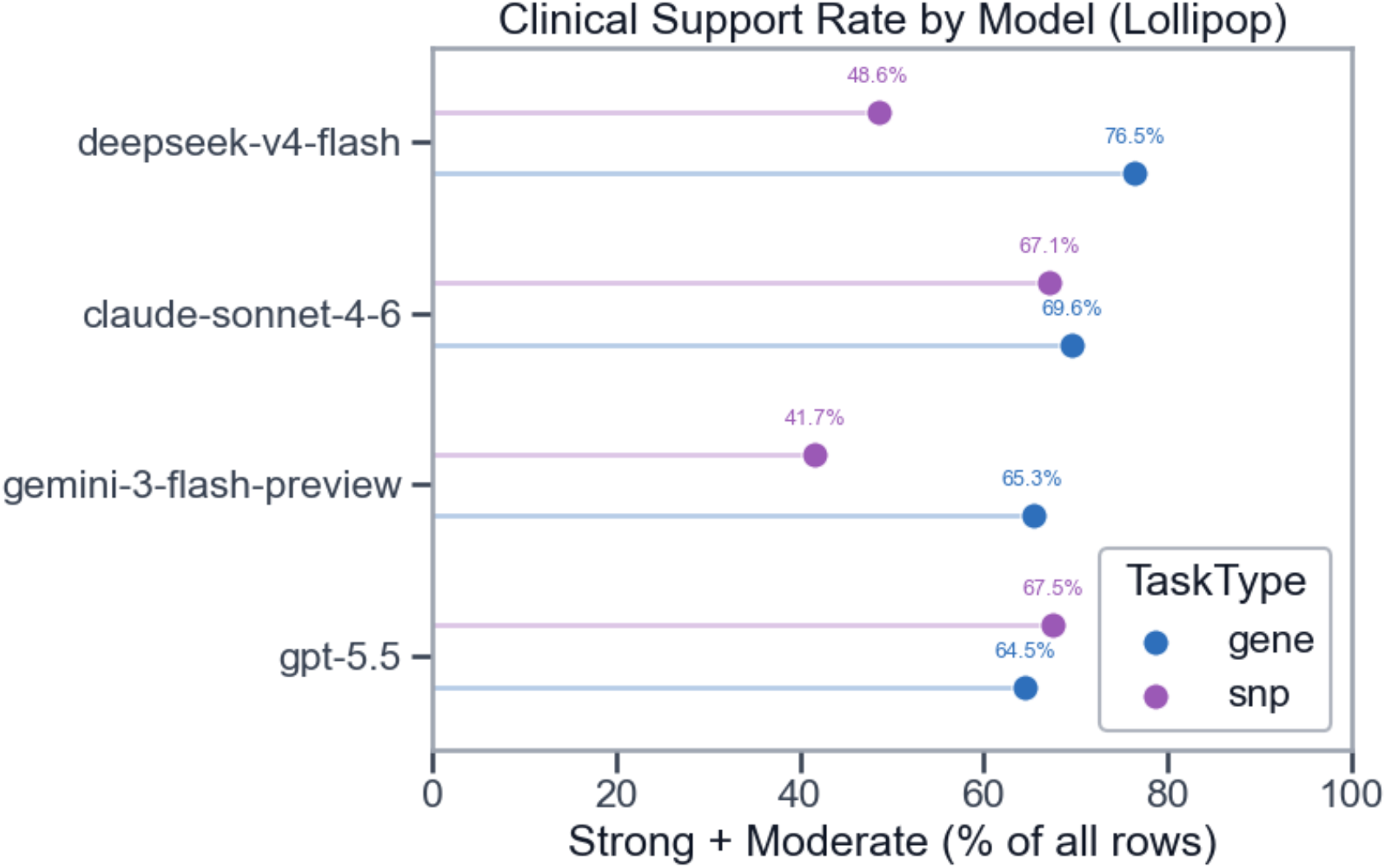
Documentation rates of model-generated gene and SNP associations across all generated associations.

### Task-level performance across six categories

Matched rates varied markedly across the six task categories (Supplementary Figure S1). Overall, gene-related tasks performed better than SNP-related tasks, though the gap was modest (76.2% vs 68.8%). Phenotype-gene-common showed consistently high matched rates across all models, and phenotype-gene-rare also remained relatively well supported. In contrast, SNP-related rates were far more variable: phenotype-snp-common achieved high matched rates across all models, whereas phenotype-snp-rare showed by far the weakest performance, with matched relations recovered only sporadically by three of the four models and none by DeepSeek V4 Flash. SNP-phenotype, by contrast, achieved a notably higher matched rate than either phenotype-snp task, indicating that the reverse SNP-to-phenotype direction was not uniformly harder than the forward direction. Directional differences were in fact task-dependent rather than uniformly favoring forward prompts: gene-phenotype remained somewhat lower than the corresponding phenotype-gene tasks, consistent with forward prompts being easier for gene-related generation, but SNP-phenotype outperformed the pooled phenotype-SNP tasks, reversing this pattern for SNP-related generation.

### GWAS and OMIM evidence patterns

As shown in Supplementary Figure S2, GWAS and OMIM contributed unevenly across task types. In gene-related tasks, OMIM matched some documented relations, but the large majority of relations with strong GWAS evidence had no corresponding OMIM match. In SNP-related tasks, OMIM contributed substantially rather than being absent: OMIM strong or weak evidence was present in roughly two-thirds of SNP-related rows, alongside GWAS Catalog documentation. Figure 5 further shows that strong labels required both sources, moderate labels came mainly from GWAS alone, and weak labels drew on either source, with OMIM-only cases more common. This pattern is consistent with the different scope of the two databases: GWAS Catalog captures many common variant-trait associations, whereas OMIM focuses mainly on Mendelian gene-phenotype associations and does not systematically index variant-level associations outside known Mendelian loci.

**Figure 5.**
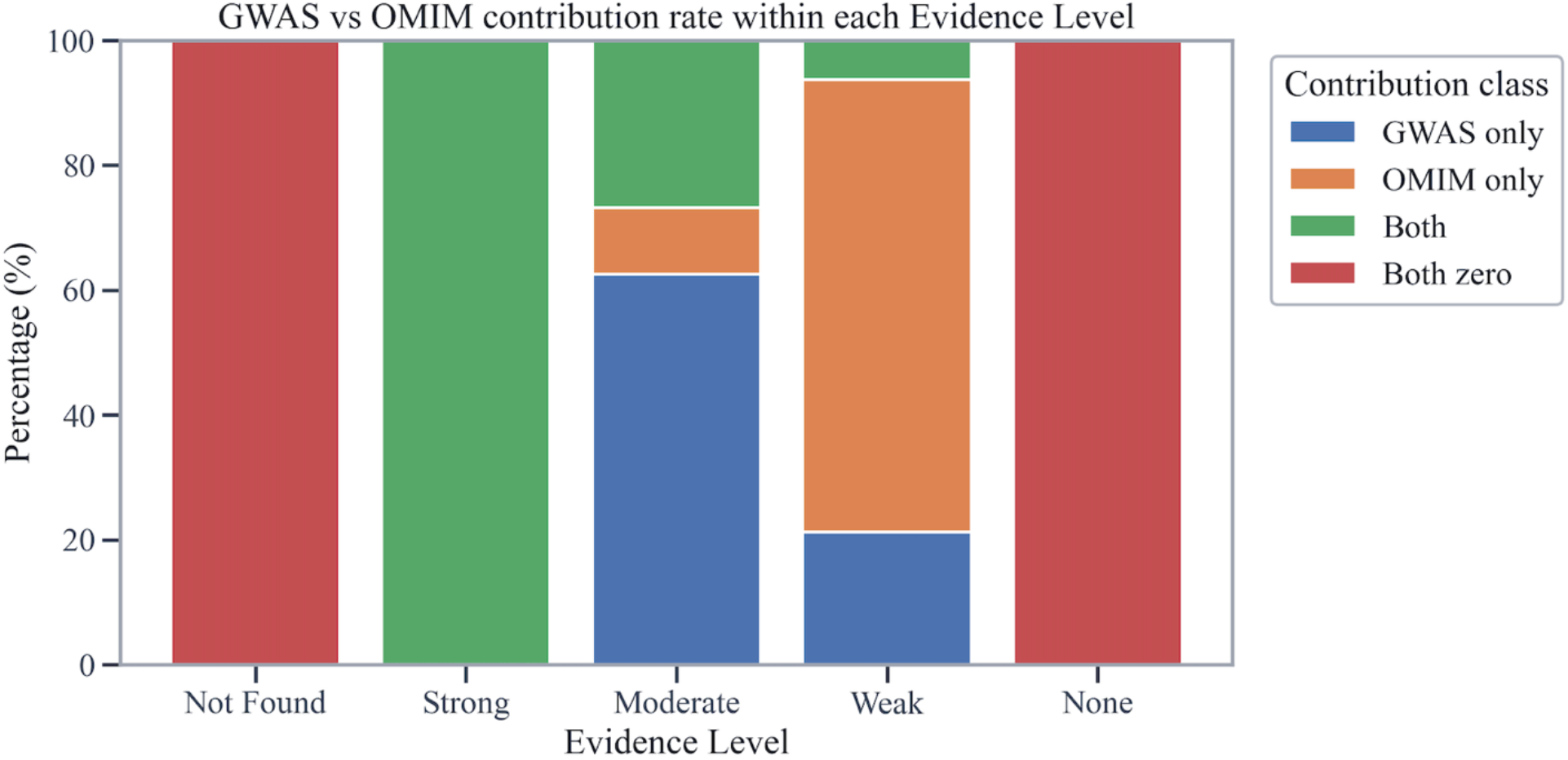
Contribution of GWAS and OMIM evidence sources within each evidence level.

### Cross-Model Differences in Generation and Verification

Consistency of generated relations across models was further analyzed by examining overlap within each task category. As shown in Figure 6, the degree of agreement varied substantially across tasks. Within phenotype-gene tasks, overlap was heterogeneous: phenotype-gene-common showed a low four-model overlap of 9.1%, with only 120 of 1,322 distinct associations shared by all four models, whereas phenotype-gene-rare showed much higher overlap, with 36 of 63 associations, or 57.1%, shared across all four models. For reverse gene-phenotype generation, overlap remained low at 2.8%, with only 14 of 494 generated associations shared by all four models, reflecting the increased difficulty of reconstructing phenotype descriptions from genetic entities. This variability may also reflect the inherent complexity of genotype-phenotype relationships in clinical practice, where a given variant can be associated with phenotypes not manifested in every patient. A different pattern was observed in SNP-related tasks. For phenotype-snp-common and phenotype-snp-rare, all four models covered the same 10 input phenotypes, producing complete input-level overlap of 100%, although this does not imply complete overlap of the generated SNP associations. In contrast, the SNP-phenotype task exhibited low overlap at 2.4%, with only 7 of 286 generated associations shared across all four models, indicating substantial variability in reverse SNP-phenotype generation.

**Figure 6.**
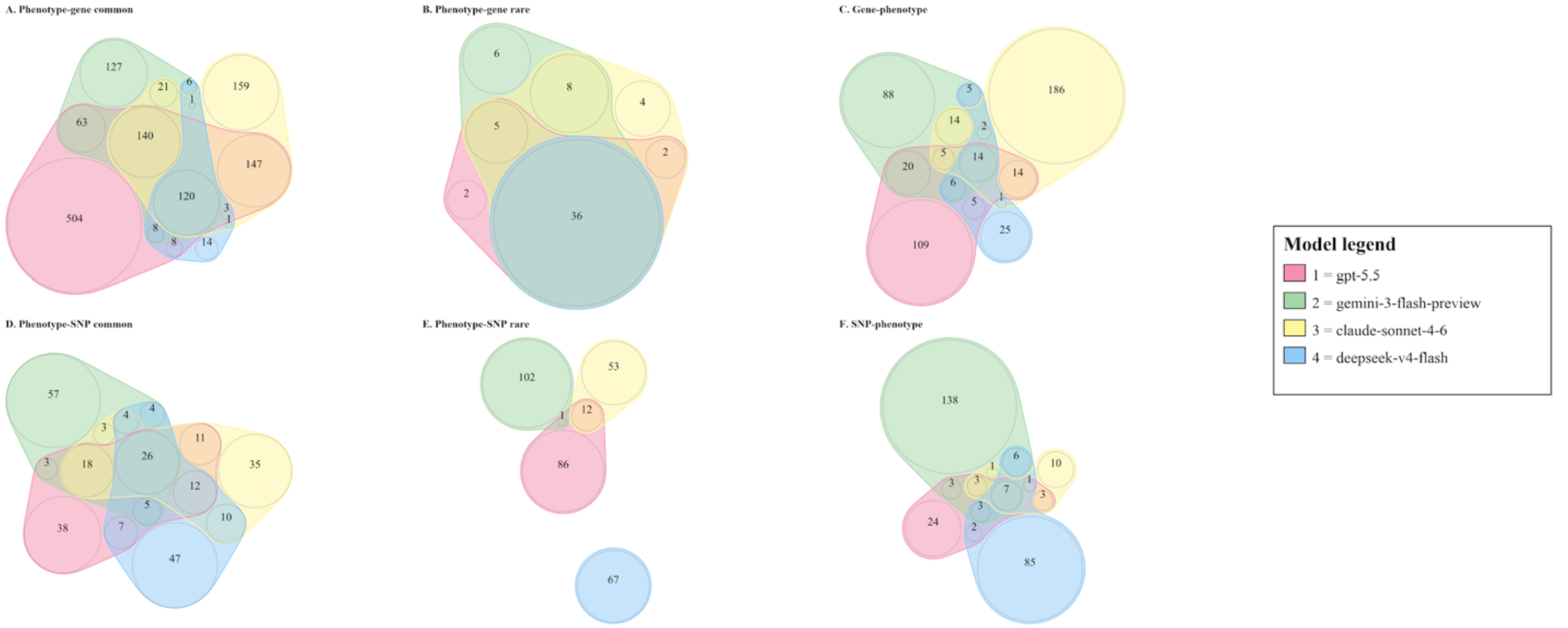
Weighted Venn diagrams showing cross-model overlap of generated phenotype-gene and phenotype-SNP associations across six task categories.

## DISCUSSION

This study directly addresses two gaps identified in prior genomic LLM evaluations: the lack of head-to-head comparisons among frontier models on identical phenotype-genotype generation tasks, and the lack of relation-level verification against external genomic databases. By evaluating four models across six zero-shot task categories and assigning evidence labels to each generated association, our benchmark separates model generation behavior from database-supported documentation and provides a more granular assessment of genomic reliability.

### Interpreting Cross-Model Differences

The four models differed markedly in both the volume of associations generated and the proportion documented by external evidence: GPT-5.5 generated the broadest set of candidate relations while maintaining a documentation rate competitive with the other models, including for SNP-related tasks, whereas DeepSeek V4 Flash produced the smallest output. Output volume did not translate directly into documentation rate. Model differences depended strongly on relation type: DeepSeek V4 Flash showed the highest gene documentation rate but lower SNP documentation, whereas Claude Sonnet 4.6 and GPT-5.5 showed stronger SNP documentation. These differences likely reflect each model’s default generation behavior, including output length and reasoning configuration, rather than any single controlled factor. One possible explanation is that gene- and SNP-related tasks draw on structurally different knowledge bases: gene-phenotype relations are well captured by Mendelian disease resources such as OMIM, whereas SNP-phenotype relations depend more heavily on population-scale GWAS evidence. A model that encodes one body of evidence more completely may therefore perform well on one task type but not the other. SNP identifier instability across database builds may further contribute to the variability observed in SNP tasks. These hypotheses remain speculative given that the training data composition of all four models is not publicly disclosed.

### Cross-Model Consensus and Knowledge Scope

Low overlap between models does not mean the differing outputs are wrong. In open-ended tasks such as phenotype-gene or phenotype-SNP generation, the candidate space is large and many associations may be valid, so different models can document different but legitimate subsets depending on how evidence appears in their training data. The knowledge of these models may also reach beyond the databases used for validation: trained on large biomedical corpora, they can encode associations reported in the literature but not yet curated into OMIM or the GWAS Catalog. An association that fails verification may then lie outside database coverage rather than outside valid biology. High overlap is not always strong evidence either. For some SNP tasks, all four models generated largely the same associations around well-known variants such as rs3184504, which are heavily represented in the literature and training data. This apparent agreement may reflect shared exposure to prominent associations rather than independent confirmation, and may not extend to less-studied variants. These patterns shape how consensus should be used. When two or more models independently generate the same association and it also appears in the GWAS Catalog or OMIM, this agreement offers a practical filter for prioritizing candidates without extra manual curation. Associations from a single model should not be discarded automatically, since they may reflect broader recall or valid relations that the selected databases cover poorly, and instead warrant case-by-case review. Model agreement is informative but not definitive, and its absence should be read in light of task type, database coverage, and the number of possible associations. Because an unverifiable association cannot be told apart from a confident error at the point of use, relation-level verification remains a practical safeguard against the pathogenicity misclassification and unsupported clinical summaries that motivated this work.

### Database-Dependent Validation

Documentation rates depended heavily on how well the chosen reference databases covered each task type. The two resources used here cover different, limited parts of genomic evidence. OMIM is oriented toward curated, largely monogenic gene-phenotype and Mendelian disease relationships, where a single gene is linked to a well-characterized clinical phenotype [5]. The GWAS Catalog, by contrast, aggregates common variant-trait associations that reach genome-wide significance in population-scale studies, and is therefore weighted toward frequent variants of modest individual effect [23]. Gene-level and common-disease tasks fell within this combined coverage and could usually be matched, though matching depended on querying the current HGNC-approved symbol rather than a deprecated alias [22]. SNP-level tasks showed uneven alignment overall, and the gap widened sharply for rare disease phenotypes. Common SNP tasks overlapped extensively with the GWAS Catalog, since frequent variants are exactly what genome-wide studies are powered to detect. Rare disease phenotypes, however, are systematically underrepresented: conventional genome-wide designs have limited statistical power at low allele frequencies and typically apply frequency thresholds that exclude such variants altogether [30], so rare phenotype-SNP relationships rarely appear in the Catalog, and where documented at all sit in resources outside this pipeline [2,6]. Gene-level verification is therefore more reliable than SNP-level verification, because the available databases curate gene-disease relationships far more completely than rare variant associations.

### Limitations

The pipeline relied on Ensembl-based identifier retrieval and evidence from the GWAS Catalog and OMIM. This allowed a structured comparison across models but limited the ability to distinguish between nomenclature drift, alias mismatch, under-documented associations, and unsupported or possibly incorrect relations. Future work should incorporate additional reference databases. For rare-disease tasks, disease-specific locus databases and sequencing-derived variant resources would be needed to provide adequate coverage. Adding these resources would help separate reasons for non-documentation and improve the interpretability of documentation rates across task types. Beyond database coverage, the manual review of unmatched associations found that hallucinations were the dominant failure mode, taking three forms: fabricated identifiers absent from any database, digit-level transcription errors in real rsIDs, and valid identifiers linked to unrelated diseases. Mitigation strategies such as identifier pre-validation and retrieval-augmented generation could reduce but not eliminate this risk.

## CONCLUSION

This study shows that relation-level validation against curated biomedical databases is necessary to assess the reliability of LLM-generated phenotype-genotype associations, and that model-level evaluation alone is insufficient for this purpose. Across four frontier models, performance varied by output volume, relation type, task direction, and evidence availability. Gene-level and common disease associations were generally more documentable than SNP-level and rare disease associations, while rare phenotype-SNP generation exposed limitations in both LLM knowledge and current validation databases.

These findings highlight that identifier validity and association validity are distinct: a gene or SNP may be documented in Ensembl without the corresponding phenotype association being supported by GWAS Catalog or OMIM. Overall, LLMs may serve as useful tools for generating candidate genomic associations, but their outputs should be treated as partially documented hypotheses requiring structured database verification, especially for rare disease phenotypes, where current databases remain incomplete.

## SUPPLEMENTARY MATERIAL

Supplementary Table S1. Gene-task Ensembl identifier validation summary by model. Supplementary

Table S2. SNP-task Ensembl identifier validation summary by model. Supplementary

Table S3. Gene-task evidence distribution by model.

Supplementary Table S4. SNP-task evidence distribution by model.

Supplementary Data S1. Full phenotype-gene relation-level validation output.

Supplementary Data S2. Full phenotype-SNP relation-level validation output.

Supplementary Data S3. Phenotype normalization and ontology mapping results.

Supplementary Data S4. Manual validation of 30 unmatched associations.

Supplementary Figure S1. Task-level matched rates by model across six task categories.

Supplementary Figure S2. GWAS and OMIM evidence source contributions by task category.

Supplementary File S1. Prompt templates used for model querying.

