## Supplementary material for "Assessing the Reliability of LLM-Generated Phenotype-Genotype Associations Through External Validation": 8models_Generation.ipynb

````
# %% [code]
import csv
import json
import os
import re
import time
from typing import Optional

from openai import OpenAI

# --- Configuration ---
OUT_PREFIX = ""
OPENAI_WEB_SEARCH_TOOLS = [{"type": "web_search"}]
OPENAI_REASONING: Optional[dict] = None  # None = provider default
OPENAI_ENABLE_WEB_SEARCH = True
OPENAI_REQUEST_TIMEOUT_SEC = 600.0
GEMINI_THINKING_BUDGET: Optional[int] = None  # None = provider default
GEMINI_MAX_RETRIES = 6
GEMINI_RETRY_BASE_SEC = 8
FLUSH_CSV_AFTER_EACH_TASK = True  # write CSV after each task finishes

ANTHROPIC_WEB_SEARCH_TOOLS = [
    {"type": "web_search_20250305", "name": "web_search", "max_uses": 5},
]

# --- Task scope ---
# SANITY_CHECK_TASK: None = all six tasks
SANITY_CHECK_TASK = None
SANITY_CHECK_TASK_KEYS: Optional[list[str]] = None
N_RUNS = 1  # repeats per input (run_number 1..N_RUNS)
EXPORT_CSV_TASK_KEYS: Optional[list[str]] = None  # None = all six CSVs per model
OPENAI_GPT55_USE_SPLIT_TASK_KEYS = False
OPENAI_GPT55_TASK_KEYS: Optional[list[str]] = None

SANITY_TASK_KEYS = frozenset(
    {
        "phenotype_to_snp_common",
        "phenotype_to_snp_rare",
        "phenotype_to_gene_common",
        "phenotype_to_gene_rare",
        "gene_to_phenotype",
        "SNP_to_phenotype",
    }
)


def _env_key(*names: str) -> str:
    # Read the first non-empty API key from environment variables.
    for name in names:
        value = os.environ.get(name, "").strip()
        if value:
            return value
    return ""


def _slice_items(items, max_items):
    if max_items is None:
        return items
    return items[:max_items]


# --- System prompts and input lists ---
SYSTEM_PROMPT_COMMON = (
    "You are a human medical geneticist specializing in common, complex traits and population-level disease genetics "
    "(e.g. GWAS, polygenic architecture, common variants). "
    "Your sole task is to return valid JSON. "
    "Do not include any explanations, disclaimers, markdown code fences, or text outside the JSON object. "
    "Use HGNC official gene symbols for human genes. "
    "Prioritize top-supported associations first (e.g. genome-wide significant, replicated GWAS, or strong clinical evidence), "
    "then add further supported items in decreasing strength. "
    "Do not impose a maximum number of items or any character/word limit; omit anything you cannot support."
)

SYSTEM_PROMPT_RARE = (
    "You are a human medical geneticist specializing in rare disease and Mendelian genetics "
    "(monogenic causes, syndromes, ultrarare variants, OMIM-style gene–disease relationships). "
    "Your sole task is to return valid JSON. "
    "Do not include any explanations, disclaimers, or text outside the JSON object. "
    "You must not return any surrounding text, markdown formatting, or comments. "
    "Use HGNC official gene symbols for human genes. "
    "Prioritize top-supported associations first (e.g. strong Mendelian/clinical or functional evidence), "
    "then add further supported items in decreasing strength. "
    "Do not impose a maximum number of items or any character/word limit; omit anything you cannot support."
)


SNP_JSON_RULES = (
    "Order SNPs from strongest to weaker evidence for this phenotype. "
    "Each entry must be one string GENE_SYMBOL:rs######## (HGNC; lowercase rs). "
    "No limit on how many; include all you can justify."
)

GENE_JSON_RULES = (
    "Order genes from strongest to weaker evidence for this phenotype. "
    "HGNC symbols only. No limit on how many; omit if uncertain."
)

PHENO_JSON_RULES = (
    "Order phenotypes from strongest to weaker evidence. "
    "Use standard disease/trait names. No limit on how many; include all you can justify."
)

PHENOTYPES_COMMON = [
    "Rheumatoid arthritis",
    "Hypertension",
    "Osteoporosis",
    "Coronary Artery Disease",
    "Type 2 diabetes",
    "Lupus",
    "LDL cholesterol",
    "Alzheimer's Disease",
    "Lung Cancer",
    "Asthma",
]

PHENOTYPES_RARE = [
    "Zellweger Syndrome",
    "AGel Amyloidosis",
    "Sanjad-Sakati Syndrome",
    "Pituitary Gigantism",
    "Opsismodysplasia",
    "Ectopia Lentis Syndrome",
    "Lamellar ichthyosis",
    "Camurati-Engelmann disease",
    "Dysequilibrium syndrome",
    "Teebi hypertelorism syndrome",
]

GENES_G2P = ["APOE", "AKT1", "TP53", "VEGFA", "IL6", "ERBB4", "SNED1", "GNA14", "TAP2", "ESR1"]

SNPS_P2P = [
    "rs3184504",
    "rs143334272",
    "rs72865845",
    "rs6461667",
    "rs78003935",
    "rs35704505",
    "rs2532859",
    "rs6777684",
    "rs4101150",
    "rs9349205",
]


# --- Provider clients ---
def openai_client():
    key = _env_key("OPENAI_API_KEY")
    if not key:
        raise RuntimeError("Set OPENAI_API_KEY in the environment.")
    return OpenAI(api_key=key, timeout=OPENAI_REQUEST_TIMEOUT_SEC)


def deepseek_client():
    key = _env_key("DEEPSEEK_API_KEY", "DEEPSEEK_KEY")
    if not key:
        raise RuntimeError("Set DEEPSEEK_API_KEY in the environment.")
    return OpenAI(api_key=key, base_url="https://api.deepseek.com/v1", timeout=180.0)


def _openai_via_responses(
    client: OpenAI, model: str, system: str, user: str, *, use_reasoning_effort: bool = False
) -> str:
    # OpenAI Responses API with optional web search and reasoning effort.
    prompt = f"[System instructions]\n{system}\n\n[User]\n{user}"
    kwargs = {"model": model, "input": prompt}
    if OPENAI_ENABLE_WEB_SEARCH:
        kwargs["tools"] = OPENAI_WEB_SEARCH_TOOLS
    if use_reasoning_effort and OPENAI_REASONING is not None:
        kwargs["reasoning"] = OPENAI_REASONING
    t0 = time.perf_counter()
    response = client.responses.create(**kwargs)
    print(f"[OpenAI] {model} done in {time.perf_counter() - t0:.1f}s", flush=True)
    return (response.output_text or "").strip()


def make_openai_caller(model: str, prefer_chat: bool = False):
    # Retry without reasoning effort if the first Responses call fails.
    client = openai_client()

    def call(system: str, user: str) -> str:
        if prefer_chat:
            return _openai_via_responses(
                client, model, system, user, use_reasoning_effort=False
            )
        try:
            return _openai_via_responses(
                client, model, system, user, use_reasoning_effort=True
            )
        except Exception as e_first:
            try:
                return _openai_via_responses(
                    client, model, system, user, use_reasoning_effort=False
                )
            except Exception as e_second:
                raise e_second from e_first

    return call


def make_deepseek_caller(model: str):
    # DeepSeek uses chat completions; no web search tool.
    client = deepseek_client()

    def call(system: str, user: str) -> str:
        response = client.chat.completions.create(
            model=model,
            messages=[
                {"role": "system", "content": system},
                {"role": "user", "content": user},
            ],
            response_format={"type": "json_object"},
            stream=False,
        )
        return (response.choices[0].message.content or "").strip()

    return call


def make_anthropic_caller(model: str):
    try:
        import anthropic
    except ImportError as e:
        raise RuntimeError("pip install anthropic") from e
    key = _env_key("ANTHROPIC_API_KEY")
    if not key:
        raise RuntimeError("Set ANTHROPIC_API_KEY in the environment.")
    client = anthropic.Anthropic(api_key=key)

    def call(system: str, user: str) -> str:
        # Claude with web search tool.
        _kw = dict(
            model=model,
            max_tokens=32000,
            system=system,
            messages=[{"role": "user", "content": user}],
            tools=ANTHROPIC_WEB_SEARCH_TOOLS,
        )
        with client.messages.stream(**_kw) as stream:
            msg = stream.get_final_message()
        text_out = ""
        for block in msg.content:
            if hasattr(block, "text") and block.text:
                text_out = block.text
        return text_out.strip() or str(msg.content[-1]).strip()

    return call


def make_gemini_caller(model: str):
    try:
        from google import genai
        from google.genai import types
    except ImportError as e:
        raise RuntimeError("pip install google-genai") from e
    key = _env_key("GOOGLE_API_KEY", "GEMINI_API_KEY")
    if not key:
        raise RuntimeError("Set GOOGLE_API_KEY or GEMINI_API_KEY in the environment.")
    client = genai.Client(api_key=key)

    def call(system: str, user: str) -> str:
        # Gemini with Google Search grounding; retry on transient errors.
        grounding_tool = types.Tool(google_search=types.GoogleSearch())
        cfg_kwargs = dict(
            system_instruction=system,
            tools=[grounding_tool],
        )
        if GEMINI_THINKING_BUDGET is not None:
            try:
                cfg_kwargs["thinking_config"] = types.ThinkingConfig(
                    thinking_budget=GEMINI_THINKING_BUDGET
                )
            except Exception:
                try:
                    cfg_kwargs["thinking_config"] = types.ThinkingConfig(
                        thinking_level="LOW"
                    )
                except Exception:
                    pass
        cfg = types.GenerateContentConfig(**cfg_kwargs)
        max_try = max(1, int(GEMINI_MAX_RETRIES))
        base = max(1, int(GEMINI_RETRY_BASE_SEC))
        for attempt in range(1, max_try + 1):
            try:
                r = client.models.generate_content(model=model, contents=user, config=cfg)
                return (r.text or "").strip()
            except Exception as e:
                msg = str(e).lower()
                code = getattr(e, "status_code", None)
                retryable = code in (429, 500, 502, 503, 504) or any(
                    s in msg for s in ("503", "unavailable", "429", "resource exhausted", "try again later")
                )
                if retryable and attempt < max_try:
                    wait_s = min(120, base * attempt)
                    print(f"[Gemini] retry {attempt}/{max_try} in {wait_s}s", flush=True)
                    time.sleep(wait_s)
                else:
                    raise

    return call


def make_caller(provider: str, model: str):
    # Route to the provider-specific caller.
    if provider == "openai_reasoning":
        return make_openai_caller(model, prefer_chat=False)
    if provider == "openai_regular":
        return make_openai_caller(model, prefer_chat=True)
    if provider == "anthropic":
        return make_anthropic_caller(model)
    if provider == "gemini":
        return make_gemini_caller(model)
    if provider == "deepseek":
        return make_deepseek_caller(model)
    raise ValueError(f"unknown provider: {provider}")


def run_all_models_and_write():
    # Run all entries in MODEL_CONFIGS and write CSVs after each model.
    b = empty_buckets()
    for cfg in MODEL_CONFIGS:
        run_one_model_and_write_csvs(cfg, b)
        print("done:", cfg["csv_model"])
    return b


# --- Model configs ---
# Web search: on for OpenAI / Anthropic / Gemini; off for DeepSeek.
MODEL_CONFIGS = [
    {
        "title": "## **1. OpenAI · gpt-5.5 (reasoning: API default; docs medium)**",
        "provider": "openai_reasoning",
        "model": "gpt-5.5",
        "csv_model": "gpt-5.5",
        "reasoning_level": "default",
    },
    {
        "title": "## **2. Anthropic · claude-sonnet-4-6 (web_search; effort API default; docs high)**",
        "provider": "anthropic",
        "model": "claude-sonnet-4-6",
        "csv_model": "claude-sonnet-4-6",
        "reasoning_level": "default",
    },
    {
        "title": "## **3. Google · gemini-3-flash-preview (thinking: API default; docs high)**",
        "provider": "gemini",
        "model": "gemini-3-flash-preview",
        "csv_model": "gemini-3-flash-preview",
        "reasoning_level": "default",
    },
    {
        "title": "## **4. DeepSeek · deepseek-v4-flash (API defaults; docs regular high, no web)**",
        "provider": "deepseek",
        "model": "deepseek-v4-flash",
        "csv_model": "deepseek-v4-flash",
        "reasoning_level": "default",
    },
]


def _cfg_is_openai_gpt55(cfg: dict) -> bool:
    return cfg.get("provider") == "openai_reasoning" and str(cfg.get("model", "")).strip() == "gpt-5.5"


def validate_openai_gpt55_format() -> None:
    req = ("title", "provider", "model", "csv_model", "reasoning_level")
    gpt_cfg = next((c for c in MODEL_CONFIGS if _cfg_is_openai_gpt55(c)), None)
    if gpt_cfg is None:
        raise ValueError("Missing gpt-5.5 entry in MODEL_CONFIGS")
    missing = [k for k in req if k not in gpt_cfg]
    if missing:
        raise ValueError(f"gpt-5.5 config missing keys: {missing}")
    if not OPENAI_WEB_SEARCH_TOOLS or OPENAI_WEB_SEARCH_TOOLS[0].get("type") != "web_search":
        raise ValueError("OPENAI_WEB_SEARCH_TOOLS must include web_search")


def _export_reasoning_level(cfg: dict) -> str:
    # Used in CSV filename: {model}_{reasoning}_{category}.csv
    if cfg.get("provider") == "openai_reasoning":
        if OPENAI_REASONING is None:
            return "default"
        return str(OPENAI_REASONING.get("effort", "none"))
    return str(cfg.get("reasoning_level", "none"))


def _strip_json_markdown(raw: str) -> str:
    # Extract JSON from fenced ```json blocks or plain code fences.
    t = (raw or "").strip()
    m = re.search(r"```(?:json)?\s*\n", t, flags=re.I)
    if m:
        end = t.find("```", m.end())
        if end != -1:
            return t[m.end() : end].strip()
    if t.startswith("```"):
        lines = t.splitlines()
        if lines:
            lines = lines[1:]
        while lines and lines[-1].strip().startswith("```"):
            lines = lines[:-1]
        t = "\n".join(lines).strip()
    return t


def _is_missing_item(v) -> bool:
    if v is None:
        return True
    t = str(v).strip().lower()
    return t in {"", "not found", "none", "null", "n/a", "na", "unknown"}


# --- Task runners: call model, parse JSON, append flattened rows ---
def _accum_rows_phenotype_snp(
    call, system_prompt, model_label, phenotypes, dest: list, *, n_runs: int = 1, max_items: Optional[int] = None
) -> None:
    for phenotype in _slice_items(phenotypes, max_items):
        for run_number in range(1, n_runs + 1):
            prompt = f"""What genetic variants are associated with {phenotype}?
Return a single JSON object with keys "phenotype" and "SNP" only. Example shape:
{{
  "phenotype": "{phenotype}",
  "SNP": ["GENE_SYMBOL:rs########", "..."]
}}
Include at most 12 SNPs (strongest evidence first) so the JSON always ends with ] and }} — no markdown fences, no trailing commas.
{SNP_JSON_RULES}
"""
            try:
                content_clean = call(system_prompt, prompt)
            except Exception as e:
                print(f"Warning: API call failed SNP '{phenotype}' run {run_number} model={model_label}: {e}")
                continue
            try:
                data = json.loads(_strip_json_markdown(content_clean))
            except json.JSONDecodeError:
                data = None
            if not isinstance(data, dict):
                print(f"Warning: JSON parse fail SNP '{phenotype}' run {run_number} model={model_label}")
                continue
            data["model"] = model_label
            data["run_number"] = run_number
            data["phenotype"] = phenotype
            snps = data.get("SNP", [])
            if isinstance(snps, str):
                snps = [snps]
            if not isinstance(snps, list):
                print(f"Warning: SNP field not a list for '{phenotype}' model={model_label}")
                continue
            for snp in snps:
                if _is_missing_item(snp):
                    continue
                dest.append(
                    {
                        "model": data["model"],
                        "phenotype": data["phenotype"],
                        "SNP": snp,
                        "run_number": data["run_number"],
                    }
                )


def _accum_rows_phenotype_gene(
    call, system_prompt, model_label, phenotypes, dest: list, *, n_runs: int = 1, max_items: Optional[int] = None
) -> None:
    for phenotype in _slice_items(phenotypes, max_items):
        for run_number in range(1, n_runs + 1):
            prompt = f"""What gene or genes are associated with {phenotype}?
Return a single JSON object with keys "phenotype" and "Gene" only. Example shape:
{{
  "phenotype": "{phenotype}",
  "Gene": ["GENE1", "GENE2", "..."]
}}
{GENE_JSON_RULES}
"""
            try:
                content_clean = call(system_prompt, prompt)
            except Exception as e:
                print(f"Warning: API call failed Gene '{phenotype}' run {run_number} model={model_label}: {e}")
                continue
            try:
                data = json.loads(_strip_json_markdown(content_clean))
            except json.JSONDecodeError:
                print(f"JSON parse fail Gene '{phenotype}' run {run_number} model={model_label}")
                continue
            data["model"] = model_label
            data["run_number"] = run_number
            data["phenotype"] = phenotype
            for gene in data.get("Gene", []):
                if _is_missing_item(gene):
                    continue
                dest.append(
                    {
                        "model": data["model"],
                        "phenotype": data["phenotype"],
                        "Gene": gene,
                        "run_number": data["run_number"],
                    }
                )


def _accum_rows_gene_to_phenotype(
    call, model_label, dest: list, *, n_runs: int = 1, max_items: Optional[int] = None
) -> None:
    system = SYSTEM_PROMPT_COMMON
    for gene in _slice_items(GENES_G2P, max_items):
        for run_number in range(1, n_runs + 1):
            prompt = f"""What phenotype is associated with the gene {gene}?
Return a single JSON object with keys "Gene" and "phenotype" only. Example shape:
{{
  "Gene": "{gene}",
  "phenotype": ["...", "..."]
}}
{PHENO_JSON_RULES}
"""
            try:
                content_clean = call(system, prompt)
            except Exception as e:
                print(f"Warning: API call failed G2P '{gene}' run {run_number} model={model_label}: {e}")
                continue
            try:
                data = json.loads(_strip_json_markdown(content_clean))
            except json.JSONDecodeError:
                print(f"JSON parse fail G2P '{gene}' run {run_number} model={model_label}")
                continue
            data["model"] = model_label
            data["run_number"] = run_number
            data["Gene"] = gene
            for phenotype in data.get("phenotype", []):
                if _is_missing_item(phenotype):
                    continue
                dest.append(
                    {
                        "model": data["model"],
                        "Gene": data["Gene"],
                        "phenotype": phenotype,
                        "run_number": data["run_number"],
                    }
                )


def _accum_rows_snp_to_phenotype(
    call, model_label, dest: list, *, n_runs: int = 1, max_items: Optional[int] = None
) -> None:
    system = SYSTEM_PROMPT_COMMON
    for snp in _slice_items(SNPS_P2P, max_items):
        for run_number in range(1, n_runs + 1):
            prompt = f"""What phenotype is associated with SNP {snp}?
Return a single JSON object with keys "SNP" and "phenotype" only. Example shape:
{{
  "SNP": "{snp}",
  "phenotype": ["...", "..."]
}}
Include at most 25 phenotype items (strongest evidence first) so the JSON always ends cleanly with ] and }}.
No markdown fences, no comments, no trailing commas.
{PHENO_JSON_RULES}
"""
            try:
                content_clean = call(system, prompt)
            except Exception as e:
                print(f"Warning: API call failed SNP2P '{snp}' run {run_number} model={model_label}: {e}")
                continue
            try:
                data = json.loads(_strip_json_markdown(content_clean))
            except json.JSONDecodeError:
                print(f"JSON parse fail SNP2P '{snp}' run {run_number} model={model_label}")
                continue
            data["model"] = model_label
            data["run_number"] = run_number
            data["SNP"] = snp
            for phenotype in data.get("phenotype", []):
                if _is_missing_item(phenotype):
                    continue
                dest.append(
                    {
                        "model": data["model"],
                        "SNP": data["SNP"],
                        "phenotype": phenotype,
                        "run_number": data["run_number"],
                    }
                )


def run_one_model(cfg: dict, buckets: dict) -> tuple:
    # Resolve task keys, call model for each task, optionally flush CSV per task.
    if SANITY_CHECK_TASK_KEYS is not None:
        for k in SANITY_CHECK_TASK_KEYS:
            if k not in SANITY_TASK_KEYS:
                raise ValueError(
                    f"SANITY_CHECK_TASK_KEYS must only contain keys from {sorted(SANITY_TASK_KEYS)}, got {k!r}"
                )
        task_keys = tuple(SANITY_CHECK_TASK_KEYS)
    elif SANITY_CHECK_TASK is None:
        task_keys = tuple(SANITY_TASK_KEYS)
    else:
        if SANITY_CHECK_TASK not in SANITY_TASK_KEYS:
            raise ValueError(
                f"SANITY_CHECK_TASK must be None or one of {sorted(SANITY_TASK_KEYS)}, got {SANITY_CHECK_TASK!r}"
            )
        task_keys = (SANITY_CHECK_TASK,)

    if _cfg_is_openai_gpt55(cfg):
        validate_openai_gpt55_format()
        if OPENAI_GPT55_USE_SPLIT_TASK_KEYS and OPENAI_GPT55_TASK_KEYS:
            for k in OPENAI_GPT55_TASK_KEYS:
                if k not in SANITY_TASK_KEYS:
                    raise ValueError(
                        f"OPENAI_GPT55_TASK_KEYS must only contain keys from {sorted(SANITY_TASK_KEYS)}, got {k!r}"
                    )
            task_keys = tuple(OPENAI_GPT55_TASK_KEYS)

    call = make_caller(cfg["provider"], cfg["model"])
    label = cfg["csv_model"]
    nr, mi = N_RUNS, None

    for tk in task_keys:
        if tk == "phenotype_to_snp_common":
            _accum_rows_phenotype_snp(
                call, SYSTEM_PROMPT_COMMON, label, PHENOTYPES_COMMON, buckets["phenotype_to_snp_common"], n_runs=nr, max_items=mi
            )
        elif tk == "phenotype_to_snp_rare":
            _accum_rows_phenotype_snp(
                call, SYSTEM_PROMPT_RARE, label, PHENOTYPES_RARE, buckets["phenotype_to_snp_rare"], n_runs=nr, max_items=mi
            )
        elif tk == "phenotype_to_gene_common":
            _accum_rows_phenotype_gene(
                call, SYSTEM_PROMPT_COMMON, label, PHENOTYPES_COMMON, buckets["phenotype_to_gene_common"], n_runs=nr, max_items=mi
            )
        elif tk == "phenotype_to_gene_rare":
            _accum_rows_phenotype_gene(
                call, SYSTEM_PROMPT_RARE, label, PHENOTYPES_RARE, buckets["phenotype_to_gene_rare"], n_runs=nr, max_items=mi
            )
        elif tk == "gene_to_phenotype":
            _accum_rows_gene_to_phenotype(call, label, buckets["gene_to_phenotype"], n_runs=nr, max_items=mi)
        elif tk == "SNP_to_phenotype":
            _accum_rows_snp_to_phenotype(call, label, buckets["SNP_to_phenotype"], n_runs=nr, max_items=mi)
        if FLUSH_CSV_AFTER_EACH_TASK:
            write_result_csv_for_task(
                buckets,
                tk,
                model_label=label,
                reasoning_level=_export_reasoning_level(cfg),
                only_model_rows=True,
            )

    return task_keys


def empty_buckets():
    return {key: [] for key in SANITY_TASK_KEYS}


# --- CSV export ---
# task_key -> (legacy filename, columns)
CSV_EXPORT_SPECS: dict[str, tuple[str, list[str]]] = {
    "phenotype_to_snp_common": ("phenotype_to_snp_common.csv", ["model", "phenotype", "SNP", "run_number"]),
    "phenotype_to_snp_rare": ("phenotype_to_snp_rare.csv", ["model", "phenotype", "SNP", "run_number"]),
    "phenotype_to_gene_common": ("phenotype_to_gene_common.csv", ["model", "phenotype", "Gene", "run_number"]),
    "phenotype_to_gene_rare": ("phenotype_to_gene_rare.csv", ["model", "phenotype", "Gene", "run_number"]),
    "gene_to_phenotype": ("gene_to_phenotype.csv", ["model", "Gene", "phenotype", "run_number"]),
    "SNP_to_phenotype": ("SNP_to_phenotype.csv", ["model", "SNP", "phenotype", "run_number"]),
}

# Category slug used in model-specific CSV names.
TASK_KEY_TO_CATEGORY_SLUG: dict[str, str] = {
    "phenotype_to_snp_common": "phenotype-snp-common",
    "phenotype_to_snp_rare": "phenotype-snp-rare",
    "phenotype_to_gene_common": "phenotype-gene-common",
    "phenotype_to_gene_rare": "phenotype-gene-rare",
    "gene_to_phenotype": "gene-phenotype",
    "SNP_to_phenotype": "snp-phenotype",
}


def _safe_model_tag(model_label: str) -> str:
    return (
        model_label.strip().lower().replace(" ", "_").replace("-", "_").replace(".", "_")
    )


def _safe_reasoning_tag(level: str) -> str:
    return level.strip().lower().replace(" ", "_").replace(".", "_")


def write_result_csv_for_task(
    buckets: dict,
    task_key: str,
    *,
    model_label: Optional[str] = None,
    reasoning_level: Optional[str] = None,
    only_model_rows: bool = False,
) -> str:
    # Write one task bucket to CSV.
    if task_key not in CSV_EXPORT_SPECS:
        raise ValueError(f"Unknown task_key: {task_key!r}")

    fname, fields = CSV_EXPORT_SPECS[task_key]
    if model_label:
        cat = TASK_KEY_TO_CATEGORY_SLUG[task_key]
        rtag = _safe_reasoning_tag(reasoning_level or "none")
        out_path = f"{OUT_PREFIX}{_safe_model_tag(model_label)}_{rtag}_{cat}.csv"
    else:
        out_path = OUT_PREFIX + fname

    rows = buckets.get(task_key, [])
    if only_model_rows and model_label is not None:
        rows = [r for r in rows if r.get("model") == model_label]

    with open(out_path, mode="w", newline="") as file:
        w = csv.DictWriter(file, fieldnames=fields)
        w.writeheader()
        w.writerows(rows)
    print(f"Saved {len(rows)} rows to {out_path!r}")
    return out_path


def write_result_csvs(
    buckets: dict,
    *,
    model_label: Optional[str] = None,
    reasoning_level: Optional[str] = None,
    only_model_rows: bool = False,
) -> None:
    for task_key in CSV_EXPORT_SPECS:
        write_result_csv_for_task(
            buckets,
            task_key,
            model_label=model_label,
            reasoning_level=reasoning_level,
            only_model_rows=only_model_rows,
        )


def run_one_model_and_write_csvs(cfg: dict, buckets: dict) -> None:
    # Run one model, then write CSVs (all tasks unless EXPORT_CSV_TASK_KEYS is set).
    print(f"CSV directory: {os.path.abspath(os.getcwd())}", flush=True)
    executed_task_keys = run_one_model(cfg, buckets)
    label = cfg["csv_model"]
    rlevel = _export_reasoning_level(cfg)
    keys = EXPORT_CSV_TASK_KEYS
    # Per-task flush already wrote these CSVs; skip duplicate final write unless EXPORT_CSV_TASK_KEYS pins a subset.
    if FLUSH_CSV_AFTER_EACH_TASK and keys is None:
        return
    if keys is not None:
        for tk in keys:
            write_result_csv_for_task(
                buckets,
                tk,
                model_label=label,
                reasoning_level=rlevel,
                only_model_rows=True,
            )
    elif _cfg_is_openai_gpt55(cfg) and OPENAI_GPT55_USE_SPLIT_TASK_KEYS and OPENAI_GPT55_TASK_KEYS:
        for tk in executed_task_keys:
            write_result_csv_for_task(
                buckets,
                tk,
                model_label=label,
                reasoning_level=rlevel,
                only_model_rows=True,
            )
    else:
        write_result_csvs(
            buckets,
            model_label=label,
            reasoning_level=rlevel,
            only_model_rows=True,
        )

# %% [code]
# API keys — set in the environment before running (do not hardcode keys here).
import os

os.environ.setdefault("OPENAI_API_KEY", "")
os.environ.setdefault("ANTHROPIC_API_KEY", "")
os.environ.setdefault("GOOGLE_API_KEY", "")
os.environ.setdefault("DEEPSEEK_API_KEY", "")
# Fill in above locally, or export in shell before running.

_REQUIRED_KEYS = ("OPENAI_API_KEY", "ANTHROPIC_API_KEY", "GOOGLE_API_KEY", "DEEPSEEK_API_KEY")
_missing = [name for name in _REQUIRED_KEYS if not os.environ.get(name, "").strip()]
for name in _REQUIRED_KEYS:
    print(f"{name}: {'OK' if name not in _missing else 'MISSING'}")
if _missing:
    raise RuntimeError(
        "Missing API keys in environment: "
        + ", ".join(_missing)
        + ". Export them in your shell before running this notebook."
    )


# %% [markdown]
### Four models · CSV outputs

**Models:** `gpt-5.5`, `claude-sonnet-4-6`, `gemini-3-flash-preview`, `deepseek-v4-flash`.

**API keys:** set in the environment (`OPENAI_API_KEY`, `ANTHROPIC_API_KEY`, `GOOGLE_API_KEY` or `GEMINI_API_KEY`, `DEEPSEEK_API_KEY`). Do not hardcode keys in the notebook.

**Workflow:** run setup → `buckets = empty_buckets()` → run each model cell, or call `run_all_models_and_write()`.

**Tasks:** six categories (common/rare phenotype→SNP, phenotype→gene, gene→phenotype, SNP→phenotype). Default runs all six; use `SANITY_CHECK_TASK` or `SANITY_CHECK_TASK_KEYS` to subset.

**Output:** `{model}_{reasoning}_{category}.csv` (see `TASK_KEY_TO_CATEGORY_SLUG`). Re-run setup after changing task/export settings.


# %% [code]
# Reset result buckets for a fresh run.
buckets = empty_buckets()


# %% [markdown]
## 1. OpenAI · gpt-5.5


# %% [code]
# Model 1/4
run_one_model_and_write_csvs(MODEL_CONFIGS[0], buckets)
print('done:', MODEL_CONFIGS[0]['csv_model'])


# %% [markdown]
## 2. Anthropic · claude-sonnet-4-6


# %% [code]
# Model 2/4
run_one_model_and_write_csvs(MODEL_CONFIGS[1], buckets)
print('done:', MODEL_CONFIGS[1]['csv_model'])


# %% [markdown]
## 3. Google · gemini-3-flash-preview


# %% [code]
# Model 3/4
run_one_model_and_write_csvs(MODEL_CONFIGS[2], buckets)
print('done:', MODEL_CONFIGS[2]['csv_model'])


# %% [markdown]
## 4. DeepSeek · deepseek-v4-flash


# %% [code]
# Model 4/4
run_one_model_and_write_csvs(MODEL_CONFIGS[3], buckets)
print('done:', MODEL_CONFIGS[3]['csv_model'])
````
