## Supplementary material for "Assessing the Reliability of LLM-Generated Phenotype-Genotype Associations Through External Validation": generate_euler.py

```
#!/usr/bin/env python3
# Builds the 6-panel Euler diagram (Figure 6). Layout comes from the real
# nVennR R package (nvenn_render_panel.R); this script preps the per-panel
# entity sets, calls it, and checks every region count against pandas
# before compositing the combined figure.
import json
import subprocess
import sys
from collections import defaultdict
from pathlib import Path

import pandas as pd
from PIL import Image, ImageDraw, ImageFont

HERE      = Path(__file__).resolve().parent
GENE_FILE = HERE.parent / "phenotype_gene_verified_combined_by_api_with_gwas_omim_with_3_rate.xlsx"
SNP_FILE  = HERE.parent / "phenotype_snp_verified_combined_by_api_with_gwas_omim_with_3_rate.xlsx"
OUT_DIR   = HERE / "euler_nvenn_panels"
R_SCRIPT  = HERE / "nvenn_render_panel.R"
N_CYCLES  = 7000

TIMES_REGULAR = "/System/Library/Fonts/Supplemental/Times New Roman.ttf"
TIMES_BOLD    = "/System/Library/Fonts/Supplemental/Times New Roman Bold.ttf"

MODEL_ORDER = ["gpt-5.5", "gemini-3-flash-preview", "claude-sonnet-4-6", "deepseek-v4-flash"]
MODEL_COLORS = {
    "gpt-5.5": "#F48FB1",
    "gemini-3-flash-preview": "#A5D6A7",
    "claude-sonnet-4-6": "#FFF59D",
    "deepseek-v4-flash": "#90CAF9",
}

TASK_ORDER = ["phenotype-gene-common", "phenotype-gene-rare", "gene-phenotype",
              "phenotype-snp-common",  "phenotype-snp-rare",  "snp-phenotype"]
TASK_LABELS = ["A. Phenotype-gene common", "B. Phenotype-gene rare", "C. Gene-phenotype",
               "D. Phenotype-SNP common",  "E. Phenotype-SNP rare",  "F. SNP-phenotype"]

OUT_DIR.mkdir(exist_ok=True)


def clean(v):
    # pd.isna() check matters here: `v or ""` treats NaN as truthy and
    # would collapse every SNP row into one fake "phenotype||nan" entity.
    if pd.isna(v):
        return ""
    return str(v).strip().lower()


def make_key(row):
    p = clean(row.get("Phenotype"))
    g = clean(row.get("Gene"))
    s = clean(row.get("SNP"))
    return p + "||" + (g if g else s)


print("Loading data...")
gene_df = pd.read_excel(GENE_FILE)
snp_df = pd.read_excel(SNP_FILE)
if "SNP" not in gene_df.columns:
    gene_df["SNP"] = None
if "Gene" not in snp_df.columns:
    snp_df["Gene"] = None

all_df = pd.concat([gene_df, snp_df], ignore_index=True)
all_df["Model"] = all_df["Model"].astype(str).str.strip()
all_df["Source_Sheet"] = all_df["Source_Sheet"].astype(str).str.strip()
all_df["_key"] = all_df.apply(make_key, axis=1)
all_df = all_df[all_df["_key"] != "||"].copy()


def get_sets(task):
    sub = all_df[all_df["Source_Sheet"] == task]
    return {m: set(sub.loc[sub["Model"] == m, "_key"]) for m in MODEL_ORDER}


def ground_truth_regions(sets):
    """{frozenset(model_names): count}, computed directly from the raw
    entity sets -- independent of any plotting/layout step."""
    member = defaultdict(set)
    for m, ents in sets.items():
        for e in ents:
            member[e].add(m)
    counts = defaultdict(int)
    for ms in member.values():
        if ms:
            counts[frozenset(ms)] += 1
    return dict(counts)


verification_lines = []
panel_pngs = []
panel_svgs = []
all_ok = True

for task, label in zip(TASK_ORDER, TASK_LABELS):
    letter = label[0]
    print(f"\n=== Panel {letter}: {task} ===")
    sets = get_sets(task)
    active_models = [m for m in MODEL_ORDER if len(sets[m]) > 0]
    truth = ground_truth_regions(sets)

    in_json = OUT_DIR / f"panel_{letter}_input.json"
    out_svg = OUT_DIR / f"panel_{letter}.svg"
    out_counts = OUT_DIR / f"panel_{letter}_counts.json"

    json.dump({m: sorted(sets[m]) for m in active_models}, open(in_json, "w"))

    result = subprocess.run(
        ["Rscript", str(R_SCRIPT), str(in_json), str(out_svg), str(out_counts), str(N_CYCLES)],
        capture_output=True, text=True,
    )
    if result.returncode != 0:
        raise RuntimeError(f"R rendering failed for panel {letter}:\n{result.stdout}\n{result.stderr}")
    print(result.stdout.strip())

    r_counts_raw = json.load(open(out_counts))
    r_counts = {}
    for region_str, cnt in r_counts_raw.items():
        models_in_region = frozenset(x.strip() for x in region_str.split(","))
        r_counts[models_in_region] = cnt

    mismatches = []
    all_keys = set(truth.keys()) | set(r_counts.keys())
    for key in all_keys:
        t = truth.get(key, 0)
        r = r_counts.get(key, 0)
        if t != r:
            mismatches.append((sorted(key), t, r))

    verification_lines.append(f"Panel {letter} ({task}):")
    for key in sorted(truth.keys(), key=lambda k: -truth[k]):
        models_str = ", ".join(sorted(key))
        verification_lines.append(f"  [{models_str}]  pandas={truth[key]}  nVennR={r_counts.get(key, 'MISSING')}")
    if mismatches:
        verification_lines.append(f"  !!! MISMATCH: {mismatches}")
        all_ok = False
    else:
        verification_lines.append(f"  OK: all {len(truth)} regions match exactly.")
    verification_lines.append("")

    panel_pngs.append(OUT_DIR / f"panel_{letter}.png")
    panel_svgs.append(out_svg)

verification_txt = "\n".join(verification_lines)
(OUT_DIR / "verification.txt").write_text(verification_txt)
print("\n" + verification_txt)

if not all_ok:
    raise RuntimeError(
        "Region-count verification FAILED for at least one panel -- see "
        "euler_nvenn_panels/verification.txt for details. Refusing to build "
        "a combined figure from unverified data."
    )

print("All panels verified against independent pandas computation (exact match). Building combined figure...")


# combined PNG: 2x3 grid + shared legend
TITLE_H = 56
imgs = [Image.open(p) for p in panel_pngs]
cell_w = max(im.width for im in imgs)
cell_h = max(im.height for im in imgs) + TITLE_H
ncols, nrows = 3, 2

try:
    font_title = ImageFont.truetype(TIMES_BOLD, 36)
    font_leg_title = ImageFont.truetype(TIMES_BOLD, 66)
    font_leg = ImageFont.truetype(TIMES_REGULAR, 52)
except Exception:
    font_title = font_leg_title = font_leg = ImageFont.load_default()

legend_w = 1080
canvas_w = cell_w * ncols + legend_w
canvas_h = cell_h * nrows
canvas = Image.new("RGB", (canvas_w, canvas_h), "white")
draw = ImageDraw.Draw(canvas)

for idx, (im, label) in enumerate(zip(imgs, TASK_LABELS)):
    r, c = divmod(idx, ncols)
    x = c * cell_w
    y = r * cell_h
    draw.text((x + 18, y + 8), label, fill="black", font=font_title)
    canvas.paste(im, (x + (cell_w - im.width) // 2, y + TITLE_H))

leg_x = cell_w * ncols + 110
leg_y = canvas_h // 2 - 200
draw.rectangle(
    [leg_x - 35, leg_y - 100, leg_x + 900, leg_y + 100 + 92 * len(MODEL_ORDER)],
    outline="black", width=4,
)
draw.text((leg_x, leg_y - 75), "Model legend", fill="black", font=font_leg_title)
for i, m in enumerate(MODEL_ORDER):
    yy = leg_y + 30 + i * 92
    draw.rectangle([leg_x, yy, leg_x + 74, yy + 56], fill=MODEL_COLORS[m], outline="black", width=3)
    draw.text((leg_x + 96, yy + 6), f"{i + 1} = {m}", fill="black", font=font_leg)

combined_png = OUT_DIR / "combined_figure.png"
canvas.save(combined_png, dpi=(300, 300))
print("Saved:", combined_png)


# combined SVG (best-effort; PNG above is the primary deliverable)
try:
    import svgutils.transform as sg

    panel_figs = [sg.fromfile(str(p)) for p in panel_svgs]
    panel_w = float(str(panel_figs[0].width).replace("px", "") or 700)
    panel_h = float(str(panel_figs[0].height).replace("px", "") or 500)
    title_h_svg = 30
    legend_w_svg = 260
    scale = 0.85

    fig = sg.SVGFigure(
        f"{(panel_w * scale + 10) * 3 + legend_w_svg}px",
        f"{(panel_h * scale + title_h_svg + 10) * 2}px",
    )
    elements = []
    for idx, (pf, label) in enumerate(zip(panel_figs, TASK_LABELS)):
        r, c = divmod(idx, 3)
        x = c * (panel_w * scale + 10)
        y = r * (panel_h * scale + title_h_svg + 10)
        root = pf.getroot()
        root.moveto(x, y + title_h_svg, scale_x=scale, scale_y=scale)
        elements.append(root)
        title = sg.TextElement(x + 5, y + title_h_svg - 8, label, size=15, weight="bold", font="Times New Roman")
        elements.append(title)

    leg_x0 = 3 * (panel_w * scale + 10) + 10
    leg_y0 = 40
    elements.append(sg.TextElement(leg_x0, leg_y0, "Model legend", size=22, weight="bold", font="Times New Roman"))
    for i, m in enumerate(MODEL_ORDER):
        yy = leg_y0 + 40 + i * 42
        sq = sg.fromstring(
            f'<rect x="{leg_x0}" y="{yy}" width="30" height="22" '
            f'fill="{MODEL_COLORS[m]}" stroke="black" stroke-width="1"/>'
        ).getroot()
        elements.append(sq)
        elements.append(sg.TextElement(leg_x0 + 40, yy + 17, f"{i + 1} = {m}", size=18, font="Times New Roman"))

    fig.append(elements)
    combined_svg = OUT_DIR / "combined_figure.svg"
    fig.save(str(combined_svg))
    print("Saved:", combined_svg)
except Exception as e:
    print(f"WARNING: combined SVG assembly failed ({e}); combined PNG is still available.", file=sys.stderr)

print("\nDone.")
```
