## Supplementary material for "Assessing the Reliability of LLM-Generated Phenotype-Genotype Associations Through External Validation": LLM_CODE_ai_verified_final.ipynb

```
# %% [markdown]
# LLM phenotype–gene/SNP verification pipeline


# %% [markdown]
## 1. Configuration


# %% [code]
# Core imports and plotting defaults
import json
import sys
import os
import random
import re
import time
from itertools import product
from pathlib import Path

import matplotlib.pyplot as plt
import pandas as pd
import requests
import seaborn as sns

# Locate the project root (the directory containing both pipeline/ and
# config.yaml) regardless of where this notebook's kernel was launched from
# -- this notebook lives in JAMIA_abstract_all_file/, one level below the
# project root where the `pipeline` package actually is.
def _find_project_root(start: Path, markers=("pipeline", "config.yaml")) -> Path:
    for candidate in [start, *start.parents]:
        if all((candidate / m).exists() for m in markers):
            return candidate
    raise RuntimeError(
        f"Could not locate project root (looked for pipeline/ and config.yaml) "
        f"starting from {start}. Run this notebook from within the project, "
        f"or adjust markers in _find_project_root()."
    )

_PROJECT_ROOT = _find_project_root(Path.cwd())
if str(_PROJECT_ROOT) not in sys.path:
    sys.path.insert(0, str(_PROJECT_ROOT))

from pipeline.config import apply_config_to_globals, load_config
from pipeline.plot_style import apply_transparent_figure, use_transparent_plots

use_transparent_plots()

# %% [code]
# Load config.yaml
_cfg = load_config()
apply_config_to_globals(_cfg, globals())


# %% [markdown]
## 2. Data preparation, verification & analytics

Run the cells below **top to bottom**: load CSVs → verify against GWAS/OMIM → tables & figures.

### Load CSV and export workbooks


# %% [code]
def col_key(raw):
    """Lowercase key: spaces, underscores, hyphens collapsed to single underscores."""
    s = str(raw).strip().lstrip("\ufeff").lower()
    s = s.replace("_", " ").replace("-", " ")
    parts = [p for p in s.split() if p]
    return "_".join(parts)

def normalize_columns(df):
    CANON = {
        "model": "Model",
        "phenotype": "Phenotype",
        "pheno": "Phenotype",
        "trait": "Phenotype",
        "disease": "Phenotype",
        "gene": "Gene",
        "gene_symbol": "Gene",
        "genesymbol": "Gene",
        "symbol": "Gene",
        "snp": "SNP",
        "run_number": "Run Number",
        "run": "Run Number",
        "run_no": "Run Number",
        "runnr": "Run Number",
        "verification": "Verification",
        "verified": "Verification",
        "verify": "Verification",
        "note": "Note",
        "notes": "Note",
        "comment": "Note",
        "links": "Links",
        "link": "Links",
        "url": "Links",
        "gwas_phenotype": "GWAS_Phenotype",
        "study_accession": "Study_Accession",
        "sample": "Sample",
    }

    mapping = {}
    for c in df.columns:
        raw = str(c).strip()
        k = col_key(raw)
        if k in CANON:
            mapping[c] = CANON[k]
        else:
            mapping[c] = raw

    return df.rename(columns=mapping)


# %% [code]
def load_csv_task(path):
    df = pd.read_csv(path)
    df = normalize_columns(df)
    stem = Path(path).stem
    virtual_sheet = stem.split("_default_", 1)[1] if "_default_" in stem else stem
    df["Source_File"] = str(path)
    df["Source_Sheet"] = virtual_sheet
    return df

combined_df = pd.concat([load_csv_task(f) for f in FILES], ignore_index=True)
_allowed = {m.lower() for m in ALLOWED_MODELS}
combined_df = combined_df[
    combined_df["Model"].astype(str).str.strip().str.lower().isin(_allowed)
].copy()


# %% [code]
GENE_SOURCE_SHEETS = frozenset(
    {"gene-phenotype", "phenotype-gene-common", "phenotype-gene-rare"}
)
SNP_SOURCE_SHEETS = frozenset(
    {"phenotype-snp-common", "phenotype-snp-rare", "snp-phenotype"}
)

if "combined_df" not in dir() or combined_df is None or len(combined_df) == 0:
    raise RuntimeError("Please run the CSV-loading cells above first so that combined_df exists and is non-empty.")
if "Source_Sheet" not in combined_df.columns or "Model" not in combined_df.columns:
    raise RuntimeError("combined_df must contain Model and Source_Sheet columns (from load_csv_task).")

gene_df = combined_df[combined_df["Source_Sheet"].astype(str).isin(GENE_SOURCE_SHEETS)].copy()
snp_df = combined_df[combined_df["Source_Sheet"].astype(str).isin(SNP_SOURCE_SHEETS)].copy()
_other = combined_df[~combined_df["Source_Sheet"].astype(str).isin(GENE_SOURCE_SHEETS | SNP_SOURCE_SHEETS)]
if len(_other):
    print("Unmapped Source_Sheet values (not in gene/snp groups):", _other["Source_Sheet"].unique()[:15])

gene_final = gene_df.reset_index(drop=True)
snp_final = snp_df.reset_index(drop=True)

gene_final.to_excel(
    GENE_INPUT_XLSX,
    index=False
)

snp_final.to_excel(
    SNP_INPUT_XLSX,
    index=False
)

print("Tables generated:")
print(GENE_INPUT_XLSX)
print(SNP_INPUT_XLSX)
print("Gene relations:", len(gene_final))
print("SNP relations:", len(snp_final))
print("Gene by Source_Sheet:\n", gene_final["Source_Sheet"].value_counts().to_string())
print("SNP by Source_Sheet:\n", snp_final["Source_Sheet"].value_counts().to_string())


# %% [code]
try:
    _ALLOWED_MODEL_ORDER = list(ALLOWED_MODELS)
    _ALLOWED_MODEL_SET = {m.lower() for m in ALLOWED_MODELS}
except NameError:
    _ALLOWED_MODEL_ORDER = ["claude-sonnet-4-6", "deepseek-v4-flash", "gemini-3-flash-preview", "gpt-5.5"]
    _ALLOWED_MODEL_SET = {m.lower() for m in _ALLOWED_MODEL_ORDER}

_meta = ["Model", "Source_Sheet"]
_GENE_TASKS = frozenset({"gene-phenotype", "phenotype-gene-common", "phenotype-gene-rare"})
_verbose = globals().get("VERBOSE", False)

def df_for_generated_stats():
    if "combined_df" in dir():
        cdf = combined_df
        if cdf is not None and all(c in cdf.columns for c in _meta):
            if _verbose:
                print("  Source_Sheet values:", sorted(cdf["Source_Sheet"].astype(str).unique()))
            return cdf
    parts = []
    for p in (GENE_INPUT_XLSX, SNP_INPUT_XLSX):
        try:
            parts.append(pd.read_excel(p))
        except Exception:
            pass
    if not parts:
        raise FileNotFoundError(
            "combined_df is not available in memory, and phenotype_*_verified_combined_by_api.xlsx could not be read; please run the earlier load/export cells first."
        )
    return pd.concat(parts, ignore_index=True)

_stats_df = df_for_generated_stats()

# Quick check: do gene-task rows survive the model filter?
if {"Model", "Source_Sheet"}.issubset(_stats_df.columns):
    _gmask = _stats_df["Source_Sheet"].astype(str).isin(_GENE_TASKS)
    if _gmask.any():
        _pass = _stats_df.loc[_gmask, "Model"].astype(str).str.strip().str.lower().isin(_ALLOWED_MODEL_SET)
        if int(_pass.sum()) == 0:
            print(
                "  In the gene table, Model values do not match ALLOWED_MODELS; after filtering, gene-task counts become 0."
                "  Please inspect the Model column in phenotype_gene_verified_combined_by_api.xlsx, or rerun the Section 3 export cell to overwrite that file using the current combined_df."
            )
            if _verbose:
                print("  Sample Model values in gene-task rows:", _stats_df.loc[_gmask, "Model"].astype(str).unique()[:12])

_stats_df = _stats_df[_stats_df["Model"].astype(str).str.strip().str.lower().isin(_ALLOWED_MODEL_SET)].copy()
if not all(c in _stats_df.columns for c in _meta):
    raise ValueError(f"Missing required columns {_meta}; current columns: {list(_stats_df.columns)}")

_task_order = list(TASK_ORDER_SIX)
_seen_tasks = sorted(_stats_df["Source_Sheet"].astype(str).unique())
if len(_seen_tasks) < len(_task_order):
    miss = [t for t in _task_order if t not in _seen_tasks]
    print(
        "  Note: only a subset of tasks has rows in current data, so only those tasks appear in this summary. "
        "Common reasons: (1) in-memory combined_df contains only SNP-related sheets; "
        "(2) an export/xlsx branch is missing or empty, so fewer than six task types appear after load. "
        f"Missing tasks: {miss}"
    )

generated_counts = (
    _stats_df.groupby(_meta, dropna=False).size().reset_index(name="Generated_Count").sort_values(_meta)
)
_models_plot = [m for m in _ALLOWED_MODEL_ORDER if m in set(_stats_df["Model"].astype(str))]
generated_counts_full = (
    pd.DataFrame(list(product(_models_plot, _task_order)), columns=_meta)
    .merge(generated_counts, on=_meta, how="left")
)
generated_counts_full["Generated_Count"] = generated_counts_full["Generated_Count"].fillna(0).astype(int)

generated_pivot = (
    generated_counts_full.pivot(index="Model", columns="Source_Sheet", values="Generated_Count")
    .reindex(index=_models_plot, columns=_task_order, fill_value=0)
    .astype(int)
)

print("\nLong table (non-zero combinations only):")
print(generated_counts.to_string(index=False))
print("\nLong table (full grid: 6 tasks × models, missing=0):")
print(generated_counts_full.to_string(index=False))
print("\nPivot table (rows=Model, columns=Task, fixed 6 columns):")
print(generated_pivot.to_string())
print("\nTotal rows in source dataframe:", len(_stats_df))


# %% [markdown]
### Verification (GWAS + OMIM)


# %% [code]
# Load GWAS Catalog associations 
from pathlib import Path

_gwas_path = Path(GWAS_CATALOG_TSV)
if not _gwas_path.is_file():
    raise FileNotFoundError(
        f"GWAS catalog not found: {_gwas_path}\n"
        f"Download from: {GWAS_CATALOG_URL or 'config key gwas_catalog_url'}\n"
        "Save as gwas_catalog_tsv in config.yaml."
    )

gwas = pd.read_csv(_gwas_path, sep="\t", low_memory=False)
gwas.columns = gwas.columns.str.strip().str.lower()

TEXT_COLS = ["mapped_gene", "reported gene(s)", "snps", "disease/trait"]
for col in TEXT_COLS:
    if col in gwas.columns:
        gwas[col] = gwas[col].astype(str).str.lower().str.strip()

SYNONYMS = {
    "carcinoma": "cancer",
    "tumor": "cancer",
    "tumour": "cancer",
    "neoplasm": "cancer",
}
MODIFIER_TERMS = ["familial", "susceptibility", "type", "syndrome", "disease"]

def normalize_pheno_with_details(text):
    original = str(text or "")
    lowered = original.lower()
    work = lowered
    removed = []
    for w in MODIFIER_TERMS:
        if w in work:
            count = work.count(w)
            if count > 0:
                removed.extend([w] * count)
                work = work.replace(w, " ")
    replacements = []
    for k, v in SYNONYMS.items():
        if k in work:
            count = work.count(k)
            if count > 0:
                replacements.extend([f"{k}->{v}"] * count)
                work = work.replace(k, v)
    normalized = " ".join(work.split()).strip()
    return {
        "lower_text": lowered,
        "normalized_text": normalized,
        "removed_modifiers": removed,
        "synonym_replacements": replacements,
    }

def normalize_pheno(text):
    return normalize_pheno_with_details(text)["normalized_text"]


# %% [code]
# Map phenotypes to EFO / MONDO ontology terms (input Phenotype only).
OLS_BASE_URL = "https://www.ebi.ac.uk/ols4"
ONTOLOGY_ORDER = ["efo", "mondo"]
OLS_CACHE = {}
PHENO_MAP_CACHE = {}
PHENO_AUDIT = {}
PHENO_MATCH_MIN_JACCARD = 0.4

def ols_search(query: str, ontology: str, rows: int = 8):
    key = (query, ontology, rows)
    if key in OLS_CACHE:
        return OLS_CACHE[key]
    url = f"{OLS_BASE_URL}/api/search"
    params = {"q": query, "ontology": ontology, "rows": rows}
    try:
        r = requests.get(url, params=params, timeout=20)
        if r.status_code != 200:
            OLS_CACHE[key] = []
            return []
        docs = (r.json().get("response") or {}).get("docs") or []
        out = []
        for d in docs:
            label = str(d.get("label") or "").strip()
            if not label:
                continue
            out.append(
                {
                    "label": label,
                    "id": d.get("obo_id") or d.get("short_form") or "",
                    "iri": d.get("iri") or "",
                    "score": float(d.get("score") or 0.0),
                }
            )
        OLS_CACHE[key] = out
        return out
    except Exception:
        OLS_CACHE[key] = []
        return []

def _pheno_token_jaccard(a: str, b: str) -> float:
    sa, sb = set(a.split()), set(b.split())
    if not sa or not sb:
        return 0.0
    return len(sa & sb) / len(sa | sb)

def _best_ols_candidate(basic: str, candidates: list):
    """Pick the candidate whose normalized label best matches `basic`.

    Returns (candidate_or_None, match_kind, match_score); match_kind is
    one of {"exact", "substring", "jaccard", "none"}. match_score is 1.0
    for exact, a length-ratio for substring, Jaccard overlap for jaccard.
    """
    best = None
    best_kind = "none"
    best_score = -1.0
    for h in candidates:
        label_norm = normalize_pheno(h["label"])
        if not label_norm:
            continue
        if label_norm == basic:
            return h, "exact", 1.0
        if basic in label_norm or label_norm in basic:
            # Prefer the tightest containment: a label that is mostly just
            # the query beats one burying it inside a long, unrelated
            # compound/subtype phrase (e.g. "hypertension" should prefer
            # "malignant hypertension" over "pulmonary arterial
            # hypertension associated with portal hypertension").
            kind = "substring"
            score = min(len(basic), len(label_norm)) / max(len(basic), len(label_norm))
        else:
            j = _pheno_token_jaccard(basic, label_norm)
            if j < PHENO_MATCH_MIN_JACCARD:
                continue
            kind, score = "jaccard", j
        if score > best_score:
            best, best_kind, best_score = h, kind, score
    return best, best_kind, (best_score if best is not None else 0.0)

def normalize_input_phenotype(text):
    """Normalize + map ONLY input Phenotype via EFO and MONDO.

    Queries both ontologies and only accepts a candidate that actually
    overlaps the input phenotype text. Never drops a row; falls back to
    the basic normalized text when no candidate passes the overlap check.
    """
    raw = str(text or "")
    norm_info = normalize_pheno_with_details(raw)
    basic = norm_info["normalized_text"]
    if basic in PHENO_MAP_CACHE:
        return PHENO_MAP_CACHE[basic]

    candidates = []
    for onto in ONTOLOGY_ORDER:
        hits = ols_search(basic, onto, rows=8)
        for h in hits[:5]:
            candidates.append({**h, "ontology": onto.upper()})

    chosen, match_kind, match_score = _best_ols_candidate(basic, candidates)

    if chosen is None:
        mapped = basic
        status = "UNMAPPED"
        onto_id = ""
        onto_label = ""
        onto_source = ""
        conf = 0.0
    else:
        mapped = normalize_pheno(chosen["label"])
        status = "MAPPED"
        onto_id = str(chosen.get("id") or "")
        onto_label = str(chosen.get("label") or "")
        onto_source = chosen.get("ontology", "")
        conf = float(match_score)

    candidate_log = " || ".join(
        f"{c['ontology']}|{c['id']}|{c['label']}|{c['score']:.3f}" for c in candidates
    )

    PHENO_MAP_CACHE[basic] = mapped
    PHENO_AUDIT[basic] = {
        "original_text": raw,
        "lower_text": norm_info["lower_text"],
        "removed_modifiers": " | ".join(norm_info["removed_modifiers"]),
        "synonym_replacements": " | ".join(norm_info["synonym_replacements"]),
        "normalized_text": basic,
        "mapped_text": mapped,
        "mapping_status": status,
        "match_kind": match_kind,
        "ontology_source": onto_source,
        "ontology_id": onto_id,
        "ontology_label": onto_label,
        "mapping_confidence": conf,
        "candidate_options": candidate_log,
    }
    return mapped

def export_phenotype_replacement_audit(path="phenotype_replacement_audit.xlsx"):
    cols = [
        "original_text",
        "lower_text",
        "removed_modifiers",
        "synonym_replacements",
        "normalized_text",
        "mapped_text",
        "mapping_status",
        "match_kind",
        "ontology_source",
        "ontology_id",
        "ontology_label",
        "mapping_confidence",
        "candidate_options",
    ]

    if not PHENO_AUDIT:
        with pd.ExcelWriter(path, engine="openpyxl") as w:
            pd.DataFrame(columns=cols).to_excel(w, sheet_name="mapping_audit", index=False)
            pd.DataFrame(columns=["modifier_term", "removed_count"]).to_excel(
                w, sheet_name="modifier_summary", index=False
            )
            pd.DataFrame(columns=["replacement", "replace_count"]).to_excel(
                w, sheet_name="synonym_summary", index=False
            )
        return

    df_a = pd.DataFrame(PHENO_AUDIT.values())
    for c in cols:
        if c not in df_a.columns:
            df_a[c] = ""
    df_a = df_a[cols]

    mod_counts = {}
    rep_counts = {}
    for mods in df_a["removed_modifiers"].fillna(""):
        for t in [x.strip() for x in str(mods).split("|") if x.strip()]:
            mod_counts[t] = mod_counts.get(t, 0) + 1
    for reps in df_a["synonym_replacements"].fillna(""):
        for t in [x.strip() for x in str(reps).split("|") if x.strip()]:
            rep_counts[t] = rep_counts.get(t, 0) + 1

    df_mod = pd.DataFrame(
        sorted(mod_counts.items(), key=lambda kv: (-kv[1], kv[0])),
        columns=["modifier_term", "removed_count"],
    )
    df_rep = pd.DataFrame(
        sorted(rep_counts.items(), key=lambda kv: (-kv[1], kv[0])),
        columns=["replacement", "replace_count"],
    )

    with pd.ExcelWriter(path, engine="openpyxl") as w:
        df_a.to_excel(w, sheet_name="mapping_audit", index=False)
        df_mod.to_excel(w, sheet_name="modifier_summary", index=False)
        df_rep.to_excel(w, sheet_name="synonym_summary", index=False)

    if VERBOSE:
        print(f" Saved phenotype mapping audit: {path}")

# %% [code]
# GWAS support tiers: Strong = strict trait match AND p < 5e-8 (per
# prompt); Weak = trait loosely related but Strong criteria not met;
# None = no SNP/gene hit or no trait-related association.

def normalize_snp_token(raw):
    """Normalize SNP ids from Excel (float tails, missing rs prefix)."""
    if raw is None or (isinstance(raw, float) and pd.isna(raw)):
        return ""
    s = str(raw).strip().lower()
    if s in ("", "nan", "none"):
        return ""
    if re.fullmatch(r"\d+\.0", s):
        s = s[:-2]
    if re.fullmatch(r"\d+", s):
        s = "rs" + s
    return s

def trait_strict_match(phenotype_norm: str, trait_norm: str) -> bool:
    """Strict phenotype–trait match (required together with p for Strong)."""
    if not phenotype_norm or not trait_norm:
        return False
    if phenotype_norm == trait_norm:
        return True
    if phenotype_norm in trait_norm or trait_norm in phenotype_norm:
        return True
    long_tokens = [t for t in phenotype_norm.split() if len(t) > 3]
    return any(t in trait_norm for t in long_tokens)

def trait_loose_match(phenotype_norm: str, trait_norm: str) -> bool:
    """Broader relatedness for counting any GWAS support."""
    if trait_strict_match(phenotype_norm, trait_norm):
        return True
    if not phenotype_norm or not trait_norm:
        return False
    a = {x for x in phenotype_norm.split() if len(x) > 2}
    b = {x for x in trait_norm.split() if len(x) > 2}
    if not a or not b:
        return False
    inter = len(a & b)
    if inter >= 1:
        return True
    return False

def resolve_pvalue_column(gwas_df):
    if "p-value" in gwas_df.columns:
        return "p-value"
    for c in gwas_df.columns:
        c0 = c.replace(" ", "").lower()
        if c0 in ("p-value", "pvalue", "p"):
            return c
    return None

def parse_pvalue(cell) -> float:
    """Parse catalog p-value; invalid -> inf (never genome-wide significant)."""
    if cell is None or (isinstance(cell, float) and pd.isna(cell)):
        return float("inf")
    s = str(cell).strip().lower()
    if s in ("", "nan", "nr", "n/a"):
        return float("inf")
    try:
        return float(s.replace(",", ""))
    except ValueError:
        return float("inf")

def verify_gwas(row, gwas_df, mode):
    phenotype = normalize_input_phenotype(row.get("Phenotype", ""))
    if phenotype == "":
        return "None", []

    p_col = resolve_pvalue_column(gwas_df)

    if mode == "gene":
        ent = row.get("Gene")
        if ent is None or pd.isna(ent):
            return "None", []
        entity = str(ent).strip().lower().replace(" ", "")
        if not entity or entity == "nan":
            return "None", []
        pat = rf"\b{re.escape(entity)}\b"
        hits = gwas_df[
            gwas_df["mapped_gene"].str.contains(pat, regex=True, na=False)
            | gwas_df["reported gene(s)"].str.contains(pat, regex=True, na=False)
        ]
    else:
        entity = normalize_snp_token(row.get("SNP"))
        if not entity:
            return "None", []
        pat = rf"\b{re.escape(entity)}\b"
        hits = gwas_df[gwas_df["snps"].str.contains(pat, regex=True, na=False)]

    if len(hits) == 0:
        return "None", []

    has_strong = False
    has_weak = False
    results = []

    for _, h in hits.iterrows():
        trait_norm = normalize_pheno(h.get("disease/trait", ""))
        if not trait_loose_match(phenotype, trait_norm):
            continue
        pval = parse_pvalue(h.get(p_col) if p_col else None)
        results.append(h.to_dict())
        if trait_strict_match(phenotype, trait_norm) and pval < 5e-8:
            has_strong = True
        else:
            has_weak = True

    if has_strong:
        return "Strong", results
    if has_weak:
        return "Weak", results
    return "None", []


# %% [code]
# Run GWAS + OMIM verification (OMIM keys in config.yaml KEY1/KEY2/KEY3)
from pipeline.runner import load_and_configure_engine, print_verification_preflight

run_verification_pipeline = load_and_configure_engine(_cfg, globals())

if VERIFICATION_SCOPE == "both":
    _modes = RUN_VERIFICATION_MODES
elif VERIFICATION_SCOPE == "gene":
    _modes = ["gene"]
elif VERIFICATION_SCOPE == "snp":
    _modes = ["snp"]
else:
    raise ValueError('VERIFICATION_SCOPE must be "gene", "snp", or "both"')

_preflight = print_verification_preflight(_cfg, globals(), modes=_modes)
if _preflight["missing_inputs"]:
    raise FileNotFoundError(
        "Verification inputs missing — run the Load CSV export cell above first."
    )

if VERBOSE:
    print(f"Starting verification (scope={VERIFICATION_SCOPE})")

run_verification_pipeline(modes=_modes, export_audit=EXPORT_AUDIT)

from pathlib import Path as _Path_check
_snp_out = _Path_check(SNP_OUTPUT_XLSX)
if not _snp_out.is_file():
    raise RuntimeError(f"SNP verification output missing: {_snp_out}")
print(f"Verification done. SNP output: {_snp_out.name}")


# %% [markdown]
### Analytics & figures


# %% [code]
# Structured tables + summary workbook (gene + snp): count/percentage
# tables plus a Triple_Category / Matched-evidence-mix plot panel.
import os
import numpy as np
import pandas as pd
import matplotlib.pyplot as plt
import seaborn as sns

use_transparent_plots(context="notebook")

from pathlib import Path
import warnings

_verified_root = Path(PROJECT_DIR) if "PROJECT_DIR" in globals() else Path.cwd()
if not _verified_root.is_dir():
    _verified_root = PROJECT_DIR

from pipeline.io import pick_verified_xlsx

_gene_names = []
if "GENE_OUTPUT_XLSX" in globals():
    _gene_names.append(Path(str(GENE_OUTPUT_XLSX)).name)
_gene_names += [
    "phenotype_gene_verified_combined_by_api_with_gwas_omim_with_3_rate.xlsx",
    "phenotype_gene_verified_combined_by_api_with_gwas_omim.xlsx",
]
_gene_order = []
_seen_g = set()
for _n in _gene_names:
    if _n not in _seen_g:
        _seen_g.add(_n)
        _gene_order.append(_n)

_snp_names = []
if "SNP_OUTPUT_XLSX" in globals():
    _snp_names.append(Path(str(SNP_OUTPUT_XLSX)).name)
_snp_names += [
    "phenotype_snp_verified_combined_by_api_with_gwas_omim_with_3_rate.xlsx",
    "phenotype_snp_verified_combined_by_api_with_gwas_omim.xlsx",
]
_snp_order = []
_seen_s = set()
for _n in _snp_names:
    if _n not in _seen_s:
        _seen_s.add(_n)
        _snp_order.append(_n)

_gene_min = len(globals()["gene_final"]) if "gene_final" in globals() else 1000
_snp_min = len(globals()["snp_final"]) if "snp_final" in globals() else 500

GENE_FILE = str(pick_verified_xlsx(_verified_root, _gene_order, "GENE", min_rows=_gene_min))
SNP_FILE = str(pick_verified_xlsx(_verified_root, _snp_order, "SNP", min_rows=_snp_min))

if globals().get("USE_JAMIA_SUPPLEMENTARY_FOR_FIGURES") and "JAMIA_GENE_XLSX" in globals() and "JAMIA_SNP_XLSX" in globals():
    if JAMIA_GENE_XLSX.is_file() and JAMIA_SNP_XLSX.is_file():
        GENE_FILE = JAMIA_GENE_XLSX
        SNP_FILE = JAMIA_SNP_XLSX
    elif JAMIA_GENE_XLSX.is_file() or JAMIA_SNP_XLSX.is_file():
        import warnings as _warnings
        _warnings.warn(
            "JAMIA supplementary: expected both Supplementary Data S1.xlsx and S2.xlsx under JAMIA_abstract_all_file; "
            f"gene_exists={JAMIA_GENE_XLSX.is_file()} snp_exists={JAMIA_SNP_XLSX.is_file()} — keeping paths from pick_verified_xlsx."
        )

SUMMARY_XLSX = "verification_structured_summary.xlsx"

def load_verified(path: str, task_type: str) -> pd.DataFrame:
    df = pd.read_excel(path)
    df = df.copy()
    df["TaskType"] = task_type

    for c in ["Evidence_Level", "GWAS_Support", "OMIM_Match", "Model"]:
        if c not in df.columns:
            df[c] = "NA"
    if "Triple_Category" not in df.columns:
        df["Triple_Category"] = ""
    if "Ensembl_Entity_Exists" not in df.columns:
        df["Ensembl_Entity_Exists"] = ""
    return df

gene_df = load_verified(GENE_FILE, "gene")
snp_df = load_verified(SNP_FILE, "snp")
all_df = pd.concat([gene_df, snp_df], ignore_index=True)

# Standard order for tables/plots (includes Not_Found from Ensembl entity gate)
level_order = ["Not_Found", "Strong", "Moderate", "Weak", "None"]
triple_order = ["Matched", "Not_Found", "Unmatched"]
source_order = ["Strong", "Weak", "None", "NA"]
omim_order = ["Strong", "Weak", "None", "NA"]

# Evidence_Level: missing/empty -> "None"; normalize legacy Chinese labels to English
all_df["Evidence_Level"] = all_df["Evidence_Level"].fillna("None").astype(str).str.strip()
all_df.loc[all_df["Evidence_Level"].isin(["", "NA", "nan", "NaN"]), "Evidence_Level"] = "None"
all_df["Evidence_Level"] = all_df["Evidence_Level"].replace({"Not_Found": "Not_Found"})

all_df["GWAS_Support"] = all_df["GWAS_Support"].fillna("NA").astype(str)
all_df["OMIM_Match"] = all_df["OMIM_Match"].fillna("NA").astype(str)
all_df["OMIM_Match"] = all_df["OMIM_Match"].replace({"High": "Strong", "Low": "Weak"})
all_df["Model"] = all_df["Model"].fillna("NA").astype(str)

# Triple summary
all_df["Triple_Category"] = all_df["Triple_Category"].fillna("").astype(str).str.strip()
all_df["Triple_Category"] = all_df["Triple_Category"].replace(
    {"Not_Found": "Not_Found", "Matched": "Matched", "Unmatched": "Unmatched"}
)
# Excel / export variants that are not exact triple_order tokens (else they become Unmatched)
_syn_triple = {
    "Not Found": "Not_Found",
    "NotFound": "Not_Found",
    "NOT_FOUND": "Not_Found",
    "not_found": "Not_Found",
}
all_df["Triple_Category"] = all_df["Triple_Category"].replace(_syn_triple)

# Triple_Category column only: normalize spelling/case from this field (no Evidence_Level inference).
_triple_key = (
    all_df["Triple_Category"]
    .astype(str)
    .str.strip()
    .str.lower()
    .str.replace(" ", "_", regex=False)
    .str.replace("-", "_", regex=False)
)
_triple_from_key = {
    "matched": "Matched",
    "not_found": "Not_Found",
    "unmatched": "Unmatched",
    "notfound": "Not_Found",
}
_bad_triple = ~all_df["Triple_Category"].isin(triple_order)
if bool(_bad_triple.any()):
    _mapped = _triple_key.loc[_bad_triple].map(_triple_from_key)
    all_df.loc[_bad_triple, "Triple_Category"] = _mapped.fillna(all_df.loc[_bad_triple, "Triple_Category"])

all_df.loc[all_df["Triple_Category"] == "", "Triple_Category"] = "Unmatched"
all_df.loc[~all_df["Triple_Category"].isin(triple_order), "Triple_Category"] = "Unmatched"

# Canonical TaskType labels so crosstab rows match reindex (avoids "SNP"/"snp " -> dropped counts).
all_df["TaskType"] = all_df["TaskType"].astype(str).str.strip().str.lower()
all_df["TaskType"] = all_df["TaskType"].replace({"snps": "snp"})

_task_type_order = ["gene", "snp"]
_trip_ct = pd.crosstab(all_df["TaskType"], all_df["Triple_Category"])
t_triple_counts = _trip_ct.reindex(index=_task_type_order, columns=triple_order, fill_value=0)
t_triple_pct = t_triple_counts.div(t_triple_counts.sum(axis=1).replace(0, np.nan), axis=0).fillna(0) * 100
print("\n=== Table 1A: Counts by TaskType x Triple_Category (Matched / Not_Found / Unmatched) ===")
print(t_triple_counts)
print("\n=== Table 1B: Row percentages by TaskType x Triple_Category (sum to 100%) ===")
print(t_triple_pct.round(2))
_row_sums = t_triple_pct.sum(axis=1).round(3)
print("\n=== Row-sum check (each TaskType row should be ~100) ===")
print(_row_sums)

# Overall Triple_Category (after normalization)
_tc_overall = all_df["Triple_Category"].value_counts().reindex(triple_order, fill_value=0)
_tc_overall_pct = (_tc_overall / _tc_overall.sum() * 100).round(2)
print("\n=== Table 1C: Triple_Category counts (overall, all rows) ===")
print(_tc_overall.to_string())
print("\n=== Table 1D: Triple_Category percentage (overall, sums to 100%) ===")
print(_tc_overall_pct.to_string())

# Matched-only: Strong / Moderate / Weak mix
_matched_level_order = ["Strong", "Moderate", "Weak"]
_matched_df = all_df.loc[all_df["Triple_Category"] == "Matched"].copy()
print(f"\n=== Matched subset size: {len(_matched_df)} / {len(all_df)} ===")
if len(_matched_df):
    evm = _matched_df["Evidence_Level"].fillna("None").astype(str).str.strip()
    evm = evm.replace({"": "None", "NA": "None", "nan": "None", "NaN": "None"})
    _core_ov = evm.value_counts().reindex(_matched_level_order, fill_value=0)
    _den_ov = int(_core_ov.sum())
    print("\n=== Table 1E: Among Matched — Evidence_Level counts (Strong / Moderate / Weak) ===")
    print(_core_ov.to_string())
    if _den_ov:
        print("\n=== Table 1F: Among Matched — % within Strong+Moderate+Weak (3 levels) ===")
        print((_core_ov / _den_ov * 100).round(2).to_string())

    t_matched_counts = pd.crosstab(_matched_df["TaskType"], evm).reindex(columns=_matched_level_order, fill_value=0)
    t_matched_pct = t_matched_counts.div(t_matched_counts.sum(axis=1).replace(0, np.nan), axis=0).fillna(0) * 100
    print("\n=== Table 1G: Among Matched — counts by TaskType x Evidence_Level (Strong/Moderate/Weak) ===")
    print(t_matched_counts)
    print("\n=== Table 1H: Among Matched — row % by TaskType x Evidence_Level (3 levels, each row sums to 100%) ===")
    print(t_matched_pct.round(2))
else:
    t_matched_counts = pd.DataFrame()
    t_matched_pct = pd.DataFrame()

# Structured table 2: Model x TaskType x Evidence_Level
t2_counts = (
    all_df.groupby(["Model", "TaskType", "Evidence_Level"], dropna=False)
    .size()
    .reset_index(name="Count")
)

t2_total = t2_counts.groupby(["Model", "TaskType"], dropna=False)["Count"].transform("sum")
t2_counts["Percentage"] = (t2_counts["Count"] / t2_total.replace(0, np.nan) * 100).fillna(0)

t2_counts = t2_counts.sort_values(["Model", "TaskType", "Evidence_Level"])
print("\n=== Table 2: Model x TaskType x Evidence_Level (count + percentage) ===")
print(t2_counts.to_string(index=False))

# Structured table 3: Source-side distributions
t3_gwas = (
    all_df.groupby(["TaskType", "GWAS_Support"], dropna=False)
    .size()
    .reset_index(name="Count")
)
t3_gwas["Percentage"] = (
    t3_gwas["Count"]
    / t3_gwas.groupby("TaskType")["Count"].transform("sum").replace(0, np.nan)
    * 100
).fillna(0)

t3_omim = (
    all_df.groupby(["TaskType", "OMIM_Match"], dropna=False)
    .size()
    .reset_index(name="Count")
)
t3_omim["Percentage"] = (
    t3_omim["Count"]
    / t3_omim.groupby("TaskType")["Count"].transform("sum").replace(0, np.nan)
    * 100
).fillna(0)

cross = (
    all_df.groupby(["TaskType", "GWAS_Support", "OMIM_Match"], dropna=False)
    .size()
    .reset_index(name="Count")
)

print("\n=== Table 3A: GWAS_Support by TaskType (count + percentage) ===")
print(t3_gwas.sort_values(["TaskType", "GWAS_Support"]).to_string(index=False))
print("\n=== Table 3B: OMIM_Match by TaskType (count + percentage) ===")
print(t3_omim.sort_values(["TaskType", "OMIM_Match"]).to_string(index=False))
print("\n=== Table 3C: TaskType x GWAS_Support x OMIM_Match (counts) ===")
print(cross.sort_values(["TaskType", "GWAS_Support", "OMIM_Match"]).to_string(index=False))

# Evidence score for boxplot
score_map = {"Not_Found": 0, "Strong": 3, "Moderate": 2, "Weak": 1, "None": 0}
all_df["Evidence_Score"] = all_df["Evidence_Level"].map(score_map).fillna(0)

# Multi-figure panel (Triple_Category + Matched evidence mix)
fig, axes = plt.subplots(1, 2, figsize=(18, 5.5))

# Plot Triple_Category by model (row percentage within each model).
_model_order = (
    all_df["Model"].astype(str).str.strip().value_counts().index.tolist()
)
_model_triple_counts = pd.crosstab(all_df["Model"], all_df["Triple_Category"]).reindex(
    index=_model_order,
    columns=triple_order,
    fill_value=0,
)
_x = np.arange(len(_model_triple_counts.index))
_bottom = np.zeros(len(_model_triple_counts.index), dtype=float)

# Put Not_Found on top so thin orange segment is easier to see.
_plot_order = ["Matched", "Unmatched", "Not_Found"]
_display_label = {"Matched": "Matched", "Unmatched": "Unmatched", "Not_Found": "Not Found"}
_color_map = dict(zip(triple_order, sns.color_palette("deep", n_colors=len(triple_order))))
_bar_handles = {}
for cat in _plot_order:
    _vals = _model_triple_counts[cat].astype(float).to_numpy()
    bars = axes[0].bar(
        _x,
        _vals,
        bottom=_bottom,
        label=_display_label[cat],
        color=_color_map[cat],
        width=0.55,
        linewidth=0.8,
        edgecolor="0.25",
    )
    _bar_handles[cat] = bars
    _bottom += _vals

axes[0].set_title("Triple Category Counts by Model (Stacked)")
axes[0].set_xlabel("Model")
axes[0].set_ylabel("Count")
axes[0].set_xticks(_x)
axes[0].set_xticklabels(list(_model_triple_counts.index), rotation=20, ha="right")
axes[0].legend(title="Triple Category", bbox_to_anchor=(1.02, 1), loc="upper left")

if len(_matched_df):
    evm_plot = _matched_df["Evidence_Level"].fillna("None").astype(str).str.strip()
    evm_plot = evm_plot.replace({"": "None", "NA": "None", "nan": "None", "NaN": "None"})

    _matched_model_order = (
        _matched_df["Model"].astype(str).str.strip().value_counts().index.tolist()
    )
    plot_m = pd.crosstab(_matched_df["Model"], evm_plot).reindex(
        index=_matched_model_order,
        columns=_matched_level_order,
        fill_value=0,
    )
    plot_m_pct = plot_m.div(plot_m.sum(axis=1).replace(0, np.nan), axis=0).fillna(0) * 100

    bottom = np.zeros(len(plot_m_pct))
    for lvl in _matched_level_order:
        vals = plot_m_pct[lvl].values if lvl in plot_m_pct.columns else np.zeros(len(plot_m_pct))
        axes[1].bar(range(len(plot_m_pct)), vals, bottom=bottom, label=lvl)
        bottom += vals

    axes[1].set_title("Among Matched: Strong / Moderate / Weak % Within Each Model")
    axes[1].set_ylabel("Percentage (%)")
    axes[1].set_xlabel("Model")
    axes[1].set_xticks(range(len(plot_m_pct)))
    axes[1].set_xticklabels(list(plot_m_pct.index), rotation=20, ha="right")
    axes[1].legend(title="Evidence Level", bbox_to_anchor=(1.02, 1), loc="upper left")
else:
    axes[1].text(0.5, 0.5, "No Matched rows", ha="center", va="center")

plt.tight_layout()
plt.show()

# Export structured tables to one workbook
with pd.ExcelWriter(SUMMARY_XLSX) as w:
    t_triple_counts.to_excel(w, sheet_name="t1a_triple_task_counts")
    t_triple_pct.round(2).to_excel(w, sheet_name="t1b_triple_task_pct")
    _tc_overall.to_frame(name="Count").to_excel(w, sheet_name="t1c_triple_overall_counts")
    _tc_overall_pct.to_frame(name="Percent").to_excel(w, sheet_name="t1d_triple_overall_pct")
    if len(_matched_df):
        t_matched_counts.to_excel(w, sheet_name="t1e_matched_evidence_counts")
        t_matched_pct.round(2).to_excel(w, sheet_name="t1f_matched_evidence_pct")
    t2_counts.to_excel(w, sheet_name="t2_model_task_level", index=False)
    t3_gwas.sort_values(["TaskType", "GWAS_Support"]).to_excel(w, sheet_name="t3_gwas", index=False)
    t3_omim.sort_values(["TaskType", "OMIM_Match"]).to_excel(w, sheet_name="t3_omim", index=False)
    cross.sort_values(["TaskType", "GWAS_Support", "OMIM_Match"]).to_excel(w, sheet_name="t3_cross", index=False)

print(f"\nSaved structured summary workbook: {SUMMARY_XLSX}")


# %% [code]
# Manuscript figure panel
import numpy as np
import pandas as pd
import matplotlib.pyplot as plt
import seaborn as sns

use_transparent_plots(context="talk")

# Reuse all_df from the previous analytics cell only if it already has a valid
# TaskType column with "gene"/"snp" values -- other cells (e.g. the Matched % summary
# cell) build their own all_df keyed by Source_Sheet instead of TaskType, and blindly
# reusing that would silently produce an all-NaN chart here.
_all_df_reusable = (
    "all_df" in globals()
    and "TaskType" in all_df.columns
    and set(all_df["TaskType"].astype(str).str.strip().str.lower().unique()) & {"gene", "snp"}
)
if not _all_df_reusable:
    from pathlib import Path as _Path_fb
    _proj_fb = _Path_fb(PROJECT_DIR) if "PROJECT_DIR" in globals() else _Path_fb(str(PROJECT_DIR))
    if "GENE_FILE" in globals() and "SNP_FILE" in globals() and _Path_fb(GENE_FILE).is_file() and _Path_fb(SNP_FILE).is_file():
        _gene_path = _Path_fb(GENE_FILE)
        _snp_path = _Path_fb(SNP_FILE)
    elif globals().get("USE_JAMIA_SUPPLEMENTARY_FOR_FIGURES") and "JAMIA_GENE_XLSX" in globals() and "JAMIA_SNP_XLSX" in globals() and JAMIA_GENE_XLSX.is_file() and JAMIA_SNP_XLSX.is_file():
        _gene_path = JAMIA_GENE_XLSX
        _snp_path = JAMIA_SNP_XLSX
    else:
        import warnings as _warnings

        def _pick_existing(root, names, label):
            for i, name in enumerate(names):
                p = root / name
                if p.is_file():
                    if i > 0:
                        _warnings.warn(f"{label}: using {name} (preferred output not found under {root}).")
                    return p
            raise FileNotFoundError(
                f"Missing {label} workbook under {root}. Tried: " + ", ".join(names)
            )

        _gene_candidates = [
            "phenotype_gene_verified_combined_by_api_with_gwas_omim_with_3_rate.xlsx",
            "phenotype_gene_verified_combined_by_api_with_gwas_omim.xlsx",
        ]
        _snp_candidates = [
            "phenotype_snp_verified_combined_by_api_with_gwas_omim_with_3_rate.xlsx",
            "phenotype_snp_verified_combined_by_api_with_gwas_omim.xlsx",
        ]
        if "GENE_OUTPUT_XLSX" in globals():
            _gene_candidates.insert(0, _Path_fb(str(GENE_OUTPUT_XLSX)).name)
        if "SNP_OUTPUT_XLSX" in globals():
            _snp_candidates.insert(0, _Path_fb(str(SNP_OUTPUT_XLSX)).name)
        _gene_path = _pick_existing(_proj_fb, _gene_candidates, "GENE")
        _snp_path = _pick_existing(_proj_fb, _snp_candidates, "SNP")

    gene_df = pd.read_excel(_gene_path)
    snp_df = pd.read_excel(_snp_path)
    gene_df = gene_df.copy(); gene_df["TaskType"] = "gene"
    snp_df = snp_df.copy(); snp_df["TaskType"] = "snp"
    all_df = pd.concat([gene_df, snp_df], ignore_index=True)

# Normalize key columns
for c in ["Model", "TaskType", "Evidence_Level", "GWAS_Support", "OMIM_Match"]:
    if c not in all_df.columns:
        all_df[c] = "NA"
all_df["Model"] = all_df["Model"].fillna("NA").astype(str)
all_df["TaskType"] = all_df["TaskType"].fillna("NA").astype(str)
all_df["Evidence_Level"] = all_df["Evidence_Level"].fillna("None").astype(str).str.strip()
all_df.loc[all_df["Evidence_Level"].isin(["", "NA", "nan", "NaN"]), "Evidence_Level"] = "None"
all_df["GWAS_Support"] = all_df["GWAS_Support"].fillna("None").astype(str).str.strip()
all_df.loc[all_df["GWAS_Support"].isin(["", "NA", "nan", "NaN"]), "GWAS_Support"] = "None"

all_df["OMIM_Match"] = all_df["OMIM_Match"].fillna("None").astype(str).str.strip()
all_df.loc[all_df["OMIM_Match"].isin(["", "NA", "nan", "NaN"]), "OMIM_Match"] = "None"
if "norm_omim_match_series" in globals():
    all_df["OMIM_Match"] = norm_omim_match_series(all_df["OMIM_Match"])
    all_df["GWAS_Support"] = norm_gwas_support_series(all_df["GWAS_Support"])
else:
    all_df["OMIM_Match"] = all_df["OMIM_Match"].replace({"High": "Strong", "Low": "Weak"})

score_map = {"Not_Found": 0, "Strong": 3, "Moderate": 2, "Weak": 1, "None": 0}
all_df["Evidence_Score"] = all_df["Evidence_Level"].map(score_map).fillna(0)

# Figure 4 (clinical lollipop) + GWAS/OMIM agreement heatmap. Lollipop
# numerator = Strong + Moderate; denominator = all rows (Not_Found
# included), matching the Table 3 / chi-square convention.
all_df["Model"] = all_df["Model"].astype(str).str.strip()
all_df["TaskType"] = all_df["TaskType"].astype(str).str.strip().str.lower()

_LOLLIPOP_MODELS = [
    m for m in (
        globals().get("ALLOWED_MODELS")
        or ["claude-sonnet-4-6", "deepseek-v4-flash", "gemini-3-flash-preview", "gpt-5.5"]
    )
    if m in set(all_df["Model"])
]
if not _LOLLIPOP_MODELS:
    _LOLLIPOP_MODELS = sorted(all_df["Model"].unique())

_SUPPORTED_LEVELS = {"Strong", "Moderate"}
_UNSUPPORTED_LEVELS = {"Weak", "None"}

def _clinical_support_pct(df: pd.DataFrame, model: str, task_type: str) -> float:
    """Strong+Moderate % of ALL rows for this (model, task_type), Not_Found included
    in the denominator -- matches the convention used throughout the manuscript."""
    sub = df[(df["Model"] == model) & (df["TaskType"] == task_type)]
    ev = sub["Evidence_Level"].astype(str).str.strip()
    doc = int(ev.isin(_SUPPORTED_LEVELS).sum())
    denom = len(sub)
    return (100.0 * doc / denom) if denom else np.nan

_lollipop_rows = []
for _m in _LOLLIPOP_MODELS:
    for _t in ("gene", "snp"):
        _lollipop_rows.append(
            {
                "Model": _m,
                "TaskType": _t,
                "Pct": _clinical_support_pct(all_df, _m, _t),
            }
        )
lollipop_df = pd.DataFrame(_lollipop_rows)

# Y-axis: one fixed order for both tasks (sort by gene rate, low -> high)
_gene_pct = lollipop_df[lollipop_df["TaskType"] == "gene"].set_index("Model")["Pct"]
model_order = sorted(_LOLLIPOP_MODELS, key=lambda m: _gene_pct.get(m, np.nan))
model_to_y = {m: i for i, m in enumerate(model_order)}

task_style = {
    "gene": {"color": "#2E6FBB", "offset": -0.18, "label": "gene"},
    "snp": {"color": "#9B59B6", "offset": 0.18, "label": "snp"},
}

fig_lo, ax_lo = plt.subplots(figsize=(8.5, 5.5))
for _t, _style in task_style.items():
    _xs, _ys, _lbls = [], [], []
    for _m in model_order:
        _pct = lollipop_df.loc[
            (lollipop_df["Model"] == _m) & (lollipop_df["TaskType"] == _t), "Pct"
        ]
        if _pct.empty or pd.isna(_pct.iloc[0]):
            continue
        _pct = float(_pct.iloc[0])
        _yi = model_to_y[_m] + _style["offset"]
        _xs.append(_pct)
        _ys.append(_yi)
        _lbls.append(f"{_pct:.1f}%")
    for _x, _y in zip(_xs, _ys):
        ax_lo.hlines(_y, 0, _x, colors=_style["color"], alpha=0.35, linewidth=2, zorder=1)
    ax_lo.scatter(
        _xs,
        _ys,
        s=120,
        color=_style["color"],
        edgecolors="white",
        linewidths=0.8,
        label=_style["label"],
        zorder=3,
    )
    for _x, _y, _lbl in zip(_xs, _ys, _lbls):
        ax_lo.text(
            _x,
            _y + 0.14,
            _lbl,
            ha="center",
            va="bottom",
            fontsize=9,
            color=_style["color"],
            zorder=4,
        )

ax_lo.set_yticks(np.arange(len(model_order)))
ax_lo.set_yticklabels(model_order)
ax_lo.set_xlim(0, 100)
ax_lo.set_ylim(-0.55, len(model_order) - 0.45)
ax_lo.set_xlabel("Strong + Moderate (% of all rows)")
ax_lo.set_title("Clinical Support Rate by Model (Lollipop)")
ax_lo.legend(title="TaskType", loc="lower right")
plt.tight_layout()
plt.show()

print("\n=== Lollipop rates (all rows, Not_Found included) ===")
print(
    lollipop_df.pivot(index="Model", columns="TaskType", values="Pct")
    .reindex(model_order)
    .round(1)
    .to_string()
)

task_order = ["gene", "snp"] if set(["gene", "snp"]).issubset(set(all_df["TaskType"])) else sorted(all_df["TaskType"].unique())
fig_gw, ax_gw = plt.subplots(figsize=(10, 5.5))
_panel_w, _panel_gap = 3, 0.8
for i, t in enumerate(task_order):
    sub = all_df[all_df["TaskType"] == t]
    h = pd.crosstab(sub["GWAS_Support"], sub["OMIM_Match"])
    _gwas_order = globals().get("GWAS_TIER_ORDER", ["Strong", "Weak", "None"])
    _omim_order = globals().get("OMIM_TIER_ORDER", ["Strong", "Weak", "None"])
    h = h.reindex(index=_gwas_order, columns=_omim_order, fill_value=0)
    x0 = i * (_panel_w + _panel_gap)
    mat = h.values
    ax_gw.imshow(mat, cmap="Blues", alpha=0.95, extent=(x0, x0 + h.shape[1], 0, h.shape[0]), aspect="auto", origin="lower")
    for r in range(h.shape[0]):
        for c in range(h.shape[1]):
            ax_gw.text(x0 + c + 0.5, r + 0.5, str(int(mat[r, c])), ha="center", va="center", fontsize=10)
    # Panel-type label sits inside the extended ylim, below set_title()'s pad --
    # placing it at h.shape[0] + 0.18 (above the old ylim=3 ceiling) collided
    # with the title because tight_layout() doesn't reserve space for text
    # drawn outside the axes' data limits.
    ax_gw.text(x0 + h.shape[1] / 2, h.shape[0] + 0.12, t, ha="center", va="bottom", fontsize=12, fontweight="bold")

_gwas_lbls = globals().get("GWAS_TIER_AXIS_LABELS", ["GWAS Strong", "GWAS Weak", "GWAS None"])
_omim_lbls = globals().get("OMIM_TIER_AXIS_LABELS", ["OMIM Strong", "OMIM Weak", "OMIM None"])
_last_x0 = (len(task_order) - 1) * (_panel_w + _panel_gap)
ax_gw.set_xlim(0, _last_x0 + _panel_w)
ax_gw.set_ylim(0, 3.45)
ax_gw.set_yticks(np.arange(0.5, 3.5, 1))
ax_gw.set_yticklabels(_gwas_lbls)
xticks = []
xticklabels = []
for i in range(len(task_order)):
    x0 = i * (_panel_w + _panel_gap)
    for j, lbl in enumerate(_omim_lbls):
        xticks.append(x0 + j + 0.5)
        xticklabels.append(lbl)
ax_gw.set_xticks(xticks)
ax_gw.set_xticklabels(xticklabels, rotation=45, ha="right")
ax_gw.set_title("Supplementary Figure S2. GWAS vs OMIM agreement (split by task type)", pad=16)
ax_gw.set_xlabel("OMIM match tier")
ax_gw.set_ylabel("GWAS support tier")
plt.tight_layout()
plt.show()


# %% [code]
# Table 3: chi-square documentation-rate comparison across models
# (Overall / Gene / SNP). All rows included -- Not_Found counts as
# undocumented, matching the convention used throughout the manuscript.
import pandas as pd
from scipy.stats import chi2_contingency
from itertools import combinations

_all_df_reusable_t3 = (
    "all_df" in globals()
    and "TaskType" in all_df.columns
    and set(all_df["TaskType"].astype(str).str.strip().str.lower().unique()) & {"gene", "snp"}
)
if not _all_df_reusable_t3:
    gene_df = pd.read_excel(GENE_FILE); gene_df = gene_df.copy(); gene_df["TaskType"] = "gene"
    snp_df = pd.read_excel(SNP_FILE); snp_df = snp_df.copy(); snp_df["TaskType"] = "snp"
    all_df = pd.concat([gene_df, snp_df], ignore_index=True)

all_df["Model"] = all_df["Model"].astype(str).str.strip()
all_df["TaskType"] = all_df["TaskType"].astype(str).str.strip().str.lower()

_T3_MODEL_ORDER = ["claude-sonnet-4-6", "deepseek-v4-flash", "gemini-3-flash-preview", "gpt-5.5"]
_SUPPORTED = {"Strong", "Moderate"}

def _doc_counts(df):
    out = {}
    for m in _T3_MODEL_ORDER:
        sub = df[df["Model"] == m]
        doc = int(sub["Evidence_Level"].astype(str).str.strip().isin(_SUPPORTED).sum())
        out[m] = (doc, len(sub) - doc, len(sub))
    return out

def _omnibus(counts):
    table = [[counts[m][0], counts[m][1]] for m in _T3_MODEL_ORDER]
    chi2, p, dof, _ = chi2_contingency(table, correction=False)
    return chi2, dof, p

rows = []
counts_overall = _doc_counts(all_df)
counts_gene = _doc_counts(all_df[all_df["TaskType"] == "gene"])
counts_snp = _doc_counts(all_df[all_df["TaskType"] == "snp"])
for m in _T3_MODEL_ORDER:
    o_doc, _, o_tot = counts_overall[m]
    g_doc, _, g_tot = counts_gene[m]
    s_doc, _, s_tot = counts_snp[m]
    rows.append({
        "Model": m,
        "Overall doc/total": f"{o_doc}/{o_tot}", "Overall Rate (%)": round(o_doc / o_tot * 100, 1),
        "Gene doc/total": f"{g_doc}/{g_tot}", "Gene Rate (%)": round(g_doc / g_tot * 100, 1),
        "SNP doc/total": f"{s_doc}/{s_tot}", "SNP Rate (%)": round(s_doc / s_tot * 100, 1),
    })
table3_summary = pd.DataFrame(rows)
print("=== Table 3 (all rows, Not_Found included) ===")
print(table3_summary.to_string(index=False))

for label, counts in [("Overall", counts_overall), ("Gene subgroup", counts_gene), ("SNP subgroup", counts_snp)]:
    chi2, dof, p = _omnibus(counts)
    print(f"{label}: chi2({dof}) = {chi2:.2f}, p = {p:.3g}")

print()
pairwise_rows = []
n_pairs = 6
for a, b in combinations(_T3_MODEL_ORDER, 2):
    table = [[counts_overall[a][0], counts_overall[a][1]], [counts_overall[b][0], counts_overall[b][1]]]
    chi2, p, dof, _ = chi2_contingency(table, correction=False)
    p_bonf = min(p * n_pairs, 1.0)
    pairwise_rows.append({
        "Pairwise comparison": f"{a} vs {b}",
        "chi2": round(chi2, 2), "df": dof,
        "Raw p": p, "Bonferroni p": p_bonf,
        "Significant": "*" if p_bonf < 0.05 else "—",
    })
table3_pairwise = pd.DataFrame(pairwise_rows)
print("=== Table 3 pairwise (Overall, Bonferroni x6) ===")
print(table3_pairwise.to_string(index=False))


# %% [code]
# Contribution and consistency figures:
# 1) GWAS vs OMIM contribution rate within each Evidence_Level
# 2) Per-model consistency (GWAS vs OMIM agreement)
import numpy as np
import pandas as pd
import matplotlib.pyplot as plt
import seaborn as sns

use_transparent_plots(context="talk")

# Reuse all_df if available; otherwise load files.
if "all_df" not in globals():
    from pathlib import Path as _Path_fb
    _proj_fb = _Path_fb(PROJECT_DIR) if "PROJECT_DIR" in globals() else _Path_fb(str(PROJECT_DIR))
    if "GENE_FILE" in globals() and "SNP_FILE" in globals() and _Path_fb(GENE_FILE).is_file() and _Path_fb(SNP_FILE).is_file():
        _gene_path = _Path_fb(GENE_FILE)
        _snp_path = _Path_fb(SNP_FILE)
    elif globals().get("USE_JAMIA_SUPPLEMENTARY_FOR_FIGURES") and "JAMIA_GENE_XLSX" in globals() and "JAMIA_SNP_XLSX" in globals() and JAMIA_GENE_XLSX.is_file() and JAMIA_SNP_XLSX.is_file():
        _gene_path = JAMIA_GENE_XLSX
        _snp_path = JAMIA_SNP_XLSX
    else:
        import warnings as _warnings

        def _pick_existing(root, names, label):
            for i, name in enumerate(names):
                p = root / name
                if p.is_file():
                    if i > 0:
                        _warnings.warn(f"{label}: using {name} (preferred output not found under {root}).")
                    return p
            raise FileNotFoundError(
                f"Missing {label} workbook under {root}. Tried: " + ", ".join(names)
            )

        _gene_candidates = [
            "phenotype_gene_verified_combined_by_api_with_gwas_omim_with_3_rate.xlsx",
            "phenotype_gene_verified_combined_by_api_with_gwas_omim.xlsx",
        ]
        _snp_candidates = [
            "phenotype_snp_verified_combined_by_api_with_gwas_omim_with_3_rate.xlsx",
            "phenotype_snp_verified_combined_by_api_with_gwas_omim.xlsx",
        ]
        if "GENE_OUTPUT_XLSX" in globals():
            _gene_candidates.insert(0, _Path_fb(str(GENE_OUTPUT_XLSX)).name)
        if "SNP_OUTPUT_XLSX" in globals():
            _snp_candidates.insert(0, _Path_fb(str(SNP_OUTPUT_XLSX)).name)
        _gene_path = _pick_existing(_proj_fb, _gene_candidates, "GENE")
        _snp_path = _pick_existing(_proj_fb, _snp_candidates, "SNP")

    gene_df = pd.read_excel(_gene_path)
    snp_df = pd.read_excel(_snp_path)
    gene_df = gene_df.copy(); gene_df["TaskType"] = "gene"
    snp_df = snp_df.copy(); snp_df["TaskType"] = "snp"
    all_df = pd.concat([gene_df, snp_df], ignore_index=True)

# Normalize fields
for c in ["Evidence_Level", "GWAS_Support", "OMIM_Match", "Model", "TaskType"]:
    if c not in all_df.columns:
        all_df[c] = "None"

all_df = all_df.copy()
all_df["Evidence_Level"] = all_df["Evidence_Level"].fillna("None").astype(str).str.strip()
all_df["GWAS_Support"] = all_df["GWAS_Support"].fillna("None").astype(str).str.strip()
all_df["OMIM_Match"] = all_df["OMIM_Match"].fillna("None").astype(str).str.strip()
all_df["Model"] = all_df["Model"].fillna("NA").astype(str).str.strip()
all_df["TaskType"] = all_df["TaskType"].fillna("NA").astype(str).str.strip()

for col in ["Evidence_Level", "GWAS_Support", "OMIM_Match"]:
    all_df.loc[all_df[col].isin(["", "NA", "nan", "NaN"]), col] = "None"

all_df["OMIM_Match"] = all_df["OMIM_Match"].replace({"High": "Strong", "Low": "Weak"})

# Map levels to numeric strength for contribution/consistency logic
gwas_score_map = {"Strong": 2, "Weak": 1, "None": 0}
omim_score_map = {"Strong": 2, "Weak": 1, "None": 0}

all_df["gwas_score"] = all_df["GWAS_Support"].map(gwas_score_map).fillna(0)
all_df["omim_score"] = all_df["OMIM_Match"].map(omim_score_map).fillna(0)

# Source contribution logic per row:
# - both_zero: both no support
# - both: both provide support
# - gwas_only / omim_only
conditions = [
    (all_df["gwas_score"] == 0) & (all_df["omim_score"] == 0),
    (all_df["gwas_score"] > 0) & (all_df["omim_score"] > 0),
    (all_df["gwas_score"] > 0) & (all_df["omim_score"] == 0),
    (all_df["gwas_score"] == 0) & (all_df["omim_score"] > 0),
]
choices = ["both_zero", "both", "gwas_only", "omim_only"]
all_df["contrib_class"] = np.select(conditions, choices, default="both_zero")

# Figure A: contribution rate inside each Evidence_Level
level_order = ["Not_Found", "Strong", "Moderate", "Weak", "None"]
contrib_order = ["gwas_only", "omim_only", "both", "both_zero"]

contrib = (
    all_df.groupby(["Evidence_Level", "contrib_class"], dropna=False)
    .size()
    .reset_index(name="Count")
)
contrib = contrib[contrib["Evidence_Level"].isin(level_order)].copy()

# ensure full grid
full_grid = pd.MultiIndex.from_product([level_order, contrib_order], names=["Evidence_Level", "contrib_class"]).to_frame(index=False)
contrib = full_grid.merge(contrib, on=["Evidence_Level", "contrib_class"], how="left")
contrib["Count"] = contrib["Count"].fillna(0)

contrib["Total"] = contrib.groupby("Evidence_Level")["Count"].transform("sum")
contrib["Pct"] = (contrib["Count"] / contrib["Total"].replace(0, np.nan) * 100).fillna(0)

pivot_contrib = contrib.pivot(index="Evidence_Level", columns="contrib_class", values="Pct").reindex(index=level_order, columns=contrib_order, fill_value=0)

# Figure B: model consistency = normalized closeness of GWAS and OMIM
# strength: 1 - |gwas_score - omim_score| / 2, in [0, 1]
all_df["consistency_score"] = 1.0 - (all_df["gwas_score"] - all_df["omim_score"]).abs() / 2.0

consistency = (
    all_df.groupby(["Model", "TaskType"], dropna=False)
    .agg(
        MeanConsistency=("consistency_score", "mean"),
        N=("consistency_score", "size"),
        FullAgreeRate=("consistency_score", lambda s: (s == 1.0).mean() * 100),
    )
    .reset_index()
)

print("\n=== Contribution table (Evidence_Level x contribution class, %) ===")
print(pivot_contrib.round(2).to_string())
print("\n=== Model consistency table ===")
print(consistency.sort_values(["TaskType", "MeanConsistency"], ascending=[True, False]).to_string(index=False))

fig, axes = plt.subplots(1, 2, figsize=(20, 7))

# A1: stacked percentage bars
_contrib_display = {"gwas_only": "GWAS only", "omim_only": "OMIM only", "both": "Both", "both_zero": "Both zero"}
_level_display = {"Not_Found": "Not Found", "Strong": "Strong", "Moderate": "Moderate", "Weak": "Weak", "None": "None"}
bottom = np.zeros(len(pivot_contrib))
_x_labels = [_level_display[lvl] for lvl in pivot_contrib.index]
for c in contrib_order:
    vals = pivot_contrib[c].values
    axes[0].bar(_x_labels, vals, bottom=bottom, label=_contrib_display[c])
    bottom += vals
axes[0].set_ylim(0, 100)
axes[0].set_title("GWAS vs OMIM contribution rate within each Evidence Level")
axes[0].set_xlabel("Evidence Level")
axes[0].set_ylabel("Percentage (%)")
axes[0].legend(title="Contribution class", bbox_to_anchor=(1.02, 1), loc="upper left")

# A2: model consistency bars (grouped by task)
sns.barplot(
    data=consistency,
    x="Model",
    y="MeanConsistency",
    hue="TaskType",
    ax=axes[1],
)
axes[1].set_title("Per-model consistency (GWAS vs OMIM)")
axes[1].set_xlabel("Model")
axes[1].set_ylabel("Mean consistency score (0-1)")
axes[1].set_ylim(0, 1.05)
axes[1].tick_params(axis="x", rotation=35)

# annotate N
for p in axes[1].patches:
    h = p.get_height()
    if np.isfinite(h):
        axes[1].annotate(f"{h:.2f}", (p.get_x() + p.get_width() / 2, h), ha="center", va="bottom", fontsize=9, xytext=(0, 3), textcoords="offset points")

plt.tight_layout()
plt.show()


# %% [code]
# Summary tables built from the current *_with_3_rate.xlsx outputs:
# - Gene / SNP: Ensembl Found (N, Found, Not_Found)
# - Gene / SNP: Matched % + Strong/Moderate/Weak evidence mix (S/M/W)
# - Key Takeaways (best cells from the Ensembl Found tables)
import os
import warnings
import numpy as np
import pandas as pd
from pathlib import Path
from IPython.display import display, HTML

_verified_root = Path(PROJECT_DIR) if "PROJECT_DIR" in globals() else Path.cwd()
if not _verified_root.is_dir():
    _verified_root = PROJECT_DIR

def _pick_existing(root, names, label):
    for i, name in enumerate(names):
        p = root / name
        if p.is_file():
            if i > 0:
                warnings.warn(f"{label}: using {name} (preferred output not found under {root}).")
            return str(p)
    raise FileNotFoundError(
        f"Missing {label} workbook under {root}. Tried: " + ", ".join(names)
    )

_gene_names = []
if "GENE_OUTPUT_XLSX" in globals():
    _gene_names.append(Path(str(GENE_OUTPUT_XLSX)).name)
_gene_names += [
    "phenotype_gene_verified_combined_by_api_with_gwas_omim_with_3_rate.xlsx",
    "phenotype_gene_verified_combined_by_api_with_gwas_omim.xlsx",
]
_gene_order = []
_seen_g = set()
for _n in _gene_names:
    if _n not in _seen_g:
        _seen_g.add(_n)
        _gene_order.append(_n)

_snp_names = []
if "SNP_OUTPUT_XLSX" in globals():
    _snp_names.append(Path(str(SNP_OUTPUT_XLSX)).name)
_snp_names += [
    "phenotype_snp_verified_combined_by_api_with_gwas_omim_with_3_rate.xlsx",
    "phenotype_snp_verified_combined_by_api_with_gwas_omim.xlsx",
]
_snp_order = []
_seen_s = set()
for _n in _snp_names:
    if _n not in _seen_s:
        _seen_s.add(_n)
        _snp_order.append(_n)

GENE_FILE = _pick_existing(_verified_root, _gene_order, "GENE")
SNP_FILE = _pick_existing(_verified_root, _snp_order, "SNP")

if globals().get("USE_JAMIA_SUPPLEMENTARY_FOR_FIGURES") and "JAMIA_GENE_XLSX" in globals() and "JAMIA_SNP_XLSX" in globals():
    if JAMIA_GENE_XLSX.is_file() and JAMIA_SNP_XLSX.is_file():
        GENE_FILE = JAMIA_GENE_XLSX
        SNP_FILE = JAMIA_SNP_XLSX
    elif JAMIA_GENE_XLSX.is_file() or JAMIA_SNP_XLSX.is_file():
        import warnings as _warnings
        _warnings.warn(
            "JAMIA supplementary: expected both Supplementary Data S1.xlsx and S2.xlsx under JAMIA_abstract_all_file; "
            f"gene_exists={JAMIA_GENE_XLSX.is_file()} snp_exists={JAMIA_SNP_XLSX.is_file()} — keeping paths from _pick_existing."
        )

gene_df = pd.read_excel(GENE_FILE).copy()
snp_df = pd.read_excel(SNP_FILE).copy()
all_df = pd.concat([gene_df, snp_df], ignore_index=True)

# -------- normalize columns --------
if "Model" not in all_df.columns:
    raise RuntimeError("Column 'Model' not found.")
if "Source_Sheet" not in all_df.columns:
    raise RuntimeError("Column 'Source_Sheet' not found.")

all_df["Model"] = all_df["Model"].fillna("NA").astype(str).str.strip()
all_df["Source_Sheet"] = all_df["Source_Sheet"].fillna("NA").astype(str).str.strip()

# Triple_Category fallback (if available)
if "Triple_Category" in all_df.columns:
    tc = all_df["Triple_Category"].fillna("").astype(str).str.strip()
    tc = tc.replace(
        {
            "Not_Foun": "Not_Found",
            "Matched": "Matched",
            "Unmatched": "Unmatched",
            "Not Found": "Not_Found",
            "NotFound": "Not_Found",
            "NOT_FOUND": "Not_Found",
            "not_found": "Not_Found",
        }
    )
    key = tc.str.lower().str.replace(" ", "_", regex=False).str.replace("-", "_", regex=False)
    tc = key.map({"not_found": "Not_Found", "matched": "Matched", "unmatched": "Unmatched"}).fillna(tc)
else:
    tc = pd.Series([""] * len(all_df), index=all_df.index)

# Ensembl_Entity_Exists preferred when available; otherwise fallback to Triple_Category
if "Ensembl_Entity_Exists" in all_df.columns:
    exists_raw = all_df["Ensembl_Entity_Exists"].fillna("").astype(str).str.strip().str.lower()
    true_set = {"true", "1", "yes", "y", "t"}
    false_set = {"false", "0", "no", "n", "f"}

    found_status = np.where(
        exists_raw.isin(true_set),
        "Found",
        np.where(
            exists_raw.isin(false_set),
            "Not_Found",
            np.where(tc.eq("Not_Found"), "Not_Found", "Found"),
        ),
    )
else:
    found_status = np.where(tc.eq("Not_Found"), "Not_Found", "Found")

all_df["Found_Status"] = found_status

TRIPLE_LABELS = frozenset({"Matched", "Not_Found", "Unmatched"})

def _norm_triple_labels(s: pd.Series) -> pd.Series:
    x = s.fillna("").astype(str).str.strip()
    x = x.replace({"": "Unmatched"})
    x = x.replace({"Not_Found": "Not_Found", "Matched": "Matched", "Unmatched": "Unmatched"})
    x = x.replace(
        {"Not Found": "Not_Found", "NotFound": "Not_Found", "NOT_FOUND": "Not_Found", "not_found": "Not_Found"}
    )
    x = x.where(x.isin(TRIPLE_LABELS), other="Unmatched")
    return x

def _norm_evidence_levels(s: pd.Series) -> pd.Series:
    x = s.fillna("None").astype(str).str.strip()
    x = x.replace({"": "None", "NA": "None", "nan": "None", "NaN": "None", "Not_Found": "Not_Found"})
    return x

if "Triple_Category" in all_df.columns:
    all_df["Triple_Norm"] = _norm_triple_labels(all_df["Triple_Category"])
else:
    all_df["Triple_Norm"] = "Unmatched"

_PREFERRED_MODELS = ['claude-sonnet-4-6', 'deepseek-v4-flash', 'gemini-3-flash-preview', 'gpt-5.5']

def _resolve_model_order(df: pd.DataFrame) -> list[str]:
    present = set(df["Model"].astype(str).str.strip())
    ordered = [m for m in _PREFERRED_MODELS if m in present]
    ordered.extend(sorted(present.difference(ordered)))
    return ordered

MODEL_ORDER = _resolve_model_order(all_df)
GENE_TASKS = TASK_ORDER_SIX[:3]
SNP_TASKS = TASK_ORDER_SIX[3:]

def build_ensembl_found_table(df: pd.DataFrame, task_cols: list[str]) -> pd.DataFrame:
    rows = []
    for model in MODEL_ORDER:
        sub_m = df[df["Model"] == model]
        row = {"Model": model}
        for task in task_cols:
            sub = sub_m[sub_m["Source_Sheet"] == task]
            total = len(sub)
            found_cnt = int((sub["Found_Status"] == "Found").sum())
            nf_cnt = int((sub["Found_Status"] == "Not_Found").sum())

            if total == 0:
                row[task] = "NA"
            else:
                found_pct = found_cnt / total * 100
                row[task] = (
                    f"{found_pct:.2f}% "
                    f"(N={total}, Found={found_cnt}, Not Found={nf_cnt})"
                )
        rows.append(row)
    return pd.DataFrame(rows)

def build_matched_evidence_table(df: pd.DataFrame, task_cols: list[str]) -> pd.DataFrame:
    """Matched % headline; (S/M/W) = Strong / Moderate / Weak as % of all rows (within Matched only)."""
    rows = []
    for model in MODEL_ORDER:
        sub_m = df[df["Model"] == model]
        row: dict = {"Model": model}
        for task in task_cols:
            sub = sub_m[sub_m["Source_Sheet"] == task]
            total = len(sub)
            if total == 0:
                row[task] = "NA"
                continue
            matched = sub["Triple_Norm"] == "Matched"
            matched_pct = float(matched.mean() * 100.0)
            if "Evidence_Level" in sub.columns:
                ev = _norm_evidence_levels(sub.loc[matched, "Evidence_Level"])
                s_cnt = int(ev.eq("Strong").sum())
                m_cnt = int(ev.eq("Moderate").sum())
                w_cnt = int(ev.eq("Weak").sum())
            else:
                s_cnt = m_cnt = w_cnt = 0
            s_pct = s_cnt / total * 100.0
            m_pct = m_cnt / total * 100.0
            w_pct = w_cnt / total * 100.0
            row[task] = f"{matched_pct:.2f}% (S{s_pct:.2f}/M{m_pct:.2f}/W{w_pct:.2f})"
        rows.append(row)
    return pd.DataFrame(rows)

def max_found_task(found_tbl: pd.DataFrame, task_cols: list[str]) -> tuple[str, str, float]:
    best_model = None
    best_task = None
    best_pct = -1.0
    for _, r in found_tbl.iterrows():
        model = r["Model"]
        for task in task_cols:
            v = r[task]
            if isinstance(v, str) and "%" in v:
                pct = float(v.split("%", 1)[0])
                if pct > best_pct:
                    best_pct = pct
                    best_model = model
                    best_task = task
    return best_model, best_task, best_pct

def task_all_zero_not_found(df: pd.DataFrame, task_cols: list[str]) -> list[str]:
    all_zero_tasks = []
    for task in task_cols:
        sub = df[df["Source_Sheet"] == task]
        total = len(sub)
        if total == 0:
            continue
        nf_pct = (sub["Found_Status"] == "Not_Found").mean() * 100
        if np.isclose(nf_pct, 100.0):
            all_zero_tasks.append(task)
    return all_zero_tasks

def style_dark(df: pd.DataFrame, title: str):
    import html as _html

    display(HTML(f"<h2 style='margin:8px 0 10px 0; color:#E5E7EB;'>{_html.escape(title)}</h2>"))
    _table_css = (
        "<style>"
        ".style-dark-wrap table { border-collapse: collapse; width: 100%; }"
        ".style-dark-wrap th { background-color: #111827; color: #F9FAFB; "
        "border: 1px solid #374151; padding: 8px; text-align: left; }"
        ".style-dark-wrap td { background-color: #0B0F16; color: #E5E7EB; "
        "border: 1px solid #374151; padding: 8px; }"
        "</style>"
    )
    try:
        import jinja2  # noqa: F401 — pandas Styler requires jinja2

        sty = df.style.set_table_styles(
            [
                {"selector": "table", "props": "border-collapse: collapse; width: 100%;"},
                {"selector": "th", "props": "background-color: #111827; color: #F9FAFB; border: 1px solid #374151; padding: 8px; text-align: left;"},
                {"selector": "td", "props": "background-color: #0B0F16; color: #E5E7EB; border: 1px solid #374151; padding: 8px;"},
                {"selector": "tr:hover td", "props": "background-color: #111827;"},
            ]
        )
        display(sty)
        return
    except (ImportError, AttributeError):
        pass
    tbl = df.to_html(index=True, border=0, escape=True)
    display(HTML(_table_css + f'<div class="style-dark-wrap">{tbl}</div>'))

gene_tbl_found = build_ensembl_found_table(all_df, GENE_TASKS)
snp_tbl_found = build_ensembl_found_table(all_df, SNP_TASKS)
gene_tbl_smw = build_matched_evidence_table(all_df, GENE_TASKS)
snp_tbl_smw = build_matched_evidence_table(all_df, SNP_TASKS)

style_dark(gene_tbl_found, "Gene Table (Ensembl Found — N / Found / Not Found)")
style_dark(snp_tbl_found, "SNP Table (Ensembl Found — N / Found / Not Found)")
style_dark(gene_tbl_smw, "Gene Table (Matched %; S/M/W = Strong / Moderate / Weak as % of all rows)")
style_dark(snp_tbl_smw, "SNP Table (Matched %; S/M/W = Strong / Moderate / Weak as % of all rows)")

# -------- key takeaways (from Ensembl Found tables) --------
best_gene_model, best_gene_task, best_gene_pct = max_found_task(gene_tbl_found, GENE_TASKS)
best_snp_model, best_snp_task, best_snp_pct = max_found_task(snp_tbl_found, SNP_TASKS)
full_not_found_tasks = task_all_zero_not_found(all_df, GENE_TASKS + SNP_TASKS)

if full_not_found_tasks:
    bottleneck_text = ", ".join(full_not_found_tasks)
    bottleneck_conclusion = f"{bottleneck_text} has 100% Not Found across all rows."
else:
    bottleneck_conclusion = "No task is at 100% Not Found globally."

key_takeaways = pd.DataFrame(
    {
        "Topic": [
            "Gene best",
            "SNP best",
            "Not Found bottleneck",
        ],
        "Conclusion": [
            f"{best_gene_model} leads on {best_gene_task} ({best_gene_pct:.2f}% Found).",
            f"{best_snp_model} leads on {best_snp_task} ({best_snp_pct:.2f}% Found).",
            bottleneck_conclusion,
        ],
    }
)

style_dark(key_takeaways, "Key Takeaways")


# %% [markdown]
## 3. Statistical tests


# %% [code]
import numpy as np
import pandas as pd
from scipy.stats import chi2_contingency
from itertools import combinations
from IPython.display import display, HTML

# Documentation counts derived from Table 2 (all_df / t2_counts).
# doc = Strong + Moderate; undoc = Weak + None; Not_Found rows are
# excluded from all denominators here.

MODEL_LABEL = {
    "claude-sonnet-4-6": "Claude Sonnet 4.6",
    "gpt-5.5": "GPT-5.5",
    "gemini-3-flash-preview": "Gemini 3 Flash Preview",
    "deepseek-v4-flash": "DeepSeek V4 Flash",
}
_DOC_LEVELS = frozenset({"Strong", "Moderate"})
_UNDOC_LEVELS = frozenset({"Weak", "None"})


def _ensure_t2_counts() -> pd.DataFrame:
    if "t2_counts" in globals() and isinstance(t2_counts, pd.DataFrame) and len(t2_counts):
        return t2_counts.copy()
    if "all_df" not in globals():
        raise RuntimeError("Run the analytics cell above first (need all_df or t2_counts).")
    return (
        all_df.groupby(["Model", "TaskType", "Evidence_Level"], dropna=False)
        .size()
        .reset_index(name="Count")
    )


def build_documentation_data(t2: pd.DataFrame) -> dict:
    pref = list(
        globals().get(
            "ALLOWED_MODELS",
            ["claude-sonnet-4-6", "deepseek-v4-flash", "gemini-3-flash-preview", "gpt-5.5"],
        )
    )
    present = set(t2["Model"].astype(str).str.strip())
    model_slugs = [m for m in pref if m in present] + sorted(present.difference(pref))

    out = {}
    for slug in model_slugs:
        label = MODEL_LABEL.get(slug, slug)
        counts = {"gene_doc": 0, "gene_undoc": 0, "snp_doc": 0, "snp_undoc": 0}
        sub = t2[t2["Model"].astype(str).str.strip() == slug]
        for _, row in sub.iterrows():
            tt = str(row["TaskType"]).strip().lower()
            ev = str(row["Evidence_Level"]).strip()
            n = int(row["Count"])
            if ev in _DOC_LEVELS:
                key = f"{tt}_doc"
            elif ev in _UNDOC_LEVELS:
                key = f"{tt}_undoc"
            else:
                continue
            if key in counts:
                counts[key] += n
        out[label] = counts
    return out


_t2_src = _ensure_t2_counts()
data = build_documentation_data(_t2_src)
if not data:
    raise RuntimeError("No models found in Table 2 counts.")

SHORT = {label: label for label in data}

print("Documentation counts derived from Table 2 (Model × TaskType × Evidence_Level):")
for _m, _v in data.items():
    print(
        f"  {_m}: gene {_v['gene_doc']}/{_v['gene_doc'] + _v['gene_undoc']}, "
        f"snp {_v['snp_doc']}/{_v['snp_doc'] + _v['snp_undoc']}"
    )

for m, v in data.items():
    v["all_doc"]   = v["gene_doc"]   + v["snp_doc"]
    v["all_undoc"] = v["gene_undoc"] + v["snp_undoc"]
    v["all_n"]     = v["all_doc"]    + v["all_undoc"]
    v["gene_n"]    = v["gene_doc"]   + v["gene_undoc"]
    v["snp_n"]     = v["snp_doc"]    + v["snp_undoc"]
    v["all_pct"]   = v["all_doc"]    / v["all_n"]  * 100
    v["gene_pct"]  = v["gene_doc"]   / v["gene_n"] * 100
    v["snp_pct"]   = v["snp_doc"]    / v["snp_n"]  * 100

def chisq(counts):
    """Run a Pearson chi-square test on a 2D contingency table."""
    chi2, p, dof, _ = chi2_contingency(np.array(counts))
    return chi2, p, dof

# Overall 4x2 test across all tasks
ov = chisq([[v["all_doc"],  v["all_undoc"]]  for v in data.values()])
# Gene-only subgroup test
gn = chisq([[v["gene_doc"], v["gene_undoc"]] for v in data.values()])
# SNP-only subgroup test
sn = chisq([[v["snp_doc"],  v["snp_undoc"]]  for v in data.values()])

# Pairwise comparisons with Bonferroni correction (n=6 pairs, alpha=0.05/6)
model_names = list(data.keys())
pairs = []
for (i, n1), (j, n2) in combinations(enumerate(model_names), 2):
    v1, v2 = data[n1], data[n2]
    c2, p, _ = chisq([[v1["all_doc"], v1["all_undoc"]], [v2["all_doc"], v2["all_undoc"]]])
    p_bonf = min(p * 6, 1.0)
    pairs.append((SHORT[n1], SHORT[n2], c2, p, p_bonf, p_bonf < 0.05))

# Formatting helpers: HTML / LaTeX / plain-text p-value strings for the paper.
def fmt_p(p):
    """HTML p-value."""
    if p < 0.001:
        exp = int(np.floor(np.log10(p)))
        coef = p / 10**exp
        return f"{coef:.2f} &times; 10<sup>{exp}</sup>"
    return f"{p:.4f}"

def fmt_p_latex(p):
    """LaTeX p-value."""
    if p < 0.001:
        exp = int(np.floor(np.log10(p)))
        coef = p / 10**exp
        return f"${coef:.2f} \\times 10^{{{exp}}}$"
    return f"{p:.4f}"

def fmt_p_plain(p):
    """Plain text / Word p-value."""
    if p < 0.001:
        exp = int(np.floor(np.log10(p)))
        coef = p / 10**exp
        return f"{coef:.2e} (={coef:.2f}×10^{exp})"
    return f"{p:.4f}"

LATEX_MODEL = {label: label for label in data}

# Per-model documentation rate rows
rows_main = ""
for m, v in data.items():
    rows_main += f"""
    <tr>
      <td class="left">{m}</td>
      <td>{v['all_doc']}/{v['all_n']}</td>
      <td><b>{v['all_pct']:.1f}</b></td>
      <td>{v['gene_doc']}/{v['gene_n']}</td>
      <td>{v['gene_pct']:.1f}</td>
      <td>{v['snp_doc']}/{v['snp_n']}</td>
      <td>{v['snp_pct']:.1f}</td>
    </tr>"""

# Overall and subgroup chi-square summary rows
rows_overall = f"""
    <tr class="stat-row">
      <td class="left" colspan="2"><i>Overall chi-square</i></td>
      <td>&chi;&sup2;({ov[2]}) = {ov[0]:.2f}</td>
      <td colspan="4">{fmt_p(ov[1])}</td>
    </tr>
    <tr class="stat-row">
      <td class="left" colspan="2"><i>Gene subgroup</i></td>
      <td>&chi;&sup2;({gn[2]}) = {gn[0]:.2f}</td>
      <td colspan="4">{fmt_p(gn[1])}</td>
    </tr>
    <tr class="stat-row">
      <td class="left" colspan="2"><i>SNP subgroup</i></td>
      <td>&chi;&sup2;({sn[2]}) = {sn[0]:.2f}</td>
      <td colspan="4">{fmt_p(sn[1])}</td>
    </tr>"""

# Pairwise comparison rows; significant pairs are bolded
rows_pairs = ""
for n1, n2, c2, p_raw, p_bonf, sig in pairs:
    cls = 'class="sig-row"' if sig else ''
    rows_pairs += f"""
    <tr {cls}>
      <td class="left">{n1} vs {n2}</td>
      <td>{c2:.2f}</td>
      <td>1</td>
      <td>{fmt_p(p_raw)}</td>
      <td>{fmt_p(p_bonf)}</td>
      <td>{"<b>*</b>" if sig else "&mdash;"}</td>
    </tr>"""

# Assemble HTML (transparent background; inherits Jupyter light/dark theme).
html = f"""<!DOCTYPE html>
<html lang="en">
<head>
<meta charset="UTF-8">
<style>
  :root {{ color-scheme: light dark; }}
  body, .table-panel, table, thead, tbody, tr, th, td {{
    background: transparent !important;
    background-color: transparent !important;
    box-shadow: none !important;
  }}
  body {{
    font-family: "Times New Roman", Times, serif;
    font-size: 12px;
    margin: 0;
    padding: 0;
    color: #E5E7EB !important;
  }}
  .table-panel {{
    margin: 0;
    padding: 0;
    border: none;
    border-radius: 0;
    max-width: none;
    color: #E5E7EB !important;
  }}
  h3 {{
    font-size: 13px;
    font-weight: bold;
    margin: 0 0 4px 0;
    color: inherit;
  }}
  .caption {{
    font-size: 11px;
    color: inherit;
    opacity: 0.92;
    margin: 0 0 8px 0;
    line-height: 1.45;
  }}
  table {{
    border-collapse: collapse;
    width: auto;
    margin: 0 0 8px 0;
  }}
  thead tr th {{
    border-top: 2px solid currentColor;
    border-bottom: 1px solid currentColor;
    padding: 4px 8px;
    text-align: center;
    font-weight: bold;
    font-size: 11.5px;
    color: inherit;
  }}
  td, th {{
    padding: 4px 8px;
    text-align: center;
    border: none;
    color: inherit;
  }}
  tbody tr:last-child td {{ border-bottom: 2px solid currentColor; }}
  .left {{ text-align: left; }}
  .section-head td {{
    background: rgba(128, 128, 128, 0.18) !important;
    color: inherit;
    font-weight: bold;
    text-align: left;
    border-top: 1px solid rgba(128, 128, 128, 0.45);
    border-bottom: 1px solid rgba(128, 128, 128, 0.45);
    padding: 4px 8px;
  }}
  .stat-row td {{
    font-style: italic;
    color: inherit;
    font-size: 11px;
    opacity: 0.95;
  }}
  .sig-row td {{
    font-weight: bold;
    color: inherit;
  }}
  .note {{
    font-size: 10.5px;
    color: inherit;
    opacity: 0.9;
    margin: 4px 0 0 0;
    line-height: 1.45;
  }}
  @media (prefers-color-scheme: dark) {{
    body {{ color: #E5E7EB !important; }}
    .table-panel {{ color: #E5E7EB !important; }}
  }}
  sup {{ font-size: 8px; }}
</style>
</head>
<body>
<div class="table-panel">

<h3>Table 3. Statistical comparison of documentation rates across four LLMs</h3>
<p class="caption">
  Documentation rate = proportion of associations with Strong or Moderate evidence support.
  Not Found associations (entities unconfirmed in Ensembl) excluded from denominators.
  Pairwise comparisons used Bonferroni correction (&alpha; = 0.05/6 &asymp; 0.0083).
  * significant after correction.
</p>

<table>
  <thead>
    <tr>
      <th class="left" rowspan="2">Model</th>
      <th colspan="2">Overall</th>
      <th colspan="2">Gene tasks</th>
      <th colspan="2">SNP tasks</th>
    </tr>
    <tr>
      <th>doc/total</th><th>Rate (%)</th>
      <th>doc/total</th><th>Rate (%)</th>
      <th>doc/total</th><th>Rate (%)</th>
    </tr>
  </thead>
  <tbody>
    {rows_main}
    <tr class="section-head"><td colspan="7">Chi-square tests (Pearson, two-sided)</td></tr>
    {rows_overall}
  </tbody>
</table>

<table>
  <thead>
    <tr>
      <th class="left">Pairwise comparison</th>
      <th>&chi;&sup2;</th><th>df</th>
      <th>Raw <i>p</i></th>
      <th>Bonferroni-corrected <i>p</i></th>
      <th>Significant</th>
    </tr>
  </thead>
  <tbody>{rows_pairs}</tbody>
</table>

<p class="note">
  doc = documented (Strong + Moderate); df = degrees of freedom.<br>
  * Significant after Bonferroni correction (threshold p &lt; 0.0083).
</p>
</div>
</body>
</html>"""

# Paper-ready exports (copy-paste into LaTeX / Word).
# Convention for the manuscript:
#   • Table 3 (main text): documentation rates ONLY (first block below).
#   • Table 4 OR Supplementary Table: pairwise chi-square (second block).
#   • Chi-square omnibus + subgroup: footnote under Table 3 OR one sentence in Results.
# The HTML preview merges all blocks for inspection; do NOT paste HTML into the paper.

rates_rows = []
for m, v in data.items():
    rates_rows.append(
        f"{LATEX_MODEL[m]} & {v['all_doc']}/{v['all_n']} & {v['all_pct']:.1f} "
        f"& {v['gene_doc']}/{v['gene_n']} & {v['gene_pct']:.1f} "
        f"& {v['snp_doc']}/{v['snp_n']} & {v['snp_pct']:.1f} \\\\"
    )
rates_body = "\n".join(rates_rows)

pair_rows = []
for n1, n2, c2, p_raw, p_bonf, sig in pairs:
    mark = "*" if sig else "---"
    pair_rows.append(
        f"{n1} vs {n2} & {c2:.2f} & 1 & {fmt_p_latex(p_raw)} & {fmt_p_latex(p_bonf)} & {mark} \\\\"
    )
pair_body = "\n".join(pair_rows)

latex_rates = f"""% ========== TABLE 3 (main text): documentation rates ==========
% Requires: \\usepackage{{booktabs}} \\usepackage{{threeparttable}}
\\begin{{table}}[ht]
\\centering
\\caption{{Documentation rates across four LLMs by task type.}}
\\label{{tab:doc_rates}}
\\begin{{threeparttable}}
\\begin{{tabular}}{{lcccccc}}
\\toprule
 & \\multicolumn{{2}}{{c}}{{Overall}} & \\multicolumn{{2}}{{c}}{{Gene tasks}} & \\multicolumn{{2}}{{c}}{{SNP tasks}} \\\\
Model & doc/total & Rate (\\%) & doc/total & Rate (\\%) & doc/total & Rate (\\%) \\\\
\\midrule
{rates_body}
\\bottomrule
\\end{{tabular}}
\\begin{{tablenotes}}[flushleft]\\footnotesize
Documentation rate = proportion of associations with Strong or Moderate evidence.
Not Found associations excluded from denominators.
Overall $\\chi^2({ov[2]})={ov[0]:.2f}$, $p={fmt_p_latex(ov[1])}$;
gene subgroup $\\chi^2({gn[2]})={gn[0]:.2f}$, $p={fmt_p_latex(gn[1])}$;
SNP subgroup $\\chi^2({sn[2]})={sn[0]:.2f}$, $p={fmt_p_latex(sn[1])}$ (Pearson, two-sided).
\\end{{tablenotes}}
\\end{{threeparttable}}
\\end{{table}}
"""

latex_pairwise = f"""% ========== TABLE 4 (or Supplementary): pairwise comparisons ==========
\\begin{{table}}[ht]
\\centering
\\caption{{Pairwise chi-square tests of documentation rates (Bonferroni $\\alpha=0.05/6\\approx0.0083$).}}
\\label{{tab:doc_pairwise}}
\\begin{{threeparttable}}
\\begin{{tabular}}{{lccccc}}
\\toprule
Comparison & $\\chi^2$ & df & Raw $p$ & Bonferroni $p$ & Sig. \\\\
\\midrule
{pair_body}
\\bottomrule
\\end{{tabular}}
\\begin{{tablenotes}}[flushleft]\\footnotesize
Sig.: * = significant after Bonferroni correction ($p<0.0083$).
\\end{{tablenotes}}
\\end{{threeparttable}}
\\end{{table}}
"""

latex_all = latex_rates + "\n\n" + latex_pairwise

# TSV for Word: Insert > Table > Convert Text to Table (tab separator)
tsv_lines = [
    "TABLE 3\tdoc rates\t(paste as main-text table)",
    "Model\tOverall doc/total\tOverall Rate (%)\tGene doc/total\tGene Rate (%)\tSNP doc/total\tSNP Rate (%)",
]
for m, v in data.items():
    tsv_lines.append(
        f"{m}\t{v['all_doc']}/{v['all_n']}\t{v['all_pct']:.1f}\t"
        f"{v['gene_doc']}/{v['gene_n']}\t{v['gene_pct']:.1f}\t"
        f"{v['snp_doc']}/{v['snp_n']}\t{v['snp_pct']:.1f}"
    )
tsv_lines += [
    "TABLE 4\tpairwise chi-square\t(separate table number in manuscript)",
    "Comparison\tchi2\tdf\tRaw p\tBonferroni p\tSignificant",
]
for n1, n2, c2, p_raw, p_bonf, sig in pairs:
    tsv_lines.append(
        f"{n1} vs {n2}\t{c2:.2f}\t1\t{fmt_p_plain(p_raw)}\t{fmt_p_plain(p_bonf)}\t{'*' if sig else ''}"
    )
tsv_text = "\n".join(tsv_lines)

from pathlib import Path
out_dir = PROJECT_DIR if "PROJECT_DIR" in globals() else Path(".")
out_dir = Path(out_dir)
paths = {
    "html": out_dir / "stat_table.html",
    "latex": out_dir / "stat_table_for_paper.tex",
    "tsv": out_dir / "stat_table_for_word.tsv",
}
paths["html"].write_text(html, encoding="utf-8")
paths["latex"].write_text(latex_all, encoding="utf-8")
paths["tsv"].write_text(tsv_text, encoding="utf-8")

display(HTML(html))
print("=" * 72)
print("PAPER TABLE COUNT (recommended):")
print("  • 2 numbered tables: Table 3 = rates | Table 4 = pairwise chi-square")
print("  • Omnibus chi-square (overall/gene/SNP): footnote under Table 3, not a third table")
print("  • Do NOT paste HTML into Word/LaTeX — use .tex or .tsv below")
print("=" * 72)
for k, p in paths.items():
    print(f"  Saved [{k}]: {p}")
if VERBOSE:
    print("LaTeX/TSV exports saved to stat_table_for_paper.tex and stat_table_for_word.tsv")


# %% [code]
from docx import Document
from docx.shared import Pt, Cm, RGBColor
from docx.enum.text import WD_ALIGN_PARAGRAPH
from docx.oxml.ns import qn
from docx.oxml import OxmlElement
import numpy as np

if "data" not in globals() or "ov" not in globals() or "pairs" not in globals():
    raise RuntimeError("Run the chi-square / stat table cell above first.")


def _word_p(p: float) -> str:
    if p < 0.001:
        exp = int(np.floor(np.log10(p)))
        coef = p / 10**exp
        superscript = str(exp).translate(str.maketrans("0123456789-", "⁰¹²³⁴⁵⁶⁷⁸⁹⁻"))
        return f"{coef:.2f} × 10{superscript}"
    return f"{p:.4f}"

def set_cell_border(cell, top=None, bottom=None, left=None, right=None):
    """Set borders for a single cell."""
    tc = cell._tc
    tcPr = tc.get_or_add_tcPr()
    tcBorders = OxmlElement('w:tcBorders')
    for side, val in [('top', top), ('bottom', bottom), ('left', left), ('right', right)]:
        if val:
            el = OxmlElement(f'w:{side}')
            el.set(qn('w:val'), val.get('val', 'single'))
            el.set(qn('w:sz'), str(val.get('sz', 4)))
            el.set(qn('w:space'), '0')
            el.set(qn('w:color'), val.get('color', '000000'))
            tcBorders.append(el)
    tcPr.append(tcBorders)

def set_cell_text(cell, text, bold=False, italic=False, align='left', size=10):
    cell.text = ''
    para = cell.paragraphs[0]
    para.alignment = WD_ALIGN_PARAGRAPH.CENTER if align == 'center' else WD_ALIGN_PARAGRAPH.LEFT
    run = para.add_run(text)
    run.bold = bold
    run.italic = italic
    run.font.size = Pt(size)
    run.font.name = 'Times New Roman'

def merge_and_set(table, row, col_start, col_end, text, bold=False, italic=False, align='center', size=10):
    cell = table.cell(row, col_start)
    if col_end > col_start:
        cell = cell.merge(table.cell(row, col_end))
    set_cell_text(cell, text, bold=bold, italic=italic, align=align, size=size)
    return cell

doc = Document()

# Set narrow margins
for section in doc.sections:
    section.left_margin = Cm(2)
    section.right_margin = Cm(2)

# Title
title_para = doc.add_paragraph()
title_run = title_para.add_run(
    "Table 3. Chi-square tests and pairwise comparisons of documentation rates across four LLMs."
)
title_run.bold = True
title_run.font.size = Pt(10)
title_run.font.name = 'Times New Roman'

# Table 1: documentation rates (9 rows x 7 cols)
t1 = doc.add_table(rows=9, cols=7)
t1.allow_autofit = False

# Set column widths
widths = [Cm(3.8), Cm(2.0), Cm(1.6), Cm(2.0), Cm(1.6), Cm(2.0), Cm(1.6)]
for row in t1.rows:
    for i, cell in enumerate(row.cells):
        cell.width = widths[i]

THICK = {'val': 'single', 'sz': 12, 'color': '000000'}
THIN  = {'val': 'single', 'sz': 4,  'color': '000000'}
NONE  = {'val': 'none',   'sz': 0,  'color': 'FFFFFF'}

# Row 0: top border + spanning headers
merge_and_set(t1, 0, 0, 0, "Model", bold=True, align='left')
merge_and_set(t1, 0, 1, 2, "Overall", bold=True)
merge_and_set(t1, 0, 3, 4, "Gene tasks", bold=True)
merge_and_set(t1, 0, 5, 6, "SNP tasks", bold=True)

for c in range(7):
    cell = t1.cell(0, c)
    set_cell_border(cell, top=THICK, bottom=THIN, left=NONE, right=NONE)

# Row 1: sub-headers
sub = ["", "doc/total", "Rate (%)", "doc/total", "Rate (%)", "doc/total", "Rate (%)"]
for c, h in enumerate(sub):
    set_cell_text(t1.cell(1, c), h, bold=True, align='center' if c > 0 else 'left')
    set_cell_border(t1.cell(1, c), bottom=THIN, top=NONE, left=NONE, right=NONE)

# Data rows 2-5 (from dynamic chi-square cell)
data_rows = []
for _m, _v in data.items():
    data_rows.append([
        _m,
        f"{_v['all_doc']}/{_v['all_n']}",
        f"{_v['all_pct']:.1f}",
        f"{_v['gene_doc']}/{_v['gene_n']}",
        f"{_v['gene_pct']:.1f}",
        f"{_v['snp_doc']}/{_v['snp_n']}",
        f"{_v['snp_pct']:.1f}",
    ])
for r, row_data in enumerate(data_rows):
    for c, val in enumerate(row_data):
        align = 'left' if c == 0 else 'center'
        set_cell_text(t1.cell(r+2, c), val, align=align)
        set_cell_border(t1.cell(r+2, c), top=NONE, bottom=THIN, left=NONE, right=NONE)

# Row 6: section header (Chi-square label, spans all)
chi_cell = merge_and_set(t1, 6, 0, 6, "Chi-square tests (Pearson, two-sided)", bold=True, align='left')
set_cell_border(chi_cell, top=THIN, bottom=THIN, left=NONE, right=NONE)

# Rows 7-8: chi-square results, each spans all 7 cols
chi_data = [
    ("Overall",       f"χ²({ov[2]}) = {ov[0]:.2f}", f"p = {_word_p(ov[1])}"),
    ("Gene subgroup", f"χ²({gn[2]}) = {gn[0]:.2f}", f"p = {_word_p(gn[1])}"),
    ("SNP subgroup",  f"χ²({sn[2]}) = {sn[0]:.2f}", f"p = {_word_p(sn[1])}"),
]
for r, (label, stat, pval) in enumerate(chi_data):
    text = f"{label}: {stat},  {pval}"
    is_last = (r == len(chi_data) - 1)
    bottom_border = THICK if is_last else THIN
    cell = merge_and_set(t1, r+6, 0, 6, text, italic=True, align='left')
    set_cell_border(cell, top=NONE, bottom=bottom_border, left=NONE, right=NONE)

# Rebuild rows 7-8 properly (row 6 already used, shift by 1)
for r, (label, stat, pval) in enumerate(chi_data):
    text = f"{label}: {stat},  {pval}"
    is_last = (r == 2)
    bottom_border = THICK if is_last else THIN
    cell = t1.cell(r+6, 0)
    try:
        cell = cell.merge(t1.cell(r+6, 6))
    except Exception:
        pass
    set_cell_text(cell, text, italic=True, align='left')
    set_cell_border(cell, top=NONE, bottom=bottom_border, left=NONE, right=NONE)

doc.add_paragraph()

# Table 2: pairwise comparisons (7 rows x 6 cols)
t2 = doc.add_table(rows=7, cols=6)
t2.allow_autofit = False

widths2 = [Cm(5.0), Cm(1.4), Cm(1.0), Cm(2.2), Cm(2.8), Cm(1.8)]
for row in t2.rows:
    for i, cell in enumerate(row.cells):
        cell.width = widths2[i]

# Header
headers2 = ["Pairwise comparison", "χ²", "df", "Raw p", "Bonferroni-corrected p", "Significant"]
for c, h in enumerate(headers2):
    set_cell_text(t2.cell(0, c), h, bold=True, align='center' if c > 0 else 'left')
    set_cell_border(t2.cell(0, c), top=THICK, bottom=THIN, left=NONE, right=NONE)

# Data (pairwise comparisons from dynamic chi-square cell)
pair_rows = []
for n1, n2, c2, p_raw, p_bonf, sig in pairs:
    pair_rows.append([
        f"{n1} vs {n2}",
        f"{c2:.2f}",
        "1",
        _word_p(p_raw),
        _word_p(p_bonf),
        "*" if sig else "—",
    ])
for r, row_data in enumerate(pair_rows):
    is_last = (r == len(pair_rows) - 1)
    sig = row_data[5] == "*"
    bottom_border = THICK if is_last else THIN
    for c, val in enumerate(row_data):
        align = 'left' if c == 0 else 'center'
        set_cell_text(t2.cell(r+1, c), val, bold=sig, align=align)
        set_cell_border(t2.cell(r+1, c), top=NONE, bottom=bottom_border, left=NONE, right=NONE)

# Footnote
note = doc.add_paragraph()
note_run = note.add_run(
    "doc = documented (Strong + Moderate); df = degrees of freedom.\n"
    "* Significant after Bonferroni correction (threshold p < 0.0083)."
)
note_run.font.size = Pt(9)
note_run.font.name = 'Times New Roman'

doc.save("Table3_stats.docx")
print("Table3_stats.docx saved.")
```
