## Supplementary material for "Assessing the Reliability of LLM-Generated Phenotype-Genotype Associations Through External Validation": nvenn_render_panel.R

```
# Renders one nVennR panel from a JSON entity-set file. Called by
# generate_euler.py via subprocess, once per panel.
#
# Usage: Rscript nvenn_render_panel.R <input_json> <out_svg> <out_counts_json> <n_cycles>
# input_json = {"model-1": ["key1","key2",...], ...}, keys in MODEL_ORDER
# (order matters -- it drives the color mapping).
suppressMessages({
  library(nVennR)
  library(jsonlite)
  library(rsvg)
})

args <- commandArgs(trailingOnly = TRUE)
in_json     <- args[1]
out_svg     <- args[2]
out_counts  <- args[3]
n_cycles    <- as.integer(args[4])
out_png     <- sub("\\.svg$", ".png", out_svg)

sets <- fromJSON(in_json, simplifyVector = FALSE)
sets <- lapply(sets, function(x) as.character(unlist(x)))
# nVennR requires non-empty sets.
sets <- sets[sapply(sets, length) > 0]

if (length(sets) == 0) {
  writeLines("{}", out_counts)
  svg(out_svg, width = 4, height = 4)
  plot.new()
  dev.off()
  quit(status = 0)
}

MODEL_COLOR_MAP <- c(
  "gpt-5.5"                 = "#F48FB1",
  "gemini-3-flash-preview"  = "#A5D6A7",
  "claude-sonnet-4-6"       = "#FFF59D",
  "deepseek-v4-flash"       = "#90CAF9"
)
set_colors <- unname(MODEL_COLOR_MAP[names(sets)])

myV <- plotVenn(sets, nCycles = n_cycles, showPlot = FALSE)

showSVG(
  myV,
  opacity     = 0.5,
  borderWidth = 1.5,
  showLegend  = FALSE,
  outFile     = out_svg,
  setColors   = set_colors,
  labelRegions = FALSE,
  showNumbers  = TRUE,
  fontScale    = 1.9
)

# nVennR hardcodes font-family: Arial in the SVG it writes -- swap to
# Times New Roman for consistency with every other figure in the paper.
svg_text <- readLines(out_svg, warn = FALSE)
svg_text <- gsub("font-family: Arial;", "font-family: 'Times New Roman', Times, serif;", svg_text, fixed = TRUE)
writeLines(svg_text, out_svg)

rsvg_png(out_svg, out_png, width = 1400, height = 1000)

regs <- listVennRegions(myV, na.rm = TRUE)
counts <- sapply(regs, length)
# nVennR region names look like "1, 0, 1 (Name1, Name3)" -- pull out the
# parenthesized model list, which is what we key the JSON export on.
region_names <- sub(".*\\((.*)\\)", "\\1", names(counts))
out_list <- setNames(as.list(as.integer(counts)), region_names)
write(toJSON(out_list, auto_unbox = TRUE), out_counts)

cat("Rendered:", out_svg, " regions:", length(counts), "\n")
```
