## Supplementary figures and images for "Assessing the Reliability of LLM-Generated Phenotype-Genotype Associations Through External Validation"

### combined_figure.png

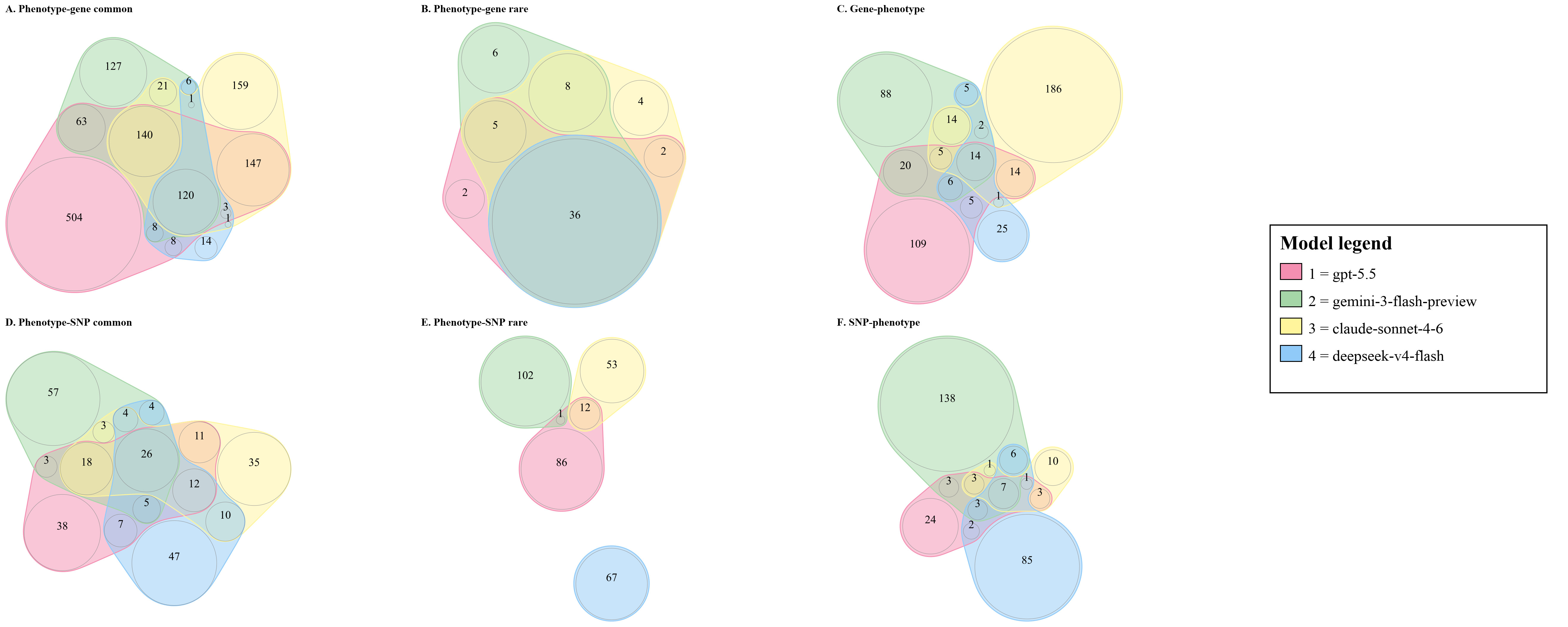

### panel_A.png

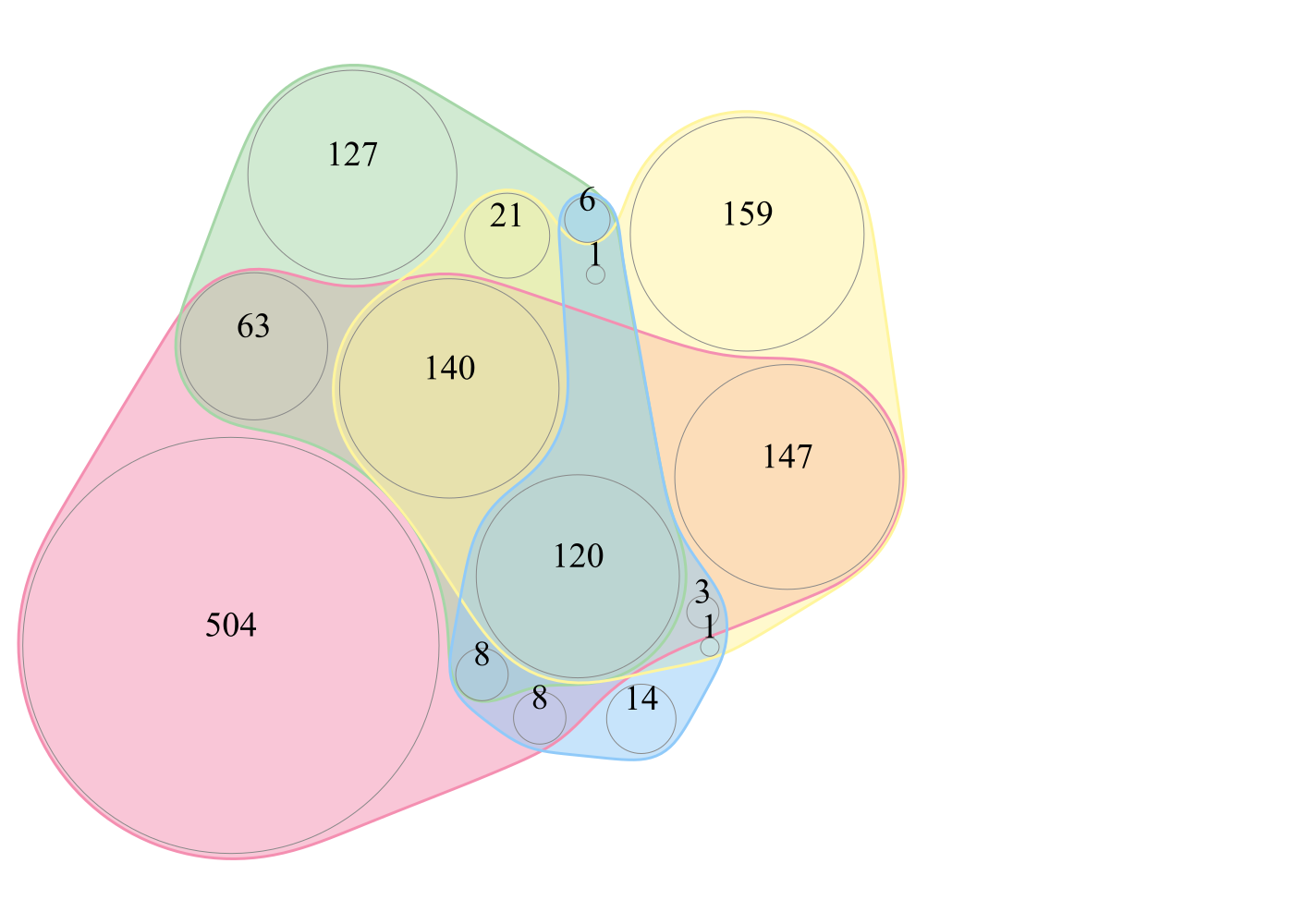

### panel_B.png

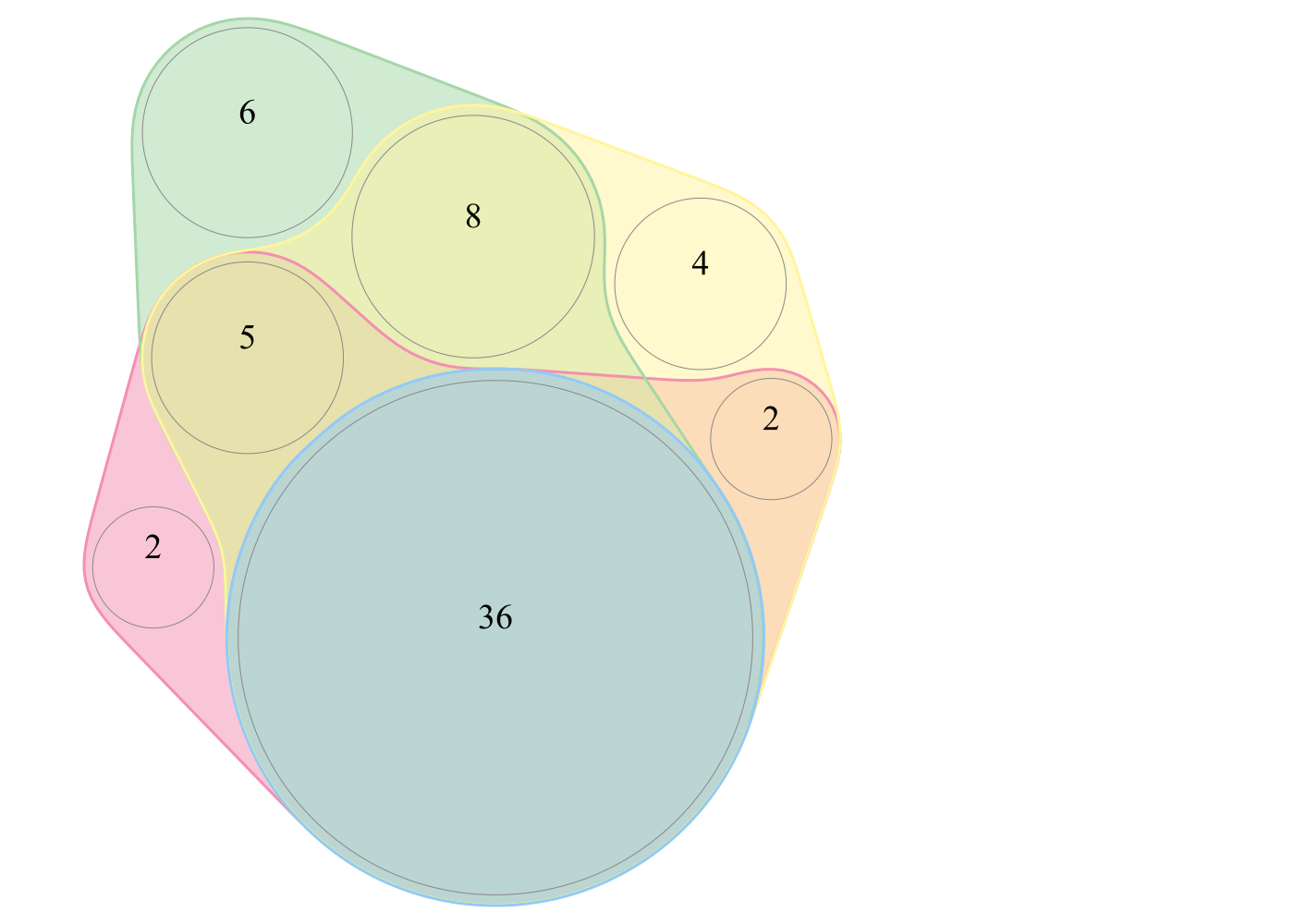

### panel_C.png

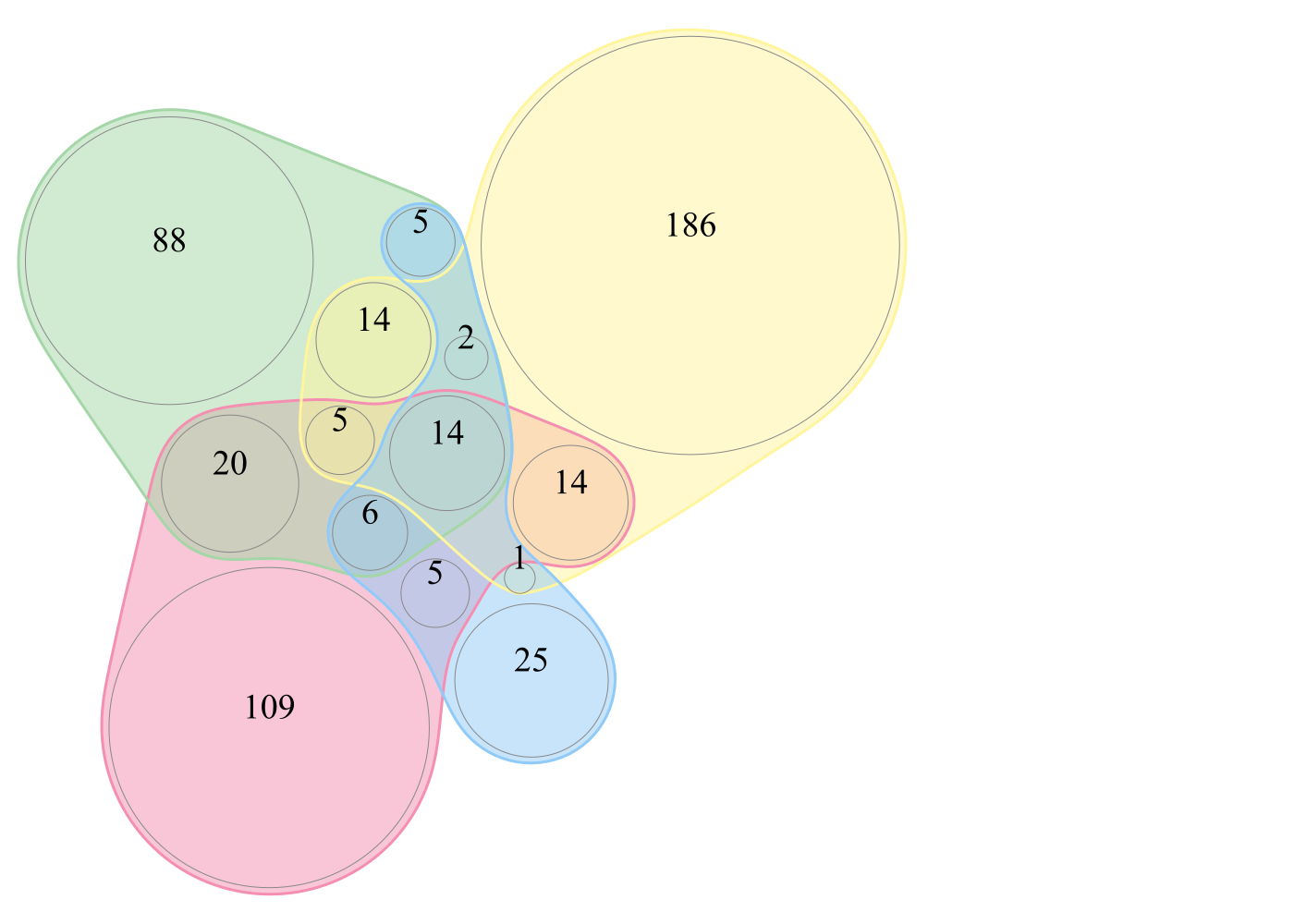

### panel_D.png

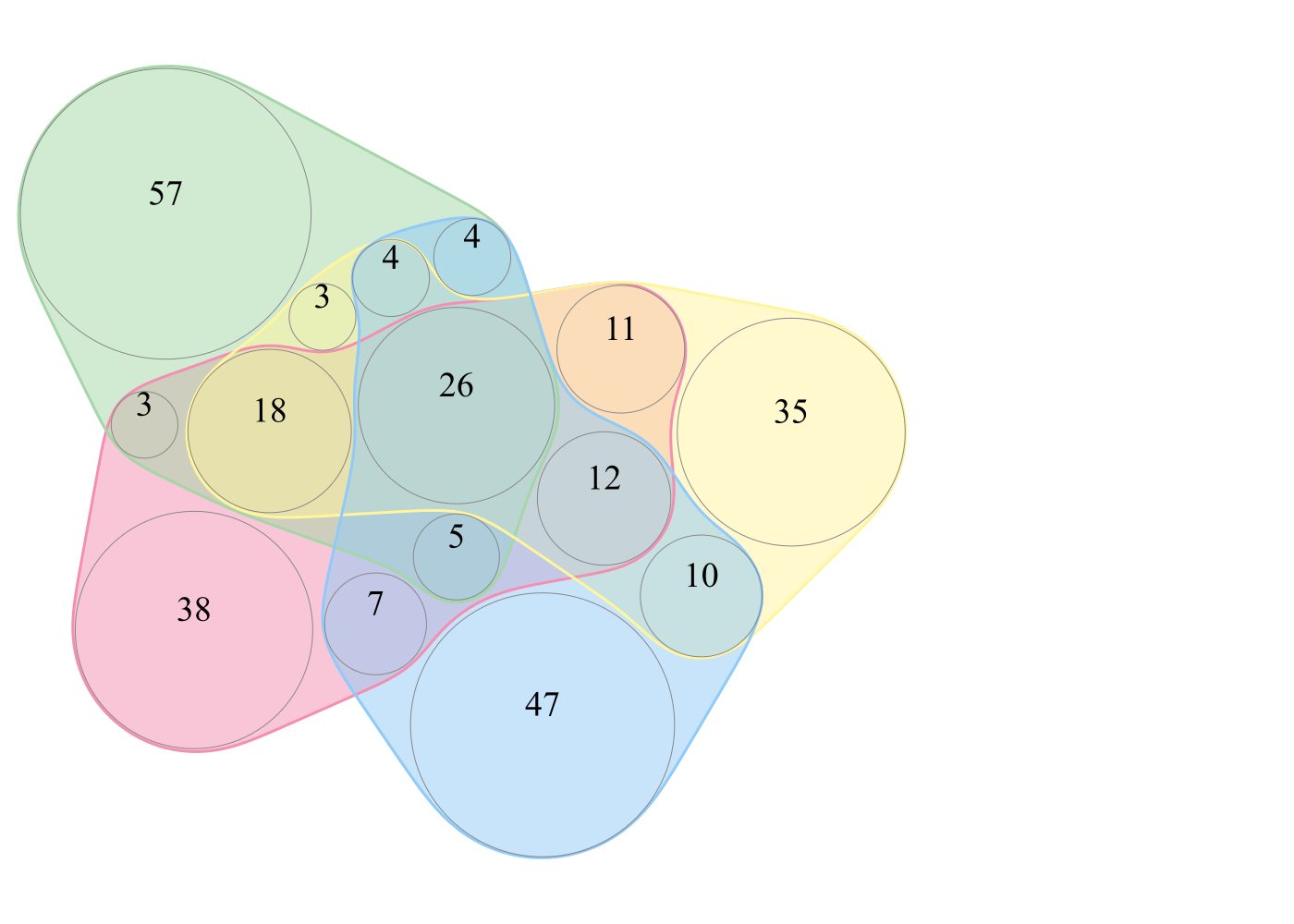

### panel_E.png

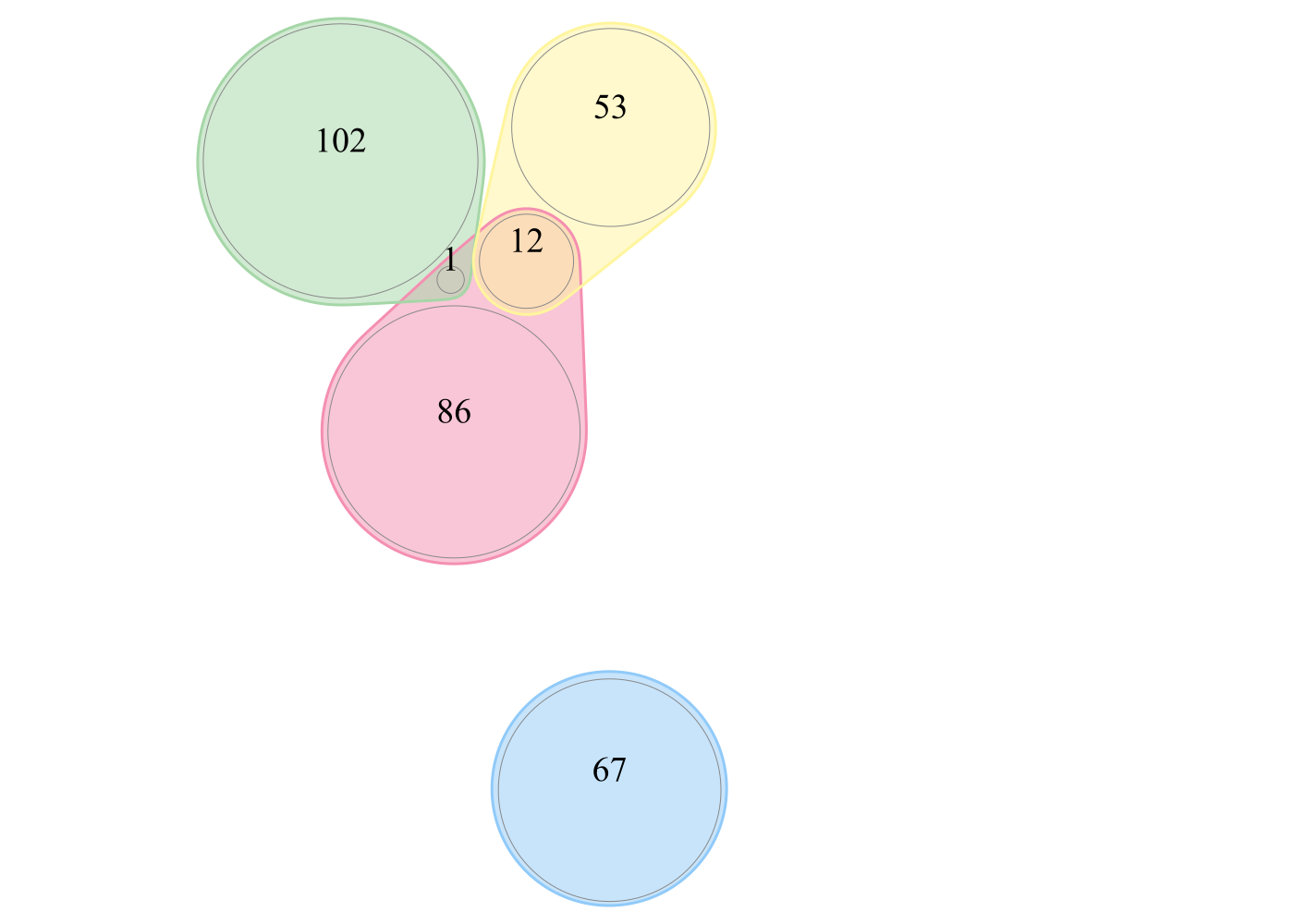

### panel_F.png

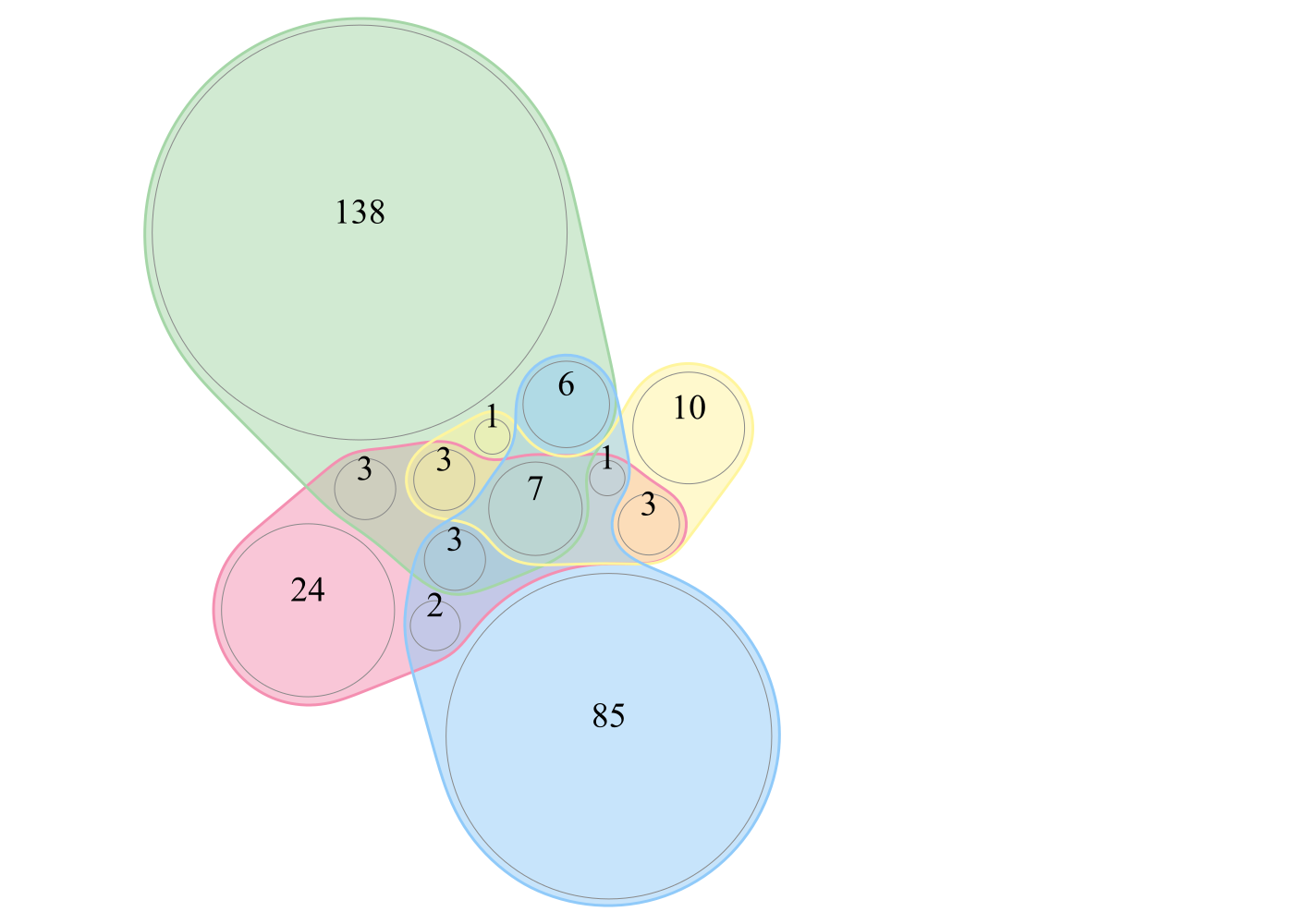

### Supplementary Figure S1. Task-level matched rates by model across six task categories.png

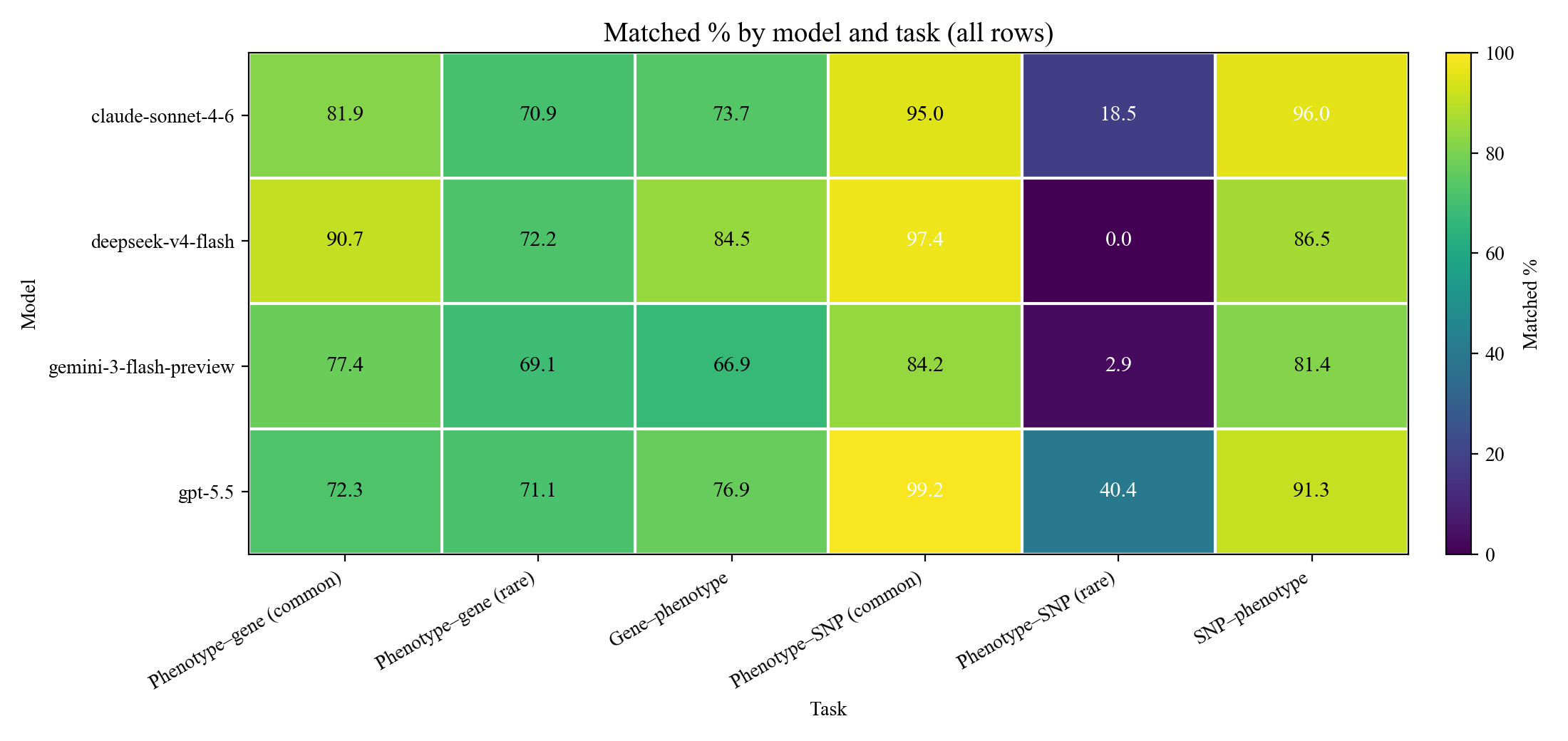

### Supplementary Figure S2. GWAS and OMIM evidence source contributions by task category.png

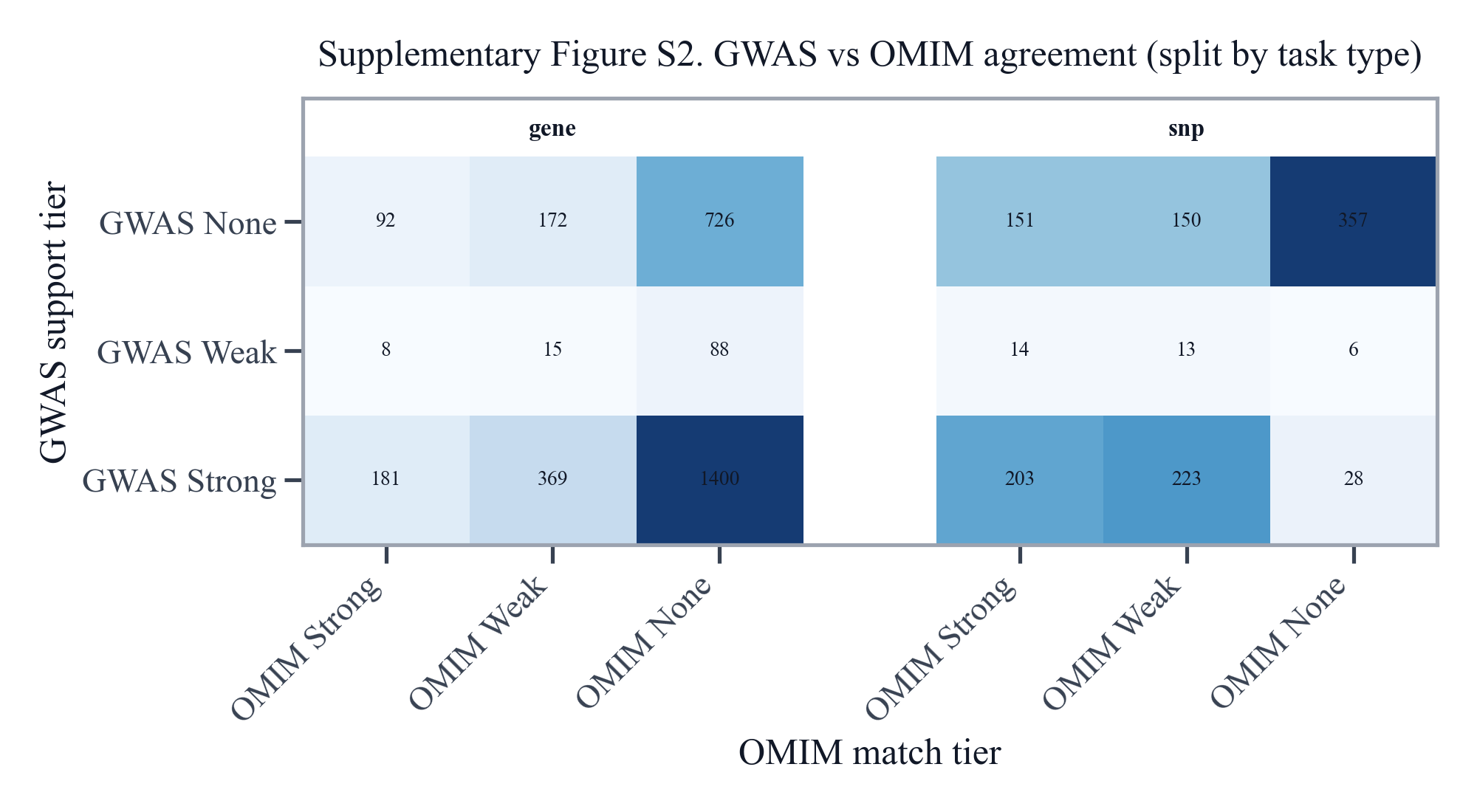

### Supplementary Table S1. Gene-task Ensembl identifier validation summary by model.jpg

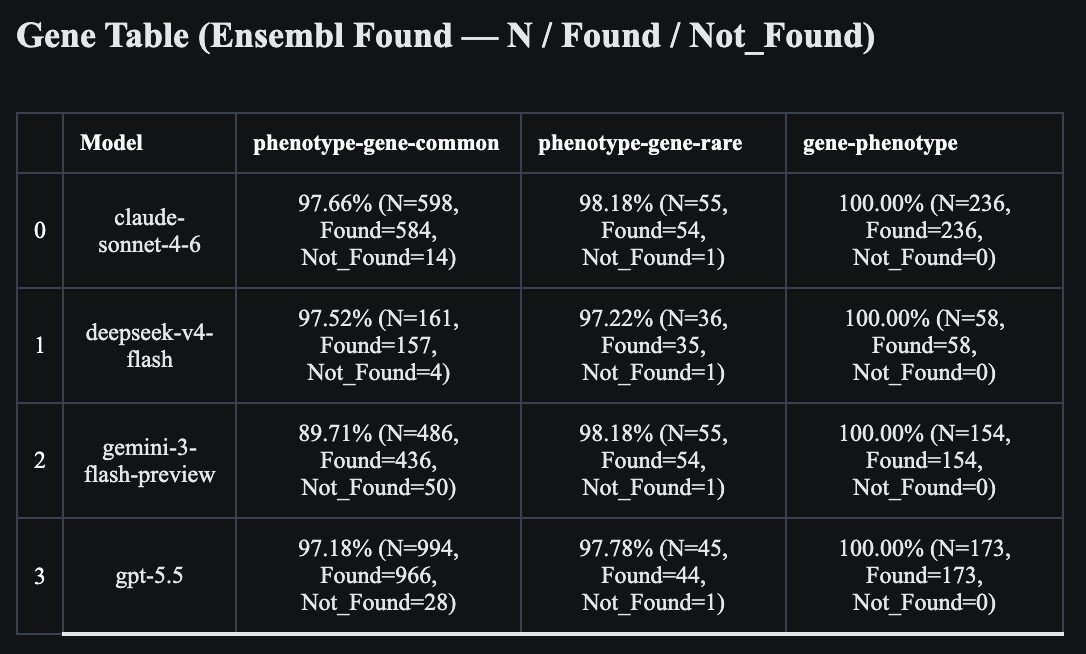

### Supplementary Table S2. SNP-task Ensembl identifier validation summary by model.jpg

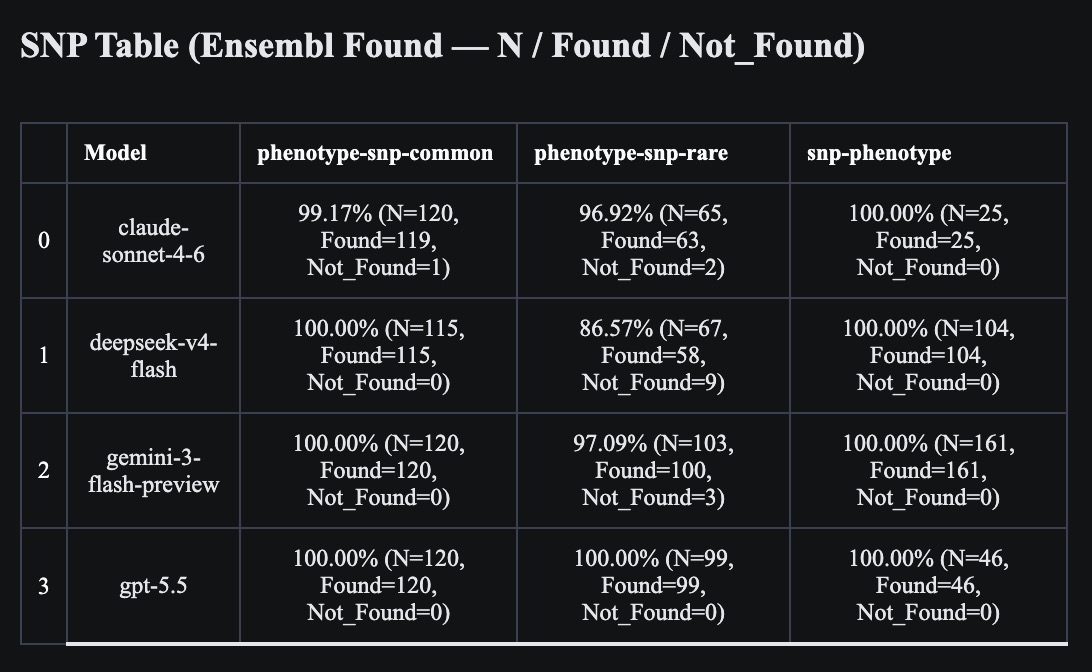

### Supplementary Table S3. Gene-task evidence distribution by model.jpg

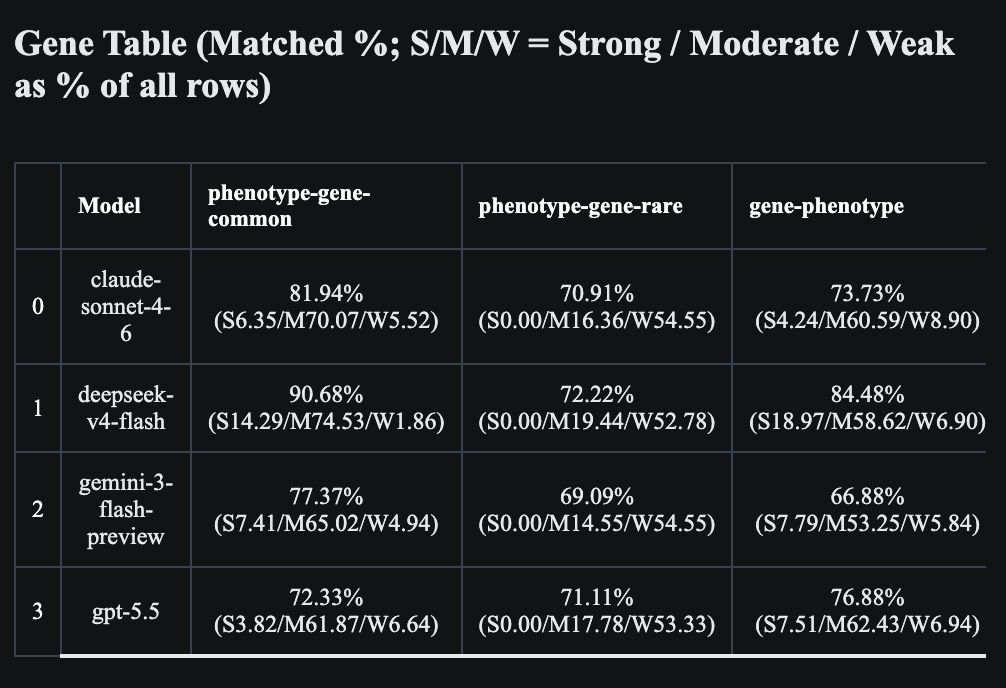

### Supplementary Table S4. SNP-task evidence distribution by model.jpg

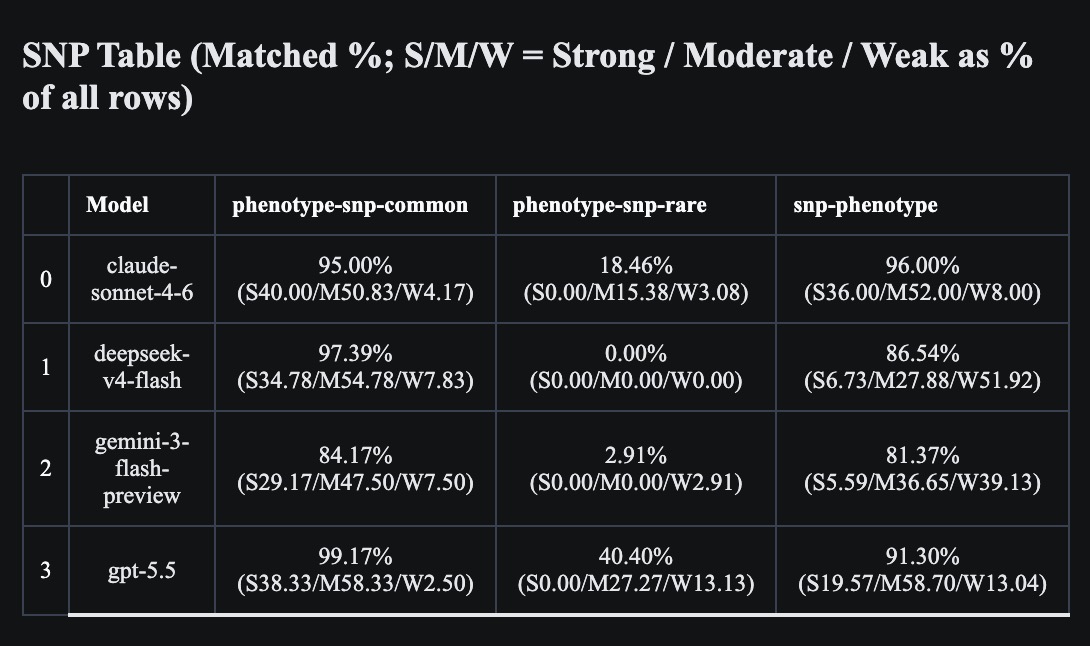
